# Probing immune signatures of conjugated pattern recognition receptor ligands identifies chimeras with adjuvant and antitumor activity

**DOI:** 10.1101/2025.04.07.647694

**Authors:** Špela Janež, Samo Guzelj, Marcela Šišić, Ruža Frkanec, Stane Pajk, Lenny Burgmeijer, Bram Slütter, Žiga Jakopin

## Abstract

Pattern recognition receptor (PRR) ligands hold great promise as adjuvants and immunotherapeutics. Here, we demonstrate that chemical conjugation of PRR ligands results in synergistic immune response amplification inaccessible to unlinked agonist mixtures. To identify potent immune agonists, we synthesized conjugated PRR ligands incorporating distinct agonist pairings, each targeting two carefully selected PRRs. We used a phenotypic screen using human peripheral blood mononuclear cells (PBMCs) to single out chimeric PRR ligands capable of inducing robust immune response both in terms of cytokine response and cytotoxicity against cancer cells. Chimeric TLR4/TLR7 and TLR7/RIG-I ligands showed broad immune activation *in vitro* as well as enhancement of antigen-specific cellular and humoral responses in mice. Intratumoral delivery of chimeric TLR4/TLR7 ligand induced robust antitumor response in a syngeneic mouse B16F10 tumor model. These results demonstrate the profound effects that conjugation can have on immune response and support the use of conjugated PRR ligands as adjuvants/immunotherapeutics.

**ONE SENTENCE SUMMARY:** Conjugated PRR ligands enhance immune activation, serve as potent adjuvants, and induce antitumor response in a mouse model.

**GRAPHICAL ABSTRACT:** 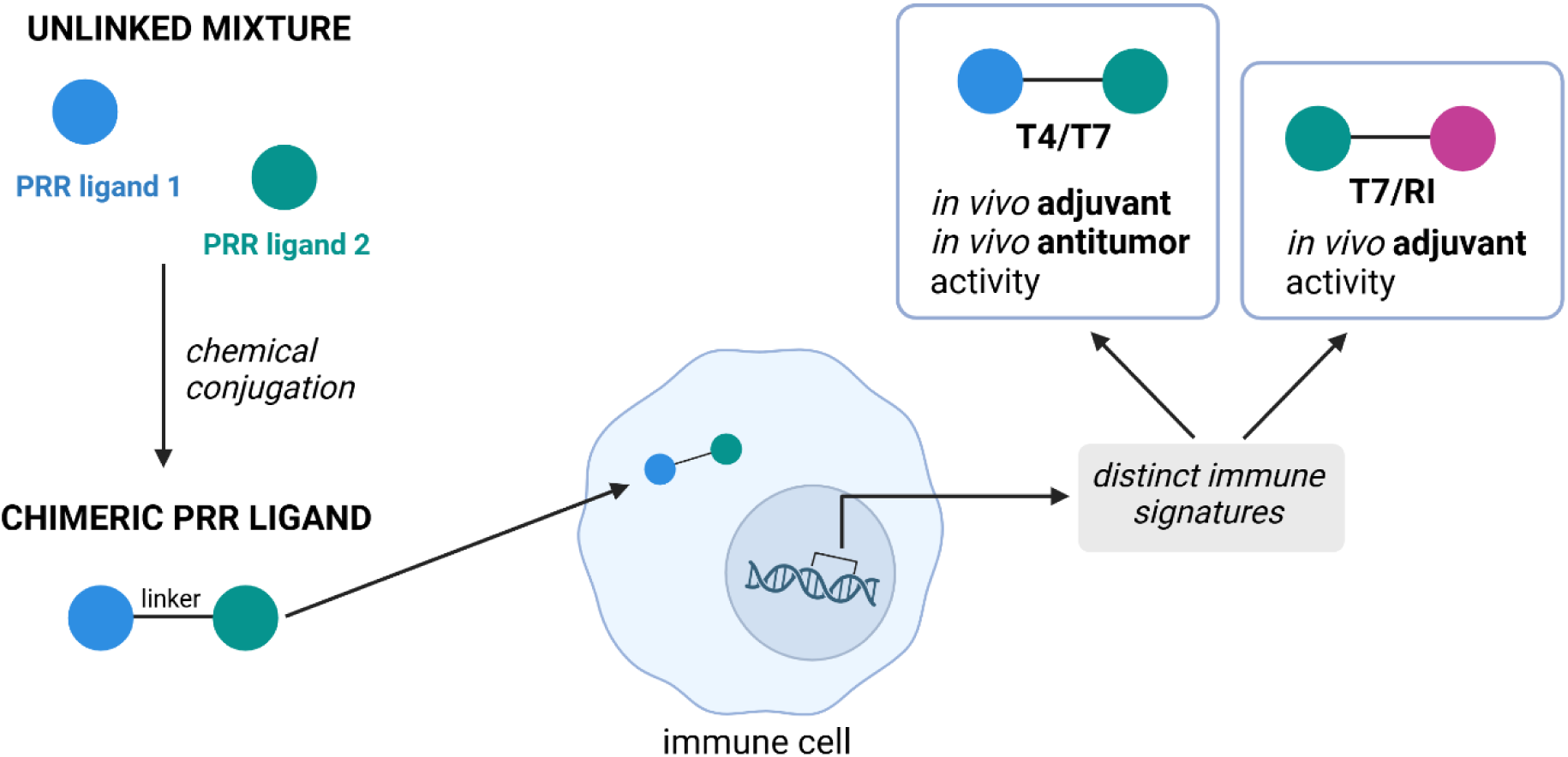

## 1 Introduction

Stimulation of the innate immune system is crucial in both effective vaccinations and immunotherapies. This is often achieved through adjuvants or immunotherapeutics, molecules that resemble pathogen-associated molecular patterns (PAMPs) and activate pattern recognition receptors (PRRs), such as Toll-like receptors (TLRs), nucleotide-binding oligomerization domain (NOD)-like receptors (NLRs), and retinoic acid-inducible gene-1-like receptors (RLRs), present in immune cells, in particular dendritic cells (DCs) (*1*). On activation, these receptors engage two innate immune signaling pathways: the nuclear factor kappa-light-chain-enhancer of activated B-cells pathway (NF-κB) and the interferon regulatory factors pathway (IRF) thus providing indispensable initial signals that determine the type, the magnitude, and the durability of adaptive immune response (*2*).

Synthetic PRR ligands are promising candidates for adjuvant and immunotherapeutic development. However, they do not always elicit robust immune responses, while excessive activation can also trigger systemic inflammation. Additionally, repeated stimulation with these agonists frequently leads to immune tolerance (*3*). During natural infections, pathogens containing multiple pathogen-associated molecular patterns (PAMPs) engage several innate immune receptors simultaneously. For instance, the yellow fever virus vaccine induces a systemic immune response through the synergistic activation of multiple PRRs, including TLRs 2, 7, 8, and 9 (*4*). This principle is also applied in clinically approved vaccine adjuvants such as AS01 and AS04 (*5*). Co-activation of distinct innate immune receptors by agonist mixtures enhances immune signaling through cross-talk and cross-regulation, amplifying immune responses (*6, 7*). Notably, chemical conjugation of such agonists enables simultaneous activation of specific targets within the same cell, potentially yielding superior immune responses while also affecting the pharmacokinetic properties (e.g. reduced systemic distribution) (*8*). Several conjugated dual TLR ligands with immunoenhancing properties have been reported, including TLR2/6/TLR9 (*9*), TLR4/9 (*10*), and TLR2/6/TLR7 (*11*). Among NLR/TLR combinations, chimeric ligands targeting NOD2/TLR2/6 (*12*), NOD2/TLR7 (*13–15*), and NOD2/TLR4 (*16*) have demonstrated synergistic *in vivo* adjuvant activity, irrespective of the localization of the PRRs. This includes the conjugation of ligands targeting PRRs in different cellular compartments, further validating the proof-of-concept for covalent linkage.

Understanding the complexity of immune cross-talk is essential for the rational design of adjuvants and immunotherapeutics, as an effective immune response requires precise coordination. Encouraged by the promising results of our conjugated NOD2/TLR7 ligands (*13, 14*), we aim to dissect the synergistic interactions within the innate immune system and identify those capable of eliciting potent immune responses. In this study, we synthesized conjugated PRR ligands incorporating various agonist combinations, each targeting two of the following PRRs: TLR2, TLR4 (both extracellular), NOD2, TLR7, and RIG-I (all intracellular). These receptors were selected based on their well-established immunostimulatory properties (*17–27*), their ability to enhance the non-specific antitumor activity of immune effector cells within the tumor microenvironment (*28–32*), and their role in promoting autophagy (*33–38*), which facilitates cross-presentation and strengthens cellular responses. Additionally, synergistic effects—both in terms of cytokine secretion (*39–49*) and adjuvant activity (*50–73*)—have been reported for all possible pairwise combinations of these receptors. To identify the most effective conjugates, we employed a two-step *in vitro* screening approach. First, we evaluated immune activation by measuring the magnitude and the breadth of cytokine responses in human peripheral blood mononuclear cells (PBMCs). Next, we assessed the ability of these conjugates to induce PBMC-mediated cytotoxicity against cancer cells. The top-performing candidates were further tested for *ex vivo* adjuvant activity in murine bone marrow-derived dendritic cells (BMDCs). Finally, we identified potent chimeric combinations, conjugated TLR4/TLR7 and TLR7/RIG-I ligands that functioned as effective *in vivo* vaccine adjuvants. Notably, the TLR4/TLR7 ligand also induced robust antitumor response in a mouse B16F10 tumor model.

## 2 Results

### Design and construction of conjugated PRR ligands

The signaling architecture of PRRs is multipronged, therefore single molecules which activate multiple PRRs can provide distinct responses. To take advantage of the synergistic cross-talk, we probed the potential of *bona fide* PRR ligands covalently linked into chimeric conjugates to manipulate their signaling activity. Specifically, the designed chimeras feature unique pairwise combinations of the following small molecule ligands: **N2** (NOD2 agonist), **T1/2** (TLR1/2 agonist), **T4** (TLR4 agonist), **T7** (TLR7 agonist) and **RI** (RIG-I agonist) (Fig. 1). These agonists were chosen for their structural simplicity and synthetic accessibility. For NOD2 activation, we used **N2**, a highly potent agonist previously identified by our group (*74*). While TLR2/6 agonists have been conjugated with NOD2 agonists in prior studies (*12, 75*), they are associated with systemic inflammation (*76*). Therefore, to target TLR2, which signals as a heterodimer with either TLR1 or TLR6, we opted for the small-molecule TLR1/2 agonist **T1/2**, which possesses notable adjuvant properties (*31, 77*). For TLR4 activation, we employed the carboxylic acid precursor **T4** of the pyrimido[5,4-b]indole agonist **T4’** (*78*). Although **T4** is only a weak agonist *per se*, its chemical properties make it well-suited for conjugation as shown previously (*10*). Similarly, the purine-based TLR7 agonist **T7** was chosen based on previous reports demonstrating its suitability for conjugation (*79, 80*). For RIG-I activation, we selected **RI**, the active form of the RIG-I agonist **RI’**, as the most viable option (*81*). All selected agonists feature functional groups that enable conjugation. In our design, we linked these agonists using either 6-aminohexanoic acid or poly(2-aminoethyl)ether spacers through efficient coupling chemistry (Schemes S1–S3), generating a panel of nine conjugates: **T4/N2**, **N2/RI**, **T1/2/N2**, **T1/2/T4**, **T1/2/T7**, **T1/2/RI**, **T4/T7**, **T4/RI**, and **T7/RI** (Fig. 1; full structures are shown in Fig. S1). Notably, this panel excludes the previously reported NOD2/TLR7 combination featured in **N2/T7** (*13*).

**Fig. 1.**
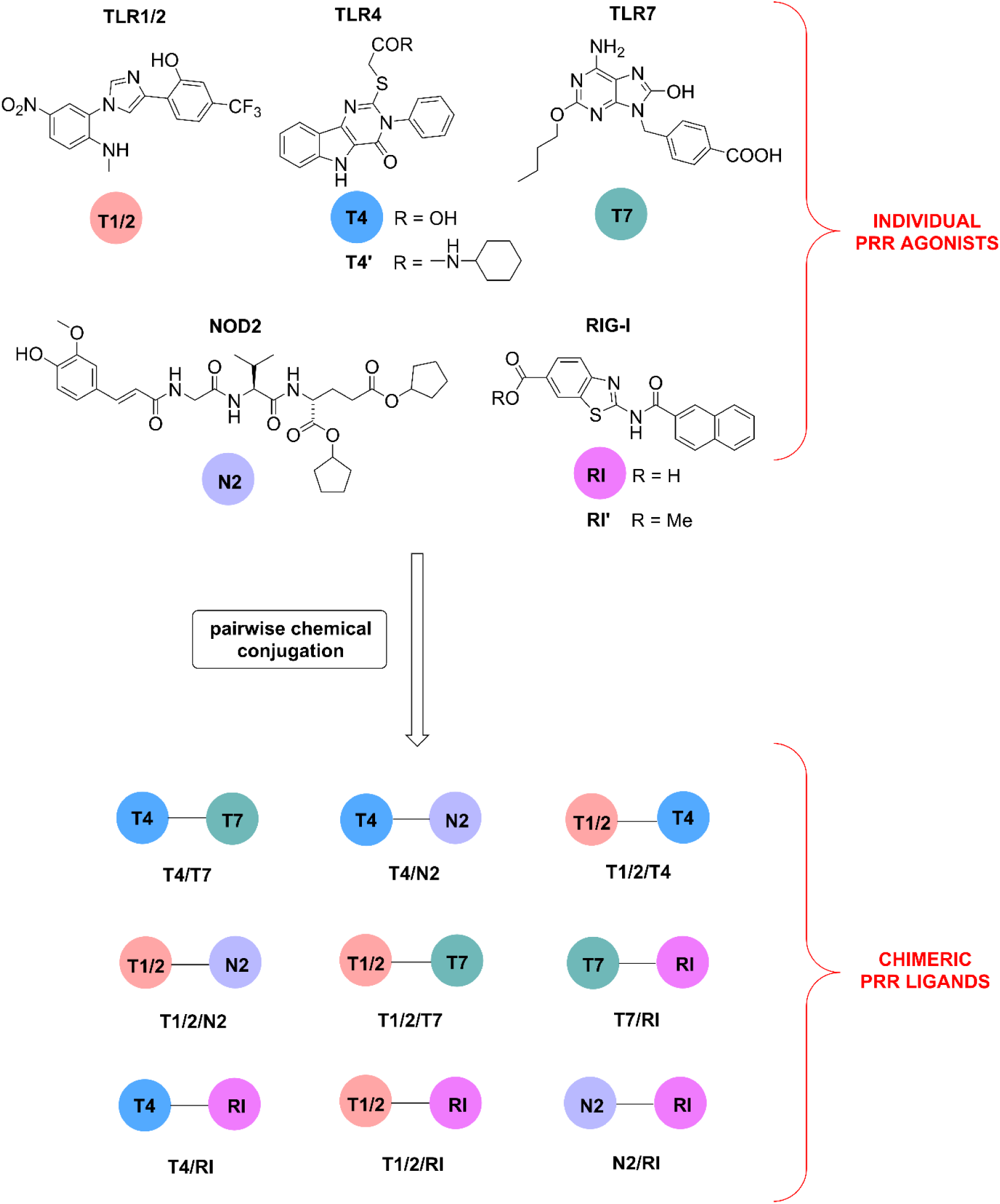
Chemical structures of the selected PRR agonists and their combinations featured in chimeric PRR ligands.

### Conjugated PRR ligands induce cytokine release from peripheral blood mononuclear cells

The immunostimulatory potential of the conjugates, single agonists, and unlinked mixtures of single agonists was first evaluated in human PBMCs, rather than the more commonly used reporter cell lines. Reporter cell lines have limited utility for conjugate screening because they express only specific innate receptors, lack metabolic activity, and thus pose a significant risk of false-negative results (*82*). Moreover, they do not accurately reflect the responses of primary leukocytes, let alone the complexity of the human immune system (e.g. lack of accessory cell effect). Given these limitations, the use of primary human cells, such as PBMCs, has recently been proposed for the discovery of novel immunoenhancing compounds (*83*). PBMCs represent a heterogeneous mixture of immune cell subpopulations, providing a physiologically relevant system for studying concomitant PRR activation. We tested our set of compounds at a concentration of 1 µM, as this was the level at which we previously observed the most distinct differences in our study of conjugated NOD2/TLR7 ligands (*13*). A similar concentration has also been used in previous studies (*84*). Conjugate **N2/T7** (*13*) and LPS served as the internal and external positive controls, respectively.

Our primary objective was to determine how the chimeric conjugates influence the cytokine response of innate immune cells. It is well established that no single *in vitro* parameter can reliably predict *in vivo* adjuvanticity. With this limitation in mind, we performed a comprehensive phenotypic screening using the BioLegend multiplex cytokine assay platform to assess the distinct immune fingerprints of conjugated agonists, both in terms of cytokine quantity and diversity. Additionally, cytokine profiles can provide insights into the nature of the immune response elicited. To this end, we simultaneously analyzed the supernatants of stimulated PBMCs for 13 cytokines and chemokines associated with Th1, Th2, and Th17 responses (Fig. 2A, Fig. S2A, Table S1). While treatment with single agonists or co-administration of individual ligands led to only modest increases in cytokine levels, certain chimeric molecules displayed clear synergistic effects. Notably, stimulation with conjugates **T4/T7** and **T7/RI** resulted in significantly higher cytokine secretion compared to their respective unlinked agonist mixtures, confirming that covalent conjugation enhances immune cell activation. Specifically, conjugates **T4/T7** and **T7/RI** elicited robust Th1 responses, as evidenced by the significant increases in IL-12p70, IFN-γ, and the IFN-inducible chemokine IP-10, with **T7/RI** showing slightly stronger effects than **T4/T7**. These cytokines show strong correlation with the IRF pathway activation and the induction of cellular immunity. IL-12p70 plays a key role in activating natural killer (NK) cells, stimulating IFN-γ production, and promoting Th1 and cytotoxic T lymphocyte polarization (*85*), while IP-10 functions as a chemoattractant for T cells and DCs (*86*). Although the proinflammatory responses induced by **T4/T7** and **T7/RI** were less pronounced, both conjugates increased IL-6, TGF-β1, and MCP-1 levels, while their effects on other cytokines were not significant. Interestingly, a previously reported TLR4/TLR7 agonist combination installed on a triazine core failed to exhibit synergy in NF-κB activation, cytokine secretion from BMDCs, or gene expression (*84*). Conjugate **T1/2/T7** also exhibited synergy, though to a lesser extent, while the remaining chimeric combinations failed to significantly boost cytokine release relative to unlinked mixtures. **T1/2/T7** synergistically induced IL-12p70 secretion and upregulated cytokines associated with type 2 response, such as IL-4 and TGF-β1. However, its effects on IL-10 and the Th17 cytokine IL-17A, though notable, did not reach statistical significance. Importantly, the levels of endogenous pyrogens TNF-α and IL-1β, which serve as baseline indicators of systemic inflammation and inflammasome activation, respectively, were not significantly elevated following stimulation with conjugates **T4/T7**, **T7/RI**, and **T1/2/T7**, suggesting a moderate systemic response. In contrast, conjugates **T4/T7**, **T7/RI**, **T1/2/T7**, and **T1/2/N2** induced substantial IL-6 secretion. While IL-6 is often considered an inflammatory marker, it plays a multifaceted role in immune regulation by stimulating B cell antibody production and directly influencing Th17, Tfh, and Treg cell differentiation (*87*).

**Fig. 2.**
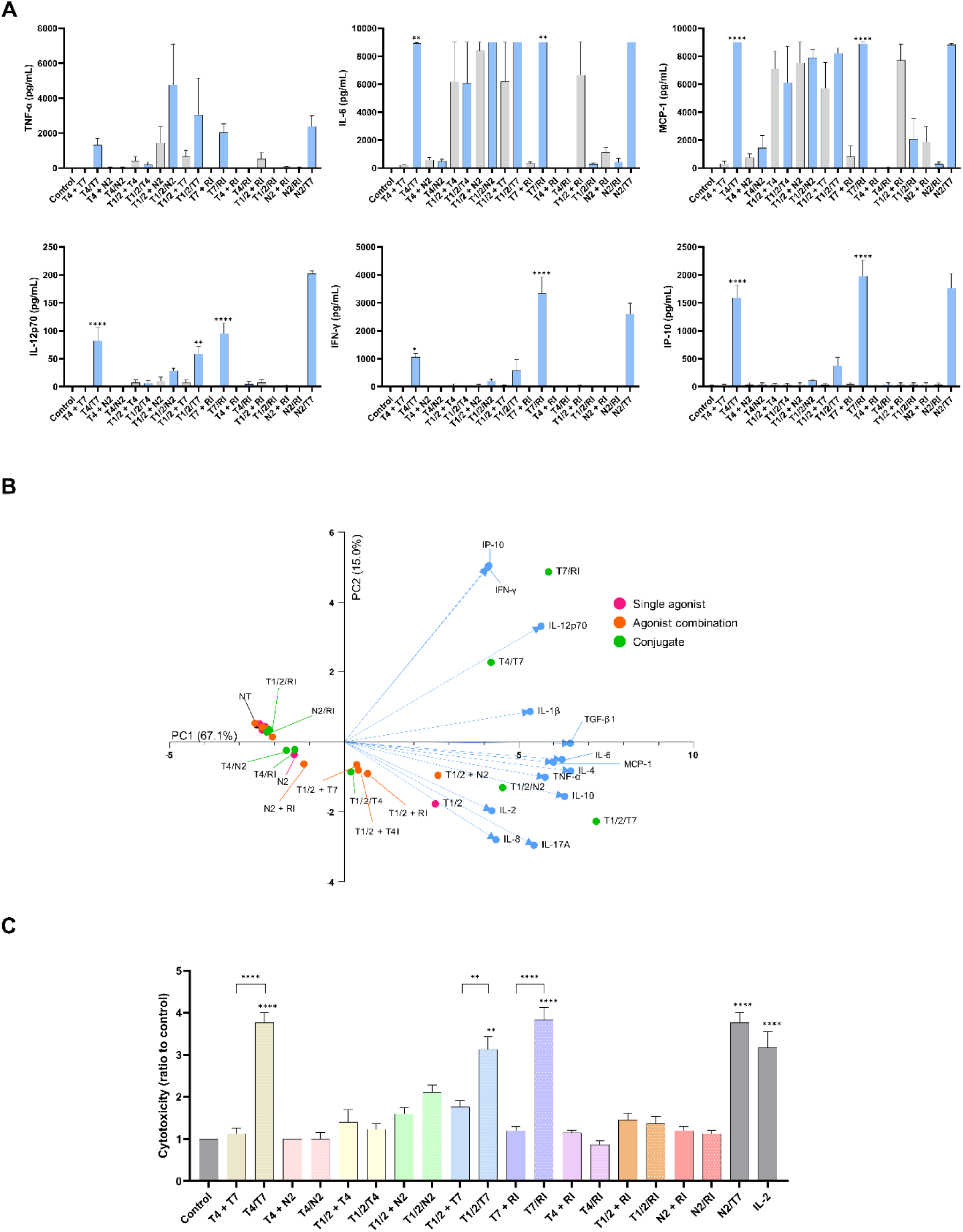
(A) Cytokine release from human PBMCs after 18 h stimulation with conjugated agonists (blue bars), unlinked mixtures (grey bars, both 1 µM), **N2/T7** (1 µM, positive control), or vehicle (0.1% DMSO). Data are mean ± SEM of three independent experiments. One-way ANOVA with Bonferroni’s test compared unlinked mixtures to conjugates. *, p < 0.05; **, p < 0.01; ***, p < 0.001; ****, p < 0.0001. (B) Principal component analysis (PCA) of cytokine/chemokine levels in PBMC supernatants after stimulation with conjugates, single agonists, and unlinked mixtures. PCA includes data from three donors. PC scores for stimulation conditions (red, orange, green) and cytokine loadings (blue) are shown. PCA captures 82.1% of variance (PC1: 67.1%, PC2: 15.0%). (C) PBMC cytotoxicity against K562 cells after 18 h treatment with conjugates, unlinked mixtures (1 µM), controls (**N2/T7**, 1 µM; IL-2, 200 U/mL), or vehicle (0.1% DMSO). Cytotoxicity was assessed after 4 h coincubation. Data are relative to the negative control (NT, 0.1% DMSO) and shown as mean ± SEM of three independent experiments. ANOVA with Bonferroni’s test compared unlinked mixtures to conjugates. **, p < 0.01; ***, p < 0.001; ****, p < 0.0001.

The remaining chimeric PRR ligands did not substantially increase cytokine levels compared to unlinked mixtures. For instance, the chimeric **T4/N2** ligand lacked immunoenhancing activity. This finding contradicts the report by Dong et al, which showed that conjugated NOD2/TLR4 agonist enhanced IL-6 secretion from BMDCs and promoted the maturation of APCs, ultimately translating to improved OVA-specific antibody and T cell responses (*16*). It is worth noting that NOD2 has shown the capacity to modulate the intensity of TLR4-mediated signaling, resulting in either synergistic stimulation of IL-12 production or its inhibition (*88*). Furthermore, the chimeric **N2/RI** ligand was also inactive, unlike the reported dual NOD2/RIG-I agonist dinucleotide SB 9200 (Inarigivir), which enhanced the efficacy of BCG vaccine against tuberculosis (*67*). The inactivity of **T1/2/T4**, **T1/2/RI**, and **T4/RI** ligands aligns with the findings of Pandey et al, who demonstrated that all pairwise combinations of TLR2, TLR4, and RIG-I agonists lacked synergy on T cell proliferation in a BMDC/T-cell coculture experiment (*89*). Additionally, both synergy and cross-tolerance have been observed between TLR2- and TLR4- mediated signaling pathways (*90, 91*), while primary activation of RIG-I desensitizes the TLR2 activation (*92*). Conversely, the same combinations of TLR2, TLR4, and RIG-I agonists have also shown potential to generate amplified immune responses (*69*).

It is important to note that the activity of innate immune agonists does not always correlate directly with the quantity of cytokines produced by PBMCs. Instead, it is a complex function influenced by multiple secreted cytokines and paracrine signaling (*93*). To better understand the architecture of the induced cytokine responses, we performed principal component analysis (PCA) on the dataset to reduce the dimensionality of the data and ease its interpretation. This dataset included cytokine distributions from conjugates, unlinked agonist mixtures, and single agonists across 13 variables. PCA revealed strong stimulus-specific clustering, with principal components PC1 and PC2 accounting for 82% of the total variance. The contributions of individual protein analytes to the two principal component axes are detailed in Table S2A. PC1 was driven by a core signature of both innate and adaptive immune responses, while PC2 differentiated the type of response: positive PC2 values were associated with IP-10, IFN-γ, and IL-12p70, whereas negative PC2 values corresponded to IL-2, IL-8, and IL-17A. As illustrated in the PCA plot (Fig. 2B), we were able to distinguish the most distinct conjugates based on their cytokine signatures (Table S2B). Conjugates **T7/RI** and **T4/T7** strongly induced IP-10, IFN-γ, and IL-12p70, while also enhancing the release of cytokines and chemokines from the PC1 group. In contrast, conjugates **T1/2/N2** and, particularly, **T1/2/T7** exhibited a unique profile, inducing cytokines and chemokines from PC1 while also enhancing those linked to negative PC2 values (i.e., IL-2, IL-17A). To prioritize compounds with the most pronounced impact on immune responses, we defined a selection criterion using a three-unit PCA radius centered around control (NT). Compounds positioned outside this boundary— **T4/T7**, **T7/RI**, **T1/2/T7** and **T1/2/N2** —were identified as having the most distinctive immune signatures. Among these, **T1/2/T7** and **T1/2/N2** exhibited particularly unique profiles, and could be further discriminated based on levels of induced cytokines and chemokines.

### Conjugated PRR ligands enhance PBMC cytotoxicity against cancer cells

To evaluate the ability of conjugates to induce an antitumor immune response, we included an additional functional assay—the PBMC cytotoxicity assay. We assessed the effects of stimulation with single agonists (Fig. S2B), their unlinked mixtures, and chimeric conjugates on the nonspecific cytolytic activity of PBMCs against K562 cancer cells (Fig. 2C, Table S3). Instead of using isolated NK cells, we employed the entire PBMC population to account for the contributions of accessory immune cells that respond to PRR stimulation and interact with NK cells through cytokine secretion. Consistent with the strong immunostimulatory activity observed for conjugates **T4/T7** and **T7/RI** in the PBMC cytokine assay, these conjugates also significantly enhanced PBMC cytolytic activity. In contrast, conjugate **T1/2/T7** exhibited a less pronounced but still notable effect, while in case of **T1/2/N2** the increase in cytolytic activity was not statistically significant. Single-agonist treatments and co-stimulation with unlinked agonist mixtures failed to induce any increase in PBMC cytotoxicity, highlighting the superior potency of chimeric conjugates in promoting immune-mediated tumor cell killing. To evaluate potential direct cytotoxic effects, PBMCs and cancer cells were independently treated with all compounds and mixtures. No significant increase in cell death was observed, confirming that the enhanced cytotoxicity induced by conjugates **T4/T7**, **T7/RI**, and **T1/2/T7** resulted from PBMC activation rather than direct toxicity and that conjugates themselves are not cytotoxic to human PBMCs (Table S3). Notably, the same conjugated PRR ligands that demonstrated the strongest immunostimulatory activity in the PBMC cytokine assay also emerged as the most promising in this assay. Having successfully completed our primary screening, we proceeded to investigate additional characteristics of this subset of conjugates to identify the most suitable candidates for preliminary *in vivo* studies.

### RNA sequencing reveals differential gene expression in conjugate-stimulated PBMCs

To gain deeper insights into the transcriptional effects induced by the conjugated agonists, PBMC mRNA was isolated, amplified, and sequenced following overnight stimulation with selected conjugates, with the vehicle serving as a negative control. Differential expression analysis revealed that chimeric ligands **T4/T7**, **T7/RI**, **T1/2/T7** and **T1/2/N2** significantly upregulated the transcription of 1,719, 1,484, 1,051, and 541 genes, respectively, while downregulating 1,538, 1,443, 854, and 629 genes compared to unstimulated control samples (Fig. 3A).

**Fig. 3.**
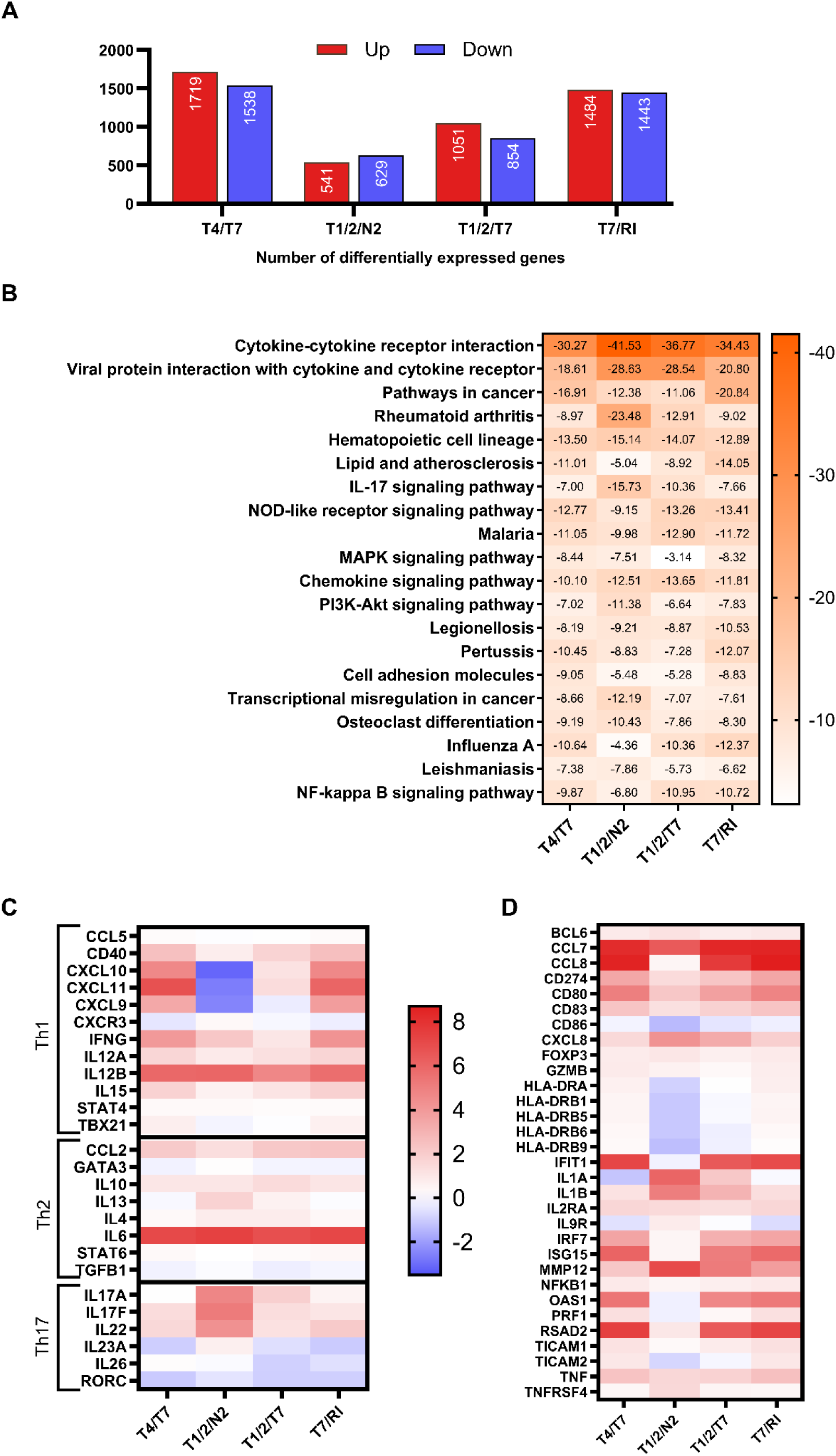
(A) Number of significantly upregulated/downregulated genes in PBMCs from three donors after 18 h treatment with conjugates (1 μM), using a false discovery rate <0.1 and fold-change >2 or <0.5 vs. vehicle (0.1% DMSO). (B) Heatmap of 20 top enriched KEGG terms colored by log(q value). Differentially expressed genes from Fig. 3A were used as the input for pathway enrichment analysis by Metascape. (C, D) Transcriptomic profiles of conjugate-stimulated PBMCs. Heatmaps show log_2_(fold change) in gene expression relative to unstimulated control: (C) Th1, Th2, and Th17 response genes; (D) other relevant differentially expressed genes. Red indicates upregulation, blue downregulation.

Pathway enrichment analysis of differentially expressed genes, conducted using the Kyoto Encyclopedia of Genes and Genomes (KEGG) database, identified several innate immune system-related pathways. All conjugates exhibited enrichment of the “cytokine-cytokine receptor interaction”, “viral protein interaction with cytokine and cytokine receptor” and “hematopoietic cell lineage” pathways. Additionally, **T1/2/N2** markedly enriched the “IL-17 signaling pathway” compared to other conjugates (Fig. 3B). We classified the synergized transcripts into four primary clusters (Figs. 3C, 3D): (i) genes associated with the Th1 immune response, (ii) genes linked to the Th2 response, (iii) genes defining the secretory phenotype of Th17 cells, and (iv) other immune-related genes that provide additional insights into immune signatures. In line with the results obtained at the protein level, stimulation with **T4/T7** and **T7/RI**, and to a lesser extent **T1/2/T7** and **T1/2/N2**, significantly enriched genes related to cytokine signaling. Both **T4/T7** and **T7/RI** strongly induced the transcription of IFN-γ and both IL-12 subunits (IL12A, IL12B), the hallmark cytokines of the Th1 response, alongside IL15, which is also associated with Th1 immunity (*94, 95*). This Th1 polarization was further reinforced by the upregulation of the T-bet transcription factor (TBX21), the master regulator of Th1 cell differentiation (*96*), as well as IFN-γ–inducible chemokines CXCL9, CXCL10, and CXCL11, which act through the CXCR3 receptor expressed on Th1 and NK cells (*97*). Importantly, genes for granzyme B (GZMB) and perforin (PRF1), which are indicative of the lytic activity of NK and CD8^+^ T cells (*98*), were also markedly upregulated. A moderate increase in Th17-associated genes (IL17F, IL22) (*99*) was observed, whereas genes linked to the Th2 response (IL4, IL13, STAT6, GATA3) (*100*) remained largely unchanged, with only minor effects on CCL2, IL6, and IL10, which promote Th2 differentiation. Additionally, **T4/T7** and **T7/RI** upregulated genes involved in IFN-signaling (IFIT1, IRF7, ISG15, OAS1, RSAD2) and chemokine recruitment (CCL7, CCL8), further supporting their role in activating Th1-biased immune responses, as well as DC maturation markers (CD40, CD80, CD83, and HLA-DR). In contrast, **T1/2/N2** had no significant impact on Th1-associated genes except for the upregulation of IL12B and, to a lesser extent, IFN-γ. Notably, key Th1 chemokines (CXCL9, CXCL10, CXCL11) were markedly downregulated. While prototypical Th2 genes (IL5, STAT6, GATA3) remained mostly unchanged, moderate increases in IL4 and IL13 were observed. Unlike the other conjugates, **T1/2/N2** selectively induced the transcription of IL-17A, IL-17F, and IL-22, which collectively define the Th17 secretory phenotype, as well as IL1B, IL23A and IL6, which are essential for Th17 differentiation (*101*). In addition, IL9R and OX40 (TNFRSF4), a co-stimulatory receptor implicated in long-term T cell immunity (*102*) and IL-9-polarized response (*103*), were also selectively upregulated. Given that the IL-9/IL-9R axis contributes to Th17 differentiation (*104*), these data strongly suggest that **T1/2/N2** preferentially activates the Th17 T cell subset. Additionally, MMP12, IL1B, and CXCL8 were significantly upregulated, whereas DC maturation markers CD80 and CD83 were only weakly increased, and CD86 was downregulated. **T1/2/T7** exhibited a distinct immune profile, characterized by weak induction of canonical Th1 genes (IL12A, IL12B, IFNG, CXCL10, CXCL11) and Th1-associated IL15. Moderate increases in IL17A, IL17F, and IL22 further suggested Th17 activation, supported by upregulation of IL1B and IL6 (although IL23A was downregulated). **T1/2/T7** displayed a weak induction of Th2-related genes (IL4, IL13, CCL2) and the Th2-promoting cytokine IL10. Similar to **T4/T7** and **T7/RI**, and likely due to the shared TLR7 agonist moiety, **T1/2/T7** significantly upregulated IFN-signaling-dependent genes (IFIT1, IRF7, ISG15, OAS1, RSAD2), chemokine attractants (CCL7, CCL8, CCL2), and DC maturation markers CD80 and CD83.

### Conjugates T4/T7 and T7/RI are taken up by PBMCs and exhibit receptor-driven activities

Next, we addressed the cell uptake of best performing conjugates **T4/T7** and **T7/RI** in comparison to single agonists under the conditions employed as well as the stability of conjugates towards the cells’ enzymatic system. To that end, we collected the extracellular media and cell lysate extracts following overnight stimulation of PBMCs with conjugates (**T4/T7**, **T7/RI**) as well as single agonists (**T4**, **T7**, and **RI**) for 18 h (all at 1 µM) and performed a comprehensive LC-MS analysis (Table S4A). First, the measured concentrations indicated that both conjugates were taken up by the cells, but **T4/T7** crossed the membrane to a lesser extent. Specifically, the extracellular and lysate concentrations were found to be 667.22 and 52.44 ng/mL for **T4/T7**, and 209.44 and 126.21 ng/mL for **T7/RI**, respectively. In contrast, the single agonists failed to cross the membrane, supporting the notion that conjugation allows for successful internalization (though it should be noted that the target of **T4** is extracellular). As shown in Tables S4B and S4C, in addition to the parent conjugates, we detected intracellular metabolites resulting from the hydrolytic cleavage of the amide bonds, including single agonists and linker-modified agonists (see Figs. S3 and S4). Our initial findings suggest that hydrolysis of amide linkages in both conjugates occurs in PBMCs, producing several metabolites that may collectively contribute to PRR stimulation in a synergistic manner. However, it is important to note that the extent of hydrolysis is relatively low, and intact conjugates remain the predominant intracellular species.

Therefore, in order to shed some light on the role of each PRR ligand featured in conjugates **T4/T7** and **T7/RI** in directing PBMC responses, we conducted preliminary mechanistic studies. To differentiate responses of conjugated PRR ligands toward their envisaged targets, we employed a panel of corresponding signaling inhibitors. Specifically, to decouple the effects of individual moieties featured in conjugated TLR4/TLR7 (**T4/T7**) and TLR7/RIG-I ligands (**T7/RI**), we performed studies in the presence of their *bona fide* inhibitors TAK242 (TLR4 antagonist) (*105*), M5049 (TLR7 antagonist) (*106*) and MRT67307 (RIG-I signaling inhibitor) (*107*), as well as combinations of these inhibitors. First, we examined whether the addition of each signaling inhibitor at a non-toxic concentration diminished cytokine production (Fig. 4A, Table S5A). The addition of either the TLR4 or TLR7 antagonist reduced the activity of **T4/T7**. A similar observation has been made in case of **T7/RI** following co-treatment with the TLR7 antagonist or the RIG-I signaling inhibitor. The combination of both signalling inhibitors further decreased cytokine release induced by both conjugates, suggesting that the activities of **T4/T7** and **T7/RI** were reliant on both TLR4 and TLR7, and TLR7 and RIG-I, respectively. Co-treatment with signaling inhibitors yielded comparable results in a functional assay, where they completely abolished the **T4/T7**- and **T7/RI**-induced cytolytic activity of PBMCs against K562 cancer cells (Fig. 4B, Table S5B).

**Fig. 4.**
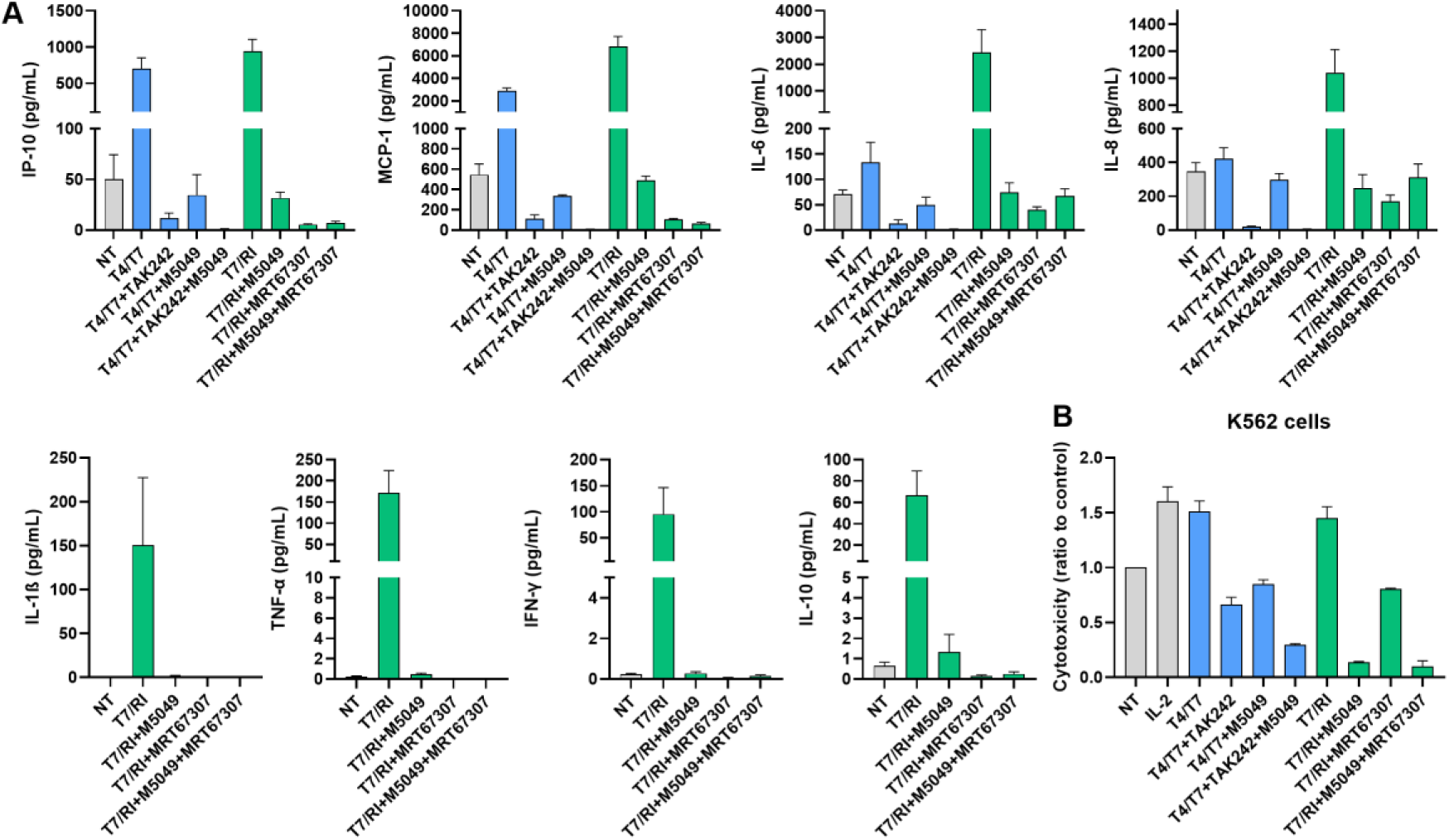
(A) Effect of signaling inhibitors on conjugate-induced cytokine release from human PBMCs. PBMCs were pretreated with corresponding inhibitors. Cytokine concentrations were measured after 18 h stimulation with conjugates (100 nM), inhibitors (5 μM) or the corresponding vehicle (0.1% DMSO). Data are expressed as mean ± SEM of two independent experiments. (B) Effect of signaling inhibitors on the cytotoxic activities of PBMCs against K562 cells. PBMCs were treated for 18 h with conjugates (100 nM), antagonists (5 μM) or the corresponding vehicle (0.1% DMSO), before the addition of K562 cells. Cytotoxicity was determined after a 4 h co-incubation. Each experiment was conducted in duplicate and repeated in two independent biological replicates. Data are shown as relative activities to the negative control (NT, 0.1% DMSO) and are means ±SEM of two independent experiments.

### Conjugated PRR ligands demonstrate *ex vivo* adjuvant activity

After demonstrating the efficacy of the conjugates in eliciting nonspecific immune responses, we next focused on their potential to enhance the development of antigen-specific immunity. A key feature of adjuvants is their ability to stimulate DCs, which, when mature and activated, instruct T and B cells to initiate effective and targeted adaptive immune responses. DC/T-cell coculture models provide an integrated readout of the signals derived from DCs, which are regulated by the combinatorial activation of PRR pathways. To assess how dual stimulation influences antigen presentation by DCs and the generated T cell responses, we evaluated the conjugates in an *ex vivo* coculture system with murine BMDCs and naïve ovalbumin (OVA)-specific CD4^+^ and CD8^+^ T cells, isolated from OT-II and OT-I mice, respectively. BMDCs were treated with conjugates and soluble OVA protein, washed, and then cocultured for 3 days with carboxyfluorescein succinimidyl ester (CFSE)-labeled T cells. To quantify the effects of the compounds, we used the proliferation index as a proxy for T cell response, which corresponds to the average number of divisions per responding T cell in the coculture. Pre-activation of BMDCs with conjugated PRR ligands significantly enhanced their ability to induce antigen-specific activation and proliferation of both CD4^+^ and CD8^+^ T cells, as measured by upregulation of the early activation marker CD25 and CFSE dilution (Figs. 5A and 5B, Table S6A). While all conjugates activated CD8^+^ T cells similarly and outperformed stimulation with LPS, conjugates **T1/2/T7** and **T1/2/N2** slightly surpassed **T4/T7** and **T7/RI** in their ability to activate CD4^+^ T cells.

**Fig. 5.**
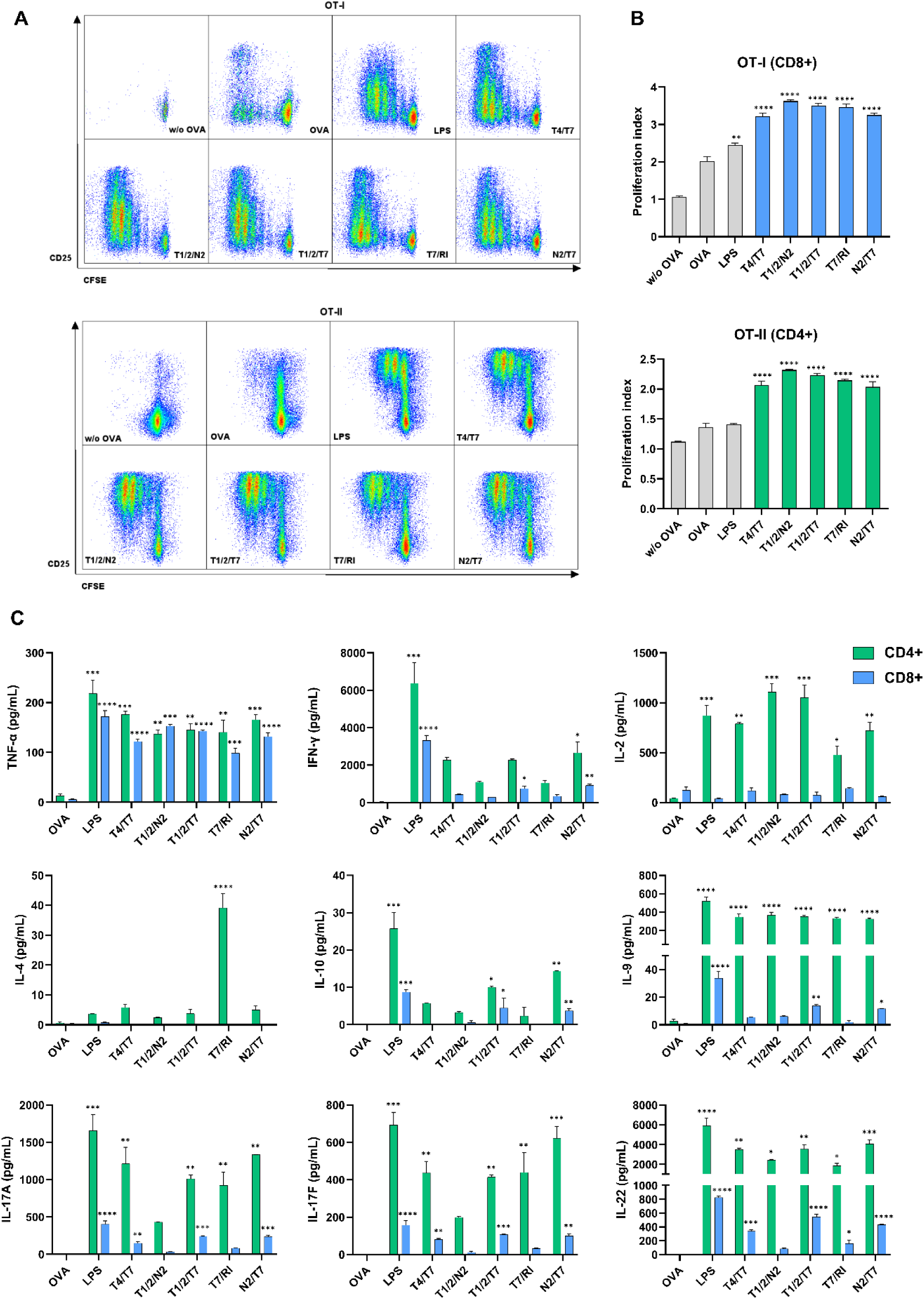
Treatment of BDMCs with conjugates promotes their antigen presenting activity and enhances antigen-specific activation and proliferation of CD4^+^ and CD8^+^ T cells. BMDCs from C57BL/6 mice were treated for 18 h with conjugates (1 μM), LPS (1 μg/mL), or vehicle (0.1% DMSO) in the presence of OVA (50 μg/mL). CFSE-labeled OVA-specific CD4+ or CD8+ T cells (isolated from OT-II or OT-I mouse splenocytes, respectively) were added to the treated and washed BMDCs and cocultured for 72 h. (A) Representative dot plots show CD25 expression and CFSE dilution in live Thy1.2+/CD4+ (OTII) and Thy1.2+/CD8+ (OTI) T cells. (B) Pooled data from (A), expressed as proliferation indexes. (C) Cytokine concentrations in BMDC/T-cell coculture supernatants following 72 h coincubation. Data are mean ± SEM of duplicates of two independent experiments. *, p < 0.05, **, p <0.01, ***, p <0.001, ****, p <0.0001 versus vehicle-treated control. Statistical significance was determined using one-way ANOVA with post-hoc Dunnett’s test comparing conjugates versus OVA.

Subsequent analysis of the cytokine profiles in the supernatants of BMDC/T-cell cocultures (Fig. 5C, Fig. S5, Table S6B) revealed notable differences between the conjugates, particularly in terms of cytokine production by CD8^+^ T cells. While all conjugates induced the secretion of TNF-α, stimulation with **T1/2/T7** resulted in significantly elevated levels of Th2 cytokines (IL-9, IL-10, IL-13) and Th17 cytokines (IL-17A, IL-17F, IL-22), along with a modest increase in Th1-associated IFN-γ. A similar, though less pronounced, Th2/Th17-polarized profile was observed with **T4/T7**. In contrast, the cytokine profiles elicited from CD4^+^ T cells were more similar across the conjugates. Stimulation with all conjugates increased TNF-α and IL-2 secretion, with the highest levels of IL-2 observed for conjugates **T1/2/T7** and **T1/2/N2**, followed by **T4/T7** and **T7/RI**. Conjugates **T1/2/T7**, **T4/T7** and **T7/RI** also significantly elevated the levels of Th2 cytokines (IL-9, IL-10, IL-13) and Th17 cytokines (IL-17A, IL-17F, IL-22), with **T7/RI** being the only conjugate that elicited detectable levels of IL-4. The effects of conjugate **T1/2/N2** on Th2- and Th17-associated cytokines were less pronounced, which contrasts with the results obtained in PBMCs, where **T1/2/N2** induced the most notable Th17-biased response. Overall, the results from this murine model were not entirely consistent with those observed in human PBMCs. Unlike the Th1-biased cytokine profiles induced by **T4/T7** and **T7/RI** in human PBMCs—supported by transcriptomic analysis—murine BMDCs stimulated with these conjugates elicited prominent levels of Th1 cytokines but also induced some Th2/Th17 cytokines. These differences are likely due to the fact that murine BMDCs respond poorly to TLR7 ligands (*108*), as well as the absence of the accessory cell effects observed in PBMCs.

### Conjugated PRR ligands exhibit *in vivo* adjuvant activity

Encouraged by the promising *in vitro* and *ex vivo* results, we next evaluated the adjuvant potential of the conjugates in a murine vaccination model *via* measurement of B cell responses. Mice were vaccinated using a prime-boost regimen with OVA as the model antigen and conjugates **T4/T7**, **T7/RI**, **T1/2/T7**, and **T1/2/N2** as adjuvants (doses were equimolar to 50 μg of **N2/T7** to enable head-to-head comparison and administered on days 0, 21, and 42). Alum (100 μg) was used as a universal benchmark (*109*). OVA-specific IgG, IgG1, and IgG2a antibody responses were measured to delineate B cell responses (Fig. 6A, Table S7). Conjugates **T4/T7** and **T7/RI** induced strong systemic immune responses, with OVA-specific IgG titers several orders of magnitude higher than those observed in mice immunized with OVA alone or with alum. The impact of **T1/2/N2** and **T1/2/T7** on total IgG titers was less pronounced but still notable. Importantly, immunization with **T4/T7** and **T7/RI** also increased the titers of both IgG1 and IgG2a, indicative of Th2 and Th1 immune responses, respectively.

**Fig. 6.**
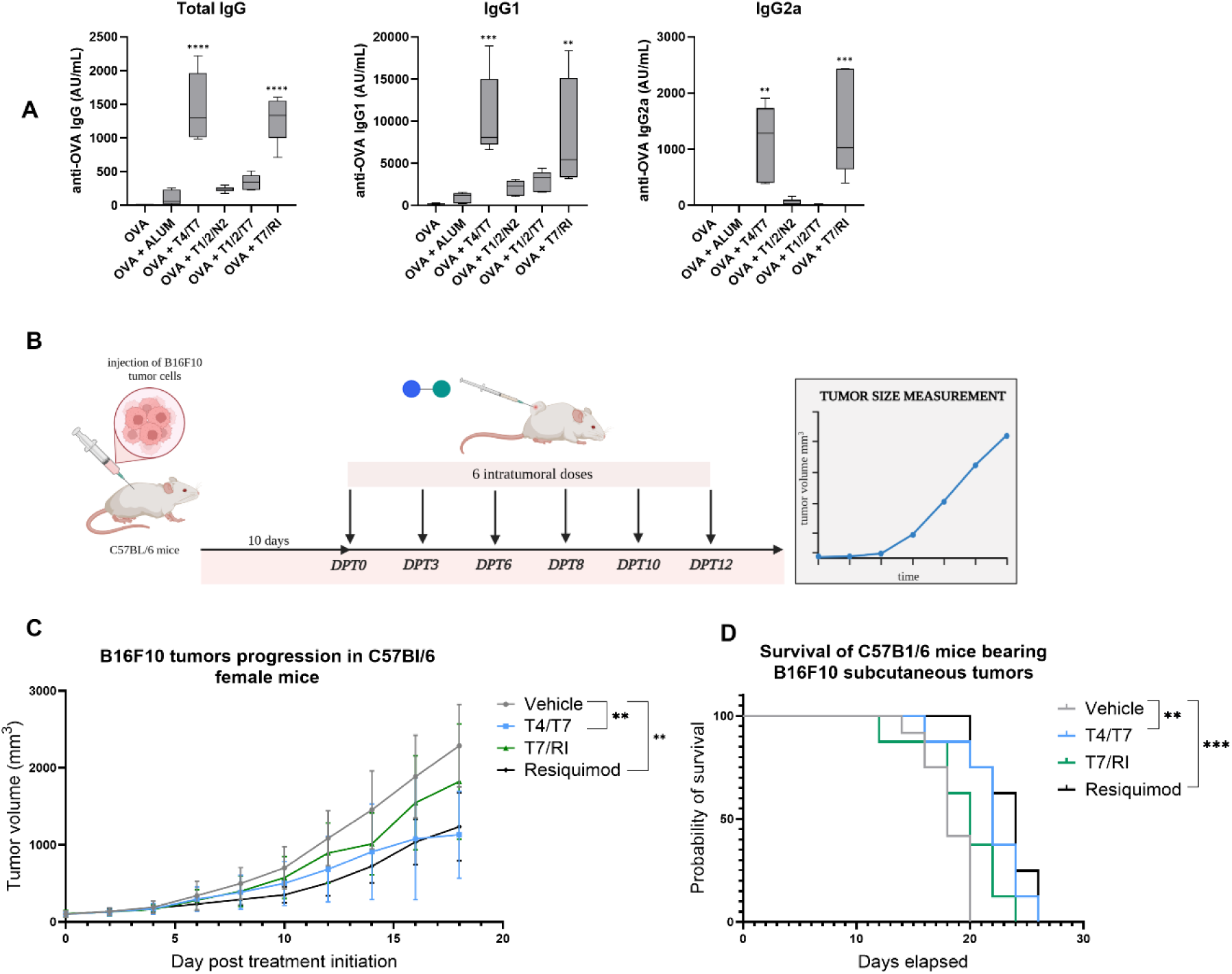
(A) *In vivo* adjuvant activity of conjugates. NIH/OlaHsd mice (n = 6/group) were immunized s.c. on days 0, 21, and 42 with OVA (10 μg) alone or with conjugates (equimolar to 50 μg **N2/T7**) or Al-hydrogel (100 μg). OVA-specific IgG, IgG1, and IgG2a responses were measured 7 days after the final dose; **, p < 0.01, *** p < 0.001, **** p < 0.0001 vs. control (one-way ANOVA, Dunnett’s test). (B) Tumor challenge experiment design. C57BL/6 mice (n = 8/group) were injected with 5 × 10⁵ B16F10 tumor cells. After tumors reached ∼80 mm³ (day 10), mice were randomized and treated intratumorally with vehicle, **T4/T7**, **T7/RI**, or resiquimod every 2–3 days for six doses. (C) Tumor growth. Volume was recorded every other day until tumors exceeded 3000 mm³. Data (DPT0–DPT18) are mean ± SEM. Tumor growth was analyzed using two-way ANOVA with a linear mixed-effects model; * p < 0.05, ** p < 0.01. (D) Survival analysis. Kaplan-Meier survival curves (DPT0–DPT26) were analyzed using the Mantel–Cox log-rank test. **, p < 0.01; ***, p < 0.001 vs. control.

In contrast, the TLR1/2 agonist-featuring conjugates **T1/2/N2** and **T1/2/T7** had a stronger effect on IgG1 than on IgG2a, suggesting a shift toward predominantly Th2-polarized responses. This is consistent with prior reports showing that TLR2 activation *in vivo* can induce antigen-specific responses with a Th2-polarized profile. However, our findings differ from previous studies where co-engagement of TLR2 with Th1-polarizing TLRs or Th2-polarizing NOD2 enhanced Th1-type responses. For instance, the co-administration of HIV-1 p24 antigen with the nanoformulated dual TLR2/6/TLR7 ligand CL413 significantly increased anti-p24 IgG titers, with IgG1 (Th2-associated) as the predominant subclass, although IgG2a (Th1-associated) was also notably elevated (*11*). Similarly, the dual TLR2/6/NOD2 ligand CL429 induced a balanced Th1/Th2 immune profile while also promoting mucosal immunity, as indicated by increased IgA levels (*12*). It is worth noting that in both cases, the p24 antigen was used, the conjugates were encapsulated in nanoparticles, and a TLR2/6 agonist was employed instead of a TLR1/2 agonist. These factors, individually or in combination, may have contributed to the observed differences between our results and previous studies. As expected, alum induced a Th2-skewed response, consistent with historical data. Interestingly, the *in vivo* adjuvant activity data do not support our findings obtained in BMDC/T-cell assay, while they are in perfect agreement with the RNA sequencing analysis results generated on PBMCs (expression of CD80, CD83, CD86, and HLA-DR) thus attesting to the importance of accessory cells. Collectively, these results underscore the potential of conjugates **T4/T7** and **T7/RI** as vaccine adjuvants capable of eliciting both cellular and humoral immune responses in a balanced manner. Promotion of both types of responses has recently also been highlighted as a new paradigm for cancer immunotherapy (*110*).

### Conjugate T4/T7 exhibits antitumor activity *in vivo*

Based on the results obtained in the PBMC-K562 assay and RNA sequencing analysis, we narrowed our focus to evaluating the capacity of conjugates **T4/T7** and **T7/RI** to attain antitumor response in a B16F10 tumor model (*111*). Their therapeutic efficacy was assessed in comparison to resiquimod – a potent imidazoquinoline TLR7/8 agonist (*112*) exhibiting antitumor activity in a range of murine tumor models (*113–115*). Mice were injected with B16F10 melanoma tumor cells, and tumor growth was monitored (Fig. 6B). Four groups of mice were treated via intratumoral injection with vehicle, resiquimod at a low dose of 20 µg/mouse (*i.e.* 1 mg/kg) (*113*) as the positive control, and **T4/T7**, **T7/RI** (at doses equimolar to resiquimod) every two or three days (six injections in total) to maintain a refractory state over a two-week period. Tumor volume was monitored every two days during the treatment and until the study ended (Table S8A). Treatment with **T4/T7** and resiquimod resulted in statistically significant suppression of tumor growth (Fig. 6C), as determined by the ANOVA mixed effects model (p = 0.0033 and p = 0.0014, respectively, *vs.* vehicle). Tumor growth in the **T4/T7**- and resiquimod-treated groups was significantly inhibited during the treatment period and delayed after treatment cessation. Specifically, **T4/T7** exhibited a tumor growth inhibition (TGI) of 59%, while resiquimod showed a TGI of 56% on day 18. The effect of **T7/RI** was less pronounced, with a TGI of 31%. Furthermore, the calculated tumor/control (T/C) ratios on day 18 (DPT18: 48% for **T4/T7** vs. 53% for resiquimod) further demonstrated that **T4/T7** exhibited superior antitumor efficacy (Table S8B). Lastly, statistically significant increases in survival time were observed for the **T4/T7** and resiquimod groups compared to the vehicle group (p = 0.0019 and p = 0.0004, respectively; Fig. 6D and Table S8C). To preliminarily assess the toxicity of the conjugates, mouse body weights were continuously monitored throughout the treatment period, with no significant weight changes observed in any group (Fig. S6). Despite the B16F10 melanoma model being poorly immunogenic and highly aggressive, our results clearly demonstrate the antitumor efficacy of T4/T7 as a standalone immunotherapeutic. Intratumoral treatment of more immunogenic ‘hot’ tumors (e.g., CT26, MC38), which harbor higher concentrations of tumor-infiltrating lymphocytes (*116*), is likely to be even more effective.

## 3 Discussion

Herein, we took advantage of the interdependence of immune signals to explore the potential of manipulating the complex PRR signaling architecture by integrating PRR ligands into chimeric conjugates. We also implemented a unique two-step phenotypic screening approach using primary human PBMCs, allowing us to capture the complexity of multi-input effects (*i.e.* paracrine interactions between cells via cytokine secretion) to identify conjugates with innate immune-enhancing activity *in vitro*, as well as adjuvanticity and antitumor activity *in vivo*. Chimeric molecules exhibited distinct cytokine profiles and functional activities compared to their unlinked counterparts, underscoring the critical role of covalent linkage in eliciting higher-order immune effects. Our preliminary findings indicate that although a minor fraction of individual PRR ligands is released, the amide linkages remain relatively stable under the experimental conditions employed. Consequently, the conjugated forms persist as the predominant species. This is particularly important, as co-stimulation with an unlinked mixture of agonists caused individual cells to respond to only one ligand or the other, rather than integrating both signals (*117*). Such non-integrative processing suggests that simultaneous engagement of two PRRs is less effective when agonists are delivered in solution compared to covalently linked formats. Covalently linked agonists ensure co-presentation to the same immune cell, thereby enhancing activation and promoting PRR cross-talk within a single cell, ultimately resulting in a more robust immune response. Additionally, the large size of chimeric molecules may further facilitate recognition and uptake by immune cells. This is exemplified by the conjugated NOD2/TLR7 ligand CL325, which has been reported to self-assemble into liposomal or micellar structures (15), likely contributing to improved *in vivo* efficacy.

Overall, conjugate **T4/T7** elicited a pronounced Th1-skewed immune response *in vitro* while maintaining moderate proinflammatory effects. However, in an *in vivo* setting, it induced a balanced Th1/Th2 immune response against a model antigen, consistent with previous reports. Notably, formulated combinations of TLR4 and TLR7 agonists have been shown to promote balanced Th1/Th2 immune responses to influenza antigens (*57, 59, 60, 66*) and to drive Th1-biased responses against SARS-CoV-2 antigens (*118, 119*). Conjugate **T7/RI** induced a robust Th1 immune response both *in vitro* and *in vivo*, aligning with previous findings. For instance, the RNA-based TLR7/RIG-I dual agonist CV8102—while not a chimeric construct but rather a merged pharmacophore—significantly enhanced the immunogenicity of a rabies vaccine in human trials (*68*). Additionally, the co-delivery of TLR7 and RIG-I agonists via nanoparticles has been shown to generate potent cellular responses in influenza vaccination models (*72, 73*). Consistent with the strong Th1-skewed immunostimulatory activity of conjugates **T4/T7** and **T7/RI** observed in cytokine assays, both also up-regulated the expression of granzyme B and perforin and amplified the cytolytic activity of PBMCs against cancer cells. Furthermore, intratumoral administration of these conjugates revealed superior antitumor efficacy of **T4/T7** compared to **T7/RI** and resiquimod in a B16F10 tumor model. Our findings support previous studies which have demonstrated that intratumoral delivery of TLR4 and TLR7/8 agonists can synergize with checkpoint inhibitors to eradicate CT26 tumors (*65*). On the other hand, the *in vitro* cytolytic activity of **T7/RI** did not translate to antitumor activity *in vivo*, as opposed to CV8102 that showed efficacy in clinical studies for patients with hepatocellular carcinoma and in combination with PD-1 blockade for solid tumors (*120, 121*).

Conjugate **T1/2/T7** exhibited a pronounced proinflammatory signature, inducing a mixed but relatively weak immune response in PBMCs, characterized by features of Th1, Th2, and Th17 phenotypes. *In vivo*, however, it generated only a weak Th2-skewed response. The previously reported dual TLR2/6/TLR7 ligand CL413 primarily stimulated IL-6, IL-8, and IL-12 secretion from PBMCs, yet it demonstrated strong humoral and cell-mediated immune responses as a vaccine adjuvant (*11*). Additionally, co-delivery of TLR2/6 and TLR7 agonists encapsulated in nanostructured lipid carriers has been shown to enhance the adjuvant activity of HepB and influenza vaccines (*56*). In contrast to the other conjugates, **T1/2/N2** selectively induced a Th17 immune response in PBMCs. This is particularly notable as Th17 cells play a crucial role in host defense against bacterial, fungal, and viral infections at mucosal surfaces by promoting local IgA responses (*122, 123*). Furthermore, Th17-inducing DC vaccines have been shown to enhance antitumor immunity (*124*). Our findings in PBMCs align with previous studies demonstrating that TLR1/2 agonists promote Th17 cell differentiation and proliferation *in vitro*, leading to increased Th17 cytokine production (*125, 126*). Notably, NOD2 has been shown to be more potent than TLRs in driving IL-17 production in human memory T cells through IL-1β and IL-23 secretion from DCs. However, while NOD2 alone was inactive, it acted synergistically with the TLR1/2 ligand Pam3CSK4 to enhance Th17 responses (*127, 128*). Conjugate **T1/2/N2** elicited only a weak immune response in our murine vaccination model, which contrasts with previous reports of a chimeric TLR2/6/NOD2 ligand inducing high antigen-specific IgA and IgG titers at both systemic and mucosal sites following parenteral immunization (*12*). This discrepancy is likely due to differences in the TLR2 agonist moiety, the choice of antigen, and the formulation, as the TLR2/6/NOD2 ligand was delivered in nanoparticles.

Our study has several strengths, including the use of an innovative phenotypic screening approach with primary human PBMCs, the development of chimeric molecules that take advantage of the synergistic cross-talk between innate immune receptors, and the establishment of guidelines for immunostimulant design with distinct Th17 biasing properties. Additionally, we assessed both the adjuvant and antitumor activity of these conjugates *in vivo*. Of particular note, intratumoral injection of **T4/T7** resulted in tumor regression and enhanced survival in a poorly immunogenic tumor model, non-responsive to traditional immunotherapy. However, this study also has certain limitations: (i) cytokine levels and surface marker expression varied due to donor-to-donor variability; (ii) findings from murine models may not fully translate to alternative murine models, human cells or clinical outcomes; (iii) additional pathways that may contribute to activity were not explored; (iv) *in vivo* cellular immunity, different administration routes, and clinically relevant antigens were not evaluated to further define the adjuvant effects; and (v) tumor immune cell infiltration was not assessed.

In summary, we report the discovery of four synergistic combinations of conjugated innate immune agonists. The TLR1/2/NOD2 combination has demonstrated the ability to elicit a Th17 immune response, while TLR4/TLR7 and TLR7/RIG-I exhibited strong potential for broad applications. These include use as standalone immunotherapeutics or in combination with other immunomodulatory agents for the treatment of tumors and viral infections, as well as adjuvants to enhance vaccine-induced immune responses.

## 4 Materials and methods

### Mice

#### Experiments with Bone-Marrow-Derived Dendritic Cells and T Cells

C57BL/6, OT I (C57BL/6-Tg(TcraTcrb)1100Mjb/J), and OT II (C57BL/6-Tg(TcraTcrb)425Cbn/Crl) mice were obtained from Jackson Laboratory (Bar Harbor, ME) and bred at the University of Leiden (The Netherlands). The mice were housed under standard laboratory conditions with unrestricted access to food and water. Euthanasia was performed under sedation via cervical dislocation. All animal procedures complied with the European Parliament Directive 2010/63/EU and were approved by the Animal Ethics Committee of Leiden University. Details on BMDC culture conditions are provided below.

#### *In Vivo* Vaccination Model

NIH/OlaHsd inbred female mice (2.0–2.5 months old) were bred at the Institute of Immunology, Croatia. During the experiments, they were housed in the institute’s Animal Facility with unrestricted access to food and water. All animal procedures adhered to the Croatian Law on Animal Welfare (2017), in full compliance with the EC Directive 2010/63/EU. Ethical approval was obtained from the Ethical Committee of the Institute of Immunology and the Directorate of Veterinary and Food Safety of the Ministry of Agriculture, Republic of Croatia, approval number: HR-POK-009.

#### *In Vivo* Tumor Model

Female C57Bl/6 mice (9 weeks old) were obtained from Bienta’s Animal Facility. All *in vivo* procedures were approved by Bienta’s Animal Care and Use Committee (BACUC, Approval No. LU-EF-AP-0609624) and conducted in accordance with the European Convention for the Protection of Vertebrate Animals Used for Experimental and Other Scientific Purposes. Mice were housed under specific pathogen-free conditions with a 12-hour light/dark cycle and had unrestricted access to food and water in Bienta’s animal research center laboratory.

### Cell Culture

#### Peripheral Blood Mononuclear Cells

Human PBMCs were isolated from heparinized blood of healthy, consenting donors using density gradient centrifugation with Ficoll-Paque (Pharmacia, Sweden). The cells were washed twice with PBS, resuspended in RPMI 1640 medium (Sigma-Aldrich, St. Louis, MO) supplemented with 10% heat-inactivated fetal bovine serum (Gibco), 2 mM L-glutamine (Sigma-Aldrich), 100 U/mL penicillin (Sigma-Aldrich), and 100 μg/mL streptomycin (Sigma-Aldrich), and subsequently used in assays.

#### Cancer Cell Line

K562, a chronic myelogenous leukemia cell line (Cat. No. CCL-243; ATCC, Manassas, VA) (*129*), was cultured in RPMI 1640 medium (Sigma-Aldrich, St. Louis, MO) supplemented with 10% heat-inactivated fetal bovine serum (Gibco), 2 mM L-glutamine (Sigma-Aldrich), 100 U/mL penicillin (Sigma-Aldrich), and 100 μg/mL streptomycin (Sigma-Aldrich). The cells were incubated in a humidified atmosphere at 37 °C and 5% CO_2_.

#### Bone-Marrow-Derived Dendritic Cells

Bone-marrow cells were harvested from the tibiae of C57BL/6 mice and cultured in RPMI 1640 with HEPES (Lonza, Basel, Switzerland), supplemented with 10% heat-inactivated fetal bovine serum (Lonza), 2 mM L-glutamine (Lonza), 100 U/mL penicillin (Lonza), 100 μg/mL streptomycin (Lonza), 50 μM β-mercaptoethanol (Lonza) and 20 ng/mL granulocyte-macrophage colony-stimulating factor (ImmunoTools, Friesoythe, Germany) for 10 days at 37°C and 5% CO_2_. BMDC purity was assessed via flow cytometry using a PE-labeled anti-mouse CD11c antibody (BioLegend, San Diego, CA), with >90% of cells confirmed as CD11c-positive.

### Cytokine Release from Peripheral Blood Mononuclear Cells

Peripheral blood mononuclear cells were seeded (1 × 10^6^ cells/mL) in 24-well plates in 500 μL of growth medium and treated with conjugated agonists (1 μM), individual agonists (1 μM), unlinked mixtures of agonists (1 μM), lipopolysaccharide (LPS; 1 μg/mL), or the corresponding vehicle (0.1% DMSO). For mechanistic studies of **T4/T7** and **T7/RI**, PBMCs were pre-treated with: (a) TLR4 antagonist TAK242 (5 µM), TLR7 antagonist M5049 (5 µM), or both for 1 h, before the addition of **T4/T7** (100 nM); (b) TLR7 antagonist M5049 (5 µM), RIG-I signaling inhibitor MRT67307 (5 µM), or both for 1 h, before the addition of **T7/RI** (100 nM) or (c) the corresponding vehicle (0.1% DMSO). Cell-free supernatants were collected after 18 h of incubation (37 °C, 5% CO_2_) and stored at −80 °C until tested. Cytokine production was determined with LEGENDplex™ Human Essential Immune Response Panel (Biolegend, San Diego, CA) (comprising IL-4, IL-2, CXCL10 (IP-10), IL-1β, TNF-α, CCL2 (MCP-1), IL-17A, IL-6, IL-10, IFN-γ, IL-12p70, CXCL8 (IL-8), TGF-β1) on an Attune NxT flow cytometer (Thermo Fisher Scientific, Waltham, MA). Standard curves were generated using recombinant cytokines contained in the kit. The data were analyzed using the FlowJo (Tree Star, Inc., Ashland, OR) and Prism (GraphPad, San Diego, CA) software. Statistical significance was determined using one-way ANOVA with post-hoc Bonferroni’s test

### RNA Sequencing

Peripheral blood mononuclear cells from three independent donors were seeded in 24-well plates at a density of 3 × 10^6^ cells/mL in 1 mL of growth medium. Cells were treated with either conjugates (1 μM) or vehicle control (0.1% DMSO) for 18 h at 37 °C with 5% CO_2_. Following treatment, cells were washed with PBS, resuspended in RNAlater RNA stabilization solution (Sigma-Aldrich, St. Louis, MO), and stored at −80 °C for subsequent RNA extraction. RNA extraction, library construction, and sequencing were conducted by Azenta Life Sciences (Leipzig, Germany). Total RNA was extracted using RNeasy mini kit (Qiagen, Hilden, Germany), following the manufacturer’s protocol. RNA samples were quantified using a Qubit 4.0 fluorometer (Life Technologies, Carlsbad, CA), and RNA integrity was verified with RNA kit on a 5300 Fragment Analyzer (Agilent Technologies, Palo Alto, CA), ensuring all samples had an RNA quality number ≥9.4. RNA sequencing libraries were prepared using the NEBNext Ultra II RNA library prep kit for Illumina, according to the manufacturer’s instructions (New England Biolabs, Ipswich, MA). Libraries were loaded on the Illumina NovaSeq 6000 instrument and clustering was performed directly on the NovaSeq before sequencing according to the manufacturer’s instructions. The samples were sequenced using a 2 × 150 paired-end configuration. Image analysis and base calling were conducted using NovaSeq Control Software. Raw sequencing data (.bcl files) generated from the Illumina NovaSeq were converted into fastq format and de-multiplexed using the Illumina bcl2fastq 2.20 software, allowing for one mismatch in index sequence identification. After investigating the quality of the raw data, sequence reads were trimmed to remove possible adapter sequences and nucleotides with poor quality, using Trimmomatic v.0.36. The trimmed reads were mapped to the *Homo sapiens* reference genome, as available on ENSEMBL, using STAR aligner v.2.5.2b, thus generating BAM files. Unique gene hit counts were calculated using feature counts from the Subread package v.1.5.2. Only unique reads that fell within exon regions were counted. After extraction of gene hit counts, the gene hit counts table was used for downstream differential expression analysis.

Differential gene expression analysis was conducted using iDEP 2.0 (*130*). First, low-expressed genes were filtered out (retaining those with ≥0.5 counts per million in at least two libraries). The remaining gene counts were normalized by counts per million in the EdgeR package, with a pseudo count of 4. Differential gene expression analysis was carried out with the DESeq2 method, applying a false discovery rate threshold of <0.1 and a fold-change cutoff of >2 or <0.5. The resulting list of differentially expressed genes was used for gene annotation and pathway enrichment analysis with Metascape (*131*).

### Peripheral Blood Mononuclear Cell Cytotoxicity Assay

The PBMC cytotoxicity assay using K562 cells was performed as described previously, with some modifications (*132*). PBMCs were seeded (4 × 10^5^ cells/well) in duplicate in 96-well U-bottom plates and treated with conjugated agonists (1 μM), individual agonists (1 μM), unlinked mixtures of agonists (1 μM), or vehicle (0.1% DMSO) for 18 h. For mechanistic studies of **T4/T7** and **T7/RI**, PBMCs were pre-treated with: (a) TLR4 antagonist TAK242 (5 µM), TLR7 antagonist M5049 (5 µM), or both for 1 h, before the addition of **T4/T7** (100 nM); (b) TLR7 antagonist M5049 (5 µM), RIG-I signaling inhibitor MRT67307 (5 µM), or both for 1 h, before the addition of **T7/RI** (100 nM) or (c) the corresponding vehicle (0.1% DMSO). K562 cells were stained with CFSE (Invitrogen, Carlsbad, CA), washed twice with complete medium, and added (1 × 10^4^ cells/well) to the pretreated PBMCs for a final effector cell to target tumor cell ratio of 40:1. After a 4 h coincubation (37 °C, 5% CO_2_), the cells were stained with Sytox blue dead cell stain (Invitrogen) and analyzed using an Attune NxT flow cytometer (Thermo Fisher Scientific, Waltham, MA) and the FlowJo software (Tree Star, Inc., Ashland, OR). Cells that were positive for both CFSE and Sytox blue were defined as dead K562 cells. PBMCs alone and CFSE-labeled cancer cells alone were also treated with the compounds at the same concentrations and stained with Sytox blue to exclude any direct cytotoxicity of the compounds toward the PBMCs and cancer cells. Statistical significance was determined using one-way ANOVA with post-hoc Bonferroni’s test.

### Bone-Marrow-Derived Dendritic Cell Antigen Presentation to CD4^+^ and CD8^+^ T cells

CD4^+^ and CD8^+^ T cells were purified from splenocytes of OT II and OT I transgenic mice using CD4^+^ and CD8^+^ T-cell negative selection kits (Miltenyi Biotec, Germany), according to manufacturer instructions. Purified T cells were stained with 0.5 µM CFSE in PBS at 1 x 10^6^ cells/mL (Invitrogen, Carlsbad, CA) and washed. Then, 5 × 10^4^ T cells were cultured in duplicate with 1 × 10^4^ BMDCs/well (pretreated with compounds [1 μM] or LPS [1 μg/mL] and 50 μg/mL ovalbumin (OVA) soluble protein [Invivogen, San Diego, CA] for 18 h, and then washed). After 72 h of coincubation (37 °C, 5% CO_2_), the supernatants were collected and stored at −80 °C for subsequent cytokine measurements. The cells were stained with Fixable viability dye APC-eFluor 780 (eBioscience, Thermo Fisher Scientific, MA), anti-Thy1.2 PE-Cy7 (Biolegend, San Diego, CA), anti-CD8 eFluor450 (eBioscience), anti-CD4 eFluor450 (eBioscience), and anti-CD25 APC antibodies (Biolegend) and analyzed using a Beckman Coulter Cytoflex S flow cytometer (CA) and the FlowJo software (Tree Star, Inc., Ashland, OR). Live Thy1.2^+^/CD4^+^ and Thy1.2^+^/CD8^+^ were evaluated for CFSE dilution and CD25 expression. Supernatants from T cells and BMDC cocultures were collected as described above. The cytokine concentrations were determined with the LEGENDplex MU Th Cytokine Panel (12-plex) (Biolegend, San Diego, CA) (comprising IFN-γ, IL-5, TNF-α, IL-2, IL-6, IL-4, IL-10, IL-9, IL-17A, IL17F, IL-22, IL-13) on an Attune NxT flow cytometer (Thermo Fisher Scientific, Waltham, MA). Standard curves were generated using recombinant cytokines contained in the kit. The data were analyzed using the FlowJo (Tree Star, Inc., Ashland, OR) and Prism (GraphPad, San Diego, CA) software. Statistical significance was determined using one-way ANOVA with post-hoc Dunnett’s test.

### Mouse Immunizations

Sex-matched NIH/OlaHsd mice (six per group) were immunized subcutaneously into the tail base with OVA (10 μg; Serva, Germany) alone or plus the conjugates (42 μg (**T4/T7**), 44 μg (**T7/RI**), 44 μg (**T1/2/T7**), and 64 μg (**T1/2/N2**) – doses are equimolar to 50 μg of **N2/T7**) and Al-hydrogel (100 μg; Invivogen, San Diego, CA) on days 0, 21, and 42. All experimental groups received an injection volume of 0.1 mL per mouse. Seven days after the second booster dose, mice were anesthetized via intraperitoneal administration of ketamine/xylazine (25 mg/kg each) before blood collection from the axillary plexus. Serum samples were individually processed by heat inactivation at 56 °C for 35 minutes to remove complement activity and subsequently stored at −20 °C until analysis.

### Measurement of Ovalbumin-Specific Serum Antibody Concentration

The levels of OVA-specific total IgG, IgG1, and IgG2a in mouse sera were quantified using ELISA. High-binding ELISA plates (Costar) were coated with OVA (1.5 mg/mL; Serva, Germany) in carbonate buffer (pH 9.6) and incubated overnight. To prevent nonspecific binding, plates were blocked with 0.5% (w/v) bovine serum albumin in PBS-T (0.05% [v/v] Tween 20 in PBS) for 2 hours at 37 °C. After washing, five serial dilutions of mouse sera and standard preparations were added in duplicate. Plates were incubated overnight at room temperature, followed by washing and detection using HRP-conjugated goat anti-mouse IgG (Bio-Rad Laboratories) for 2 hours at 37 °C. After further washing, the reaction was developed with 0.6 mg/mL o-phenylenediamine dihydrochloride in citrate-phosphate buffer (pH 5.0) containing 0.5 μL/mL of 30% H₂O₂ for 30 minutes at room temperature in the dark. The reaction was terminated with 12.5% H₂SO₄, and absorbance at 492 nm was measured using a microplate reader (Thermo Fisher Scientific, Waltham, MA).

For OVA-specific IgG1 and IgG2a detection, plates were incubated with biotin-conjugated rat anti-mouse IgG1 or IgG2a (PharMingen, Becton Dickinson) for 2 hours at 37 °C, followed by incubation with streptavidin-peroxidase (Pharmingen) under the same conditions. After washing, the substrate solution was added and incubated for 30 minutes at room temperature in the dark. The reaction was stopped with 12.5% H₂SO₄, and absorbance at 492 nm was recorded. Antibody levels were determined using parallel line assays with standard preparations of anti-OVA IgG (20,000 AU/mL), anti-OVA IgG1 (400,000 AU/mL), and anti-OVA IgG2a (5,000 AU/mL) (*133*). Statistical significance was assessed using one-way ANOVA followed by Dunnett’s multiple comparisons test.

### *In vivo* tumor model

The experiment was conducted following Bienta’s standard operating procedures. B16F10 cells (ATCC: CRL-6475) were cultured in DMEM (4.5 g/L glucose) supplemented with 10% FBS, 100 U/mL penicillin, and 100 μg/mL streptomycin at 37°C in a 5% CO₂ atmosphere. Cells were harvested using a 0.05% trypsin-EDTA solution, centrifuged, and resuspended in serum-free DMEM. Cell count and viability were assessed using a hemocytometer and trypan blue exclusion test. The final cell suspension, containing 5×10⁵ cells per 50 μL in DPBS, was kept on ice before injection. Each of the 90 female C57BL/6 mice received a subcutaneous inoculation of 5×10⁵ B16F10 cells (50 μL) above the upper thigh. Mice were randomized into four groups based on tumor volume and body weight between days 13 and 23 post-inoculation. Treatment was initiated when tumor nodules reached an average volume of approximately 100 mm³. Mice with tumors outside the predefined volume range were excluded from the study.

Mice received intratumoral injections of vehicle (10% DMSO in PBS) or the compounds **T4/T7**, **T7/RI**, and resiquimod (formulated in 10% DMSO in PBS) at doses of 49 μg, 52 μg, and 20 μg, respectively. Injections were administered into a single tumor site at a volume equivalent to 1/10 of the tumor volume, every two to three days, for a total of six injections over two weeks. Animals were weighed upon arrival, on the day of B16F10 cell inoculation, at each treatment session, and daily throughout the treatment and observation period. Body weight changes were expressed as a percentage relative to the initial weight (DPT 0) for each animal. Tumor size was measured every two days using a digital caliper, with two perpendicular diameters recorded: length (L, the larger diameter) and width (W, the smaller diameter). Tumor volume (V) was calculated using the formula: V = (W² × L) × 0.5 mm³. Mice were euthanized if tumor volumes exceeded 3000 mm³ or if tumor sites developed ulcerations. Euthanasia was performed via CO₂ inhalation followed by cervical dislocation, in accordance with Bienta’s SOP, by trained personnel. The relative tumor volume (RTV) was calculated using the formula: RTV = (tumor volume on measured day) / (tumor volume on day 0). Tumor growth inhibition (TGI, %) was determined using: TGI (%) = [1 − (RTV of treated group) / (RTV of control group)] × 100. Alternatively, inhibition of tumor growth was calculated as the percentage of tumor volume in the treated group relative to the control group (T/C, %). Euthanized animals were classified as deceased in the survival analysis.

### Statistics

Data analysis was performed using the Prism software (version 10; GraphPad Software, CA). Statistical differences were determined as specified in individual experimental procedures above and in the corresponding Figure captions.

## 6 Acknowledgments

### General

/

### Funding

This work was supported by grants from Slovenian Research Agency ARIS to Ž.J. (P1-0420, J3-2517, J3-4496).

### Author contributions

Conceptualization: Ž.J.; Experimentation: Š.J., L.B., M.Š., S.P.; Data analysis: Ž.J., Š.J., S.G., R.F., B.S., S.P.; Visualization: Š.J., S.G., Ž.J.; Funding acquisition: Ž.J.; Project administration: Ž.J.; Supervision: Ž.J., B.S., R.F.; Writing – original draft: Ž.J., Š.J., S.G.; Writing – review & editing: Ž.J., Š.J., S.G., R.F., M.Š., B.S., S.P.

### Data and materials availability

The data supporting the findings of this study are available within the article [and/or] its supplementary materials.

## 7 Supplementary materials

Supporting materials and methods; Supplementary Figures; Supplementary Schemes; Supplementary Tables; HPLC traces of final compounds; HRMS spectra of final compounds; 1H and 13C NMR spectra of final compounds.

