## Supplementary Information for "Probing immune signatures of conjugated pattern recognition receptor ligands identifies chimeras with adjuvant and antitumor activity"

Žiga Jakopin

### 1. Supplementary materials and methods

#### 1.1 Synthesis of conjugates

Chemicals were obtained from Sigma-Aldrich (St. Louis, MO, U.S.A.), Tokyo Chemical Industry (Tokyo, Japan), Acros Organics (Geel, Belgium), Enamine (Monmouth Junction, NJ, U.S.A.), and Apollo (Stockport, U.K.) and were used without further purification. LPS (from E. coli O55:B5), TAK242, M5049 and MRT67307 were obtained from InvivoGen (San Diego, CA, U.S.A.). Compound **N2/T7** (Ethyl $N^5$-(2-(2-(4-((6-amino-2-butoxy-8-hydroxy-9*H*-purin-9-yl)methyl)benzamido)ethoxy)ethyl)-$N^2$-((*E*)-3-(4-hydroxy-3-methoxyphenyl)acryloyl)glycyl-L-valyl-D-glutaminate) was prepared as previously described (*1*). Compounds **N2** (*2*), **T1/2** (*3*), **T4** (*4*), **T7** (*5, 6*) and **RI** (*7*) were prepared according to the reported procedures. Analytical TLC was performed on Merck 60 F254 silica gel plates (0.25 mm), with visualization using ultraviolet light, ninhydrin, and potassium permanganate. Flash column chromatography was carried out on Merck silica gel 60 (particle size 240−400 mesh) and on Biotage Isolera One Flash Chromatograph using Biotage® Sfär C18 D Duo 100 Å 30 µm 30 g. 1H and 13C NMR spectra were recorded at 400 and 100 MHz, respectively, on an Avance III spectrometer (Bruker Corporation, Billerica, MA, U.S.A.) in $CDCl_3$ or DMSO-$d_6$ (MeOD) with tetramethylsilane as the internal standard. Mass spectra were obtained using an Exactive Plus orbitrap mass spectrometer (Thermo Fisher Scientific, Waltham, MA, U.S.A.) or on Expression CMS mass spectrometer (Advion Inc., Ithaca, NY, U.S.A.). Analytical UHPLC analyses were performed on a Dionex UltiMate 3000 Rapid Separation Binary System (Thermo Fisher Scientific, Waltham, MA, U.S.A.) equipped with an autosampler, a binary pump system, a photodiode array detector, a thermostatted column compartment, and the Chromeleon Chromatography data system. The columns used were Waters Acquity UPLC BEH C18 (1.7 µm,

2.1 mm × 50 mm) or Waters Acquity UPLC CSH C18 (1.7 µm, 2.1 mm × 50 mm) with a flow rate of 0.3 mL/min. The eluent was a mixture of 0.1% TFA in water (A) and MeCN (B) with a gradient (%B) as follows: 0−10 min, 5−95%; 10−12 min, 95%; 12−12.5 min, 95−5%. For compound T7/RI the mobile phase consisted of 0.1% TFA in water (A) and MeCN (B), employing the following gradient: 95% A to 5% A in 7 min, then 95% B for 1 min, with flow rate of 0.3 mL/min. The columns were thermostatted at 40 °C. All the compounds tested were established to be ≥95% pure. The analytical data here were identical to those reported previously. The assembly of the final compounds was as described below.

##### 1.1.1 General synthetic procedures

###### 1.1.1.1 General Procedure A: Boc Deprotection

The tert-butyloxycarbonyl (Boc)-protected compound was added to an ice-chilled mixture of trifluoroacetic acid (TFA) and dichloromethane (DCM) (1:5), and the mixture was allowed to warm to room temperature. After 3 h, the solvent was evaporated *in vacuo*. The residue was washed three times with diethyl ether.

###### 1.1.1.2 General Procedure B: COMU Mediated Coupling

To an ice-chilled solution of the amine or alcohol (1 – 1.2 equiv) and carboxylic acid (1 – 1.2 equiv) in anhydrous dimethylformamide (DMF), *N*,*N*-diisopropylethylamine (DIPEA; 4 equiv) and 1-[(1-(cyano-2-ethoxy-2-oxoethylideneaminooxy)-dimethylamino-morpholinomethylene)]- methanaminium hexafluorophosphate (COMU; 1.2 equiv) were added, and the mixture was allowed to warm to room temperature. The stirring was continued overnight, after which the mixture was washed twice with saturated $NaHCO_3$, and once with brine. The organic layer was dried over anhydrous $Na_2SO_4$ and concentrated *in vacuo*.

###### 1.1.1.3 General Procedure C: HATU Mediated Coupling

To an ice-chilled solution of the amine or alcohol (1 – 1.5 equiv) and carboxylic acid (1 – 1.5 equiv) in anhydrous dimethylformamide (DMF), *N*,*N*-diisopropylethylamine (DIPEA; 4 equiv) and 1-[bis(dimethylamino)methylene]-1H-1,2,3- triazolo[4,5-b]pyridinium 3-oxide hexafluorophosphate (HATU; 1.5 equiv) were added, and the mixture was allowed to warm to room temperature. The stirring was continued overnight, after which the mixture was washed twice with 1 M HCl and saturated $NaHCO_3$, and once with brine. The organic layer was dried over anhydrous $Na_2SO_4$ and concentrated *in vacuo*.

##### 1.1.2 Compound characterization

*Tert-butyl (2-(2-(4-((6-amino-2-butoxy-8-hydroxy-9H-purin-9-yl)-methyl)benzamido)ethoxy)ethyl)carbamate (**1**).*

Tert-butyl (2-(2-aminoethoxy)ethyl)carbamate (380 mg, 1.86 mmol) and DIPEA (0.523 mL, 2.99 mmol) were dissolved in DCM (0.5 mL) and added to the stirring solution of compound T7 (200 mg, 0.56 mmol) in DMSO (3 mL). To an ice-chilled solution COMU (0.640 g, 1.49 mmol) was added, and the mixture was stirred at room temperature for 3 h. Subsequently, ethyl acetate (40 mL) was added and the mixture was cooled in ice for 1 h, after which 1 M $NaHCO_3$ (40 mL) was added. The precipitate obtained, after concentrating the mixture *in vacuo,* was filtered and washed with water and diethyl ether to give compound **1** as an off-white solid. Yield (160 mg, 53%). 1H NMR (400 MHz, DMSO-*d*6) $\delta$ = 9.98 (s, 1H), 8.44 (t, *J* = 6.4, 1H), 7.78 (d, *J* = 8.2, 2H), 7.34 (d, *J* = 8.2, 2H), 6.76 (t, *J* = 6.8, 1H), 6.47 (s, 2H), 4.90 (s, 2H), 4.12 (t, *J* =

7.0, 2H), 3.53−3.45 (m, 2H), 3.43−3.36 (m, 4H), 3.14 − 2.97 (m, 2H), 1.69−1.52 (m, 2H), 1.42−1.27 (m, 11H), 0.89 (t, *J* = 7.4, 3H).

*4-((6-amino-2-butoxy-8-hydroxy-9H-purin-9-yl)methyl)-N-(2-(2-(2-((4-oxo-3-phenyl-4,5-dihydro-3H-pyrimido[5,4-b]indol-2-yl)thio)acetamido)ethoxy)ethyl)benzamide (**T4/T7**).*

Compound **1** (89 mg, 0.164 mmol, 1.1 eq) was deprotected using General procedure A and coupled to **T4** (52 mg, 0.149 mmol, 1 eq) using General procedure B. The crude product was purified by Isolera One flash chromatography (acetonitrile/0.1% TFA 20% → 100%) to give compound **T4/T7** as a pale-yellow solid (38 mg, 10 %). $^{1}H$ NMR (400 MHz, DMSO-$d_6$) δ 12.10 (s, 1H), 9.98 (s, 1H), 8.44 (t, *J* = 5.5 Hz, 1H), 8.32 (t, *J* = 5.5 Hz, 1H), 8.06 – 7.99 (m, 1H), 7.82 – 7.74 (m, 2H), 7.64 – 7.55 (m, 3H), 7.53 – 7.42 (m, 4H), 7.39 – 7.30 (m, 2H), 7.27 – 7.19 (m, 1H), 6.47 (s, 2H), 4.88 (s, 2H), 4.12 (t, *J* = 6.6 Hz, 2H), 3.91 (s, 2H), 3.54 – 3.34 (m, 5H), 3.31 – 3.18 (m, 2H), 1.67 – 1.51 (m, 2H), 1.43 – 1.31 (m, 2H), 0.88 (t, *J* = 7.4 Hz, 3H). $^{13}C$ NMR (100 MHz, DMSO) δ 165.80, 164.73, 158.79, 153.63, 151.07, 150.89, 147.82, 146.49, 138.88, 137.60, 135.92, 134.74, 132.22, 128.55, 128.24, 126.11, 125.98, 125.88, 119.07, 119.00, 118.84, 117.94, 111.50, 96.95, 67.45, 67.42, 64.52, 40.78, 35.14, 29.25, 17.41, 12.39. HRMS *m/z* calculated for $C_{39}H_{41}O_6N_{10}S$: 777.2926 $(M + H)^+$, found 777.2903.

*Tert-Butyl (2-(2-(2-(4-((6-amino-2-butoxy-8-hydroxy-9H-purin-9-yl)methyl)benzamido)ethoxy)ethoxy)ethyl)carbamate (**2**).*

SG169 (1.157 g, 4.659 mmol, 3.33 eq) and DIPEA (1.299 mL, 7.457 mmol, 5.33 eq) were dissolved in DCM (1 mL) and added to the stirring solution of compound **T7** (500 mg, 1.399 mmol, 1 eq) in DMSO (20 mL). After cooling the reaction mixture in ice, COMU (1.594 g, 3.722

mmol, 2.66 eq) was added and the mixture was stirred at rt for 20 h. Ethyl acetate (100 mL) and 1M $NaHCO_3$ solution (75 mL) were added. After concentrating the mixture *in vacuo*, the mixture was cooled in ice for one hour. The precipitate was filtered and washed with water and diethyl ether to afford the subject compound SB57. Yield (450 mg, 55%). $^1$H NMR (400 MHz, DMSO-$d_6$) δ 10.14 (s, 1H), 8.48 (t, *J* = 5.6 Hz, 1H), 7.84 – 7.76 (m, 2H), 7.39 – 7.30 (m, 2H), 6.84 – 6.68 (m, 1H), 6.52 (s, 2H), 4.90 (s, 2H), 4.12 (t, *J* = 6.6 Hz, 2H), 3.61 – 3.45 (m, 8H), 3.45 – 3.38 (m, 2H), 3.07 – 3.02 (m, 2H), 1.69 – 1.53 (m, 2H), 1.37 – 1.29 (m, 11H), 0.89 (t, *J* = 7.4 Hz, 3H).

*2-(2-naphthamido)-N-(2-(2-(2-(4-((6-amino-2-butoxy-8-hydroxy-9H-purin-9-yl)methyl)benzamido)ethoxy)ethoxy)ethyl)benzo[d]thiazole-6-carboxamide (**T7/RI**).*

Compound **2** (120 mg, 0.205 mmol, 1 eq) was deprotected using general procedure A and coupled to **RI** (79 mg, 0.226 mmol, 1.1 eq) using general procedure B. The crude product was purified by flash chromatography (dichloromethane:methanol 25:1) to give compound **T7/RI** as a white solid (18 mg, 11%). $^1$H NMR (400 MHz, DMSO-$d_6$) δ 13.27 (s, 1H), 10.05 (s, 1H), 8.93 (s, 1H), 8.70 – 8.64 (m, 1H), 8.64 – 8.59 (m, 1H), 8.59 – 8.53 (m, 1H), 8.26 – 8.21 (m, 1H), 8.20 – 8.15 (m, 2H), 8.14 – 8.09 (m, 1H), 8.05 – 8.01 (m, 1H), 7.97 – 7.89 (m, 1H), 7.88 – 7.82 (m, 2H), 7.79 – 7.71 (m, 2H), 7.41 (d, *J* = 8.3 Hz, 2H), 6.53 (s, 2H), 4.96 (s, 2H), 4.18 (t, *J* = 6.6 Hz, 2H), 3.62 (s, 12H), 1.76 – 1.59 (m, 2H), 1.42 (q, *J* = 7.4 Hz, 2H), 0.95 (t, *J* = 7.4 Hz, 3H). $^{13}$C NMR (100 MHz, DMSO) δ 166.41, 166.39, 152.40, 149.26, 140.32, 135.34, 134.10, 132.44, 131.84, 130.37, 130.11, 129.80, 129.09, 128.81, 128.21, 127.91, 127.78, 127.58, 126.01, 124.79, 121.73, 120.26, 98.68, 70.07, 69.43, 69.39, 67.33, 62.48, 42.77, 30.80, 25.96, 19.08, 14.10. HRMS *m/z* calculated for $C_{42}H_{44}O_7N_9S$: 818.3079 $(M + H)^+$, found 818.3069.

*Ethyl 6-(2-((4-oxo-3-phenyl-4,5-dihydro-3H-pyrimido[5,4-b]indol-2-yl)thio)acetamido)hexanoate (**3**).*

Synthesized from SG44 (167 mg, 0.854 mmol, 1.5 eq) and **T4** (200 mg, 0.569 mmol, 1 eq) using General procedure C to give compound SB22 as an off-white solid. Yield (150 mg, 53%). $^1$H NMR (400 MHz, DMSO-$d_6$) δ 12.11 (s, 1H), 8.26 – 8.17 (m, 1H), 8.08 – 7.99 (m, 1H), 7.63 – 7.57 (m, 3H), 7.53 – 7.42 (m, 4H), 7.28 – 7.20 (m, 1H), 4.01 (q, *J* = 7.1 Hz, 2H), 3.88 (s, 2H), 3.05 (q, *J* = 6.6 Hz, 2H), 2.11 (t, *J* = 6.9, 6.5 Hz, 2H), 1.44 – 1.36 (m, 3H), 1.28 – 1.20 (m, 5H), 1.15 (t, *J* = 7.1 Hz, 3H).

*6-(2-((4-oxo-3-phenyl-4,5-dihydro-3H-pyrimido[5,4-b]indol-2-yl)thio)acetamido)hexanoic acid (**4**).*

To a stirring solution of compound **3** (150 mg, 0.303 mmol) in methanol (5 mL) 1M NaOH (3 mL) was added. The mixture was stirred at room temperature overnight. The next day $H_2O$ (20 mL) was added and methanol was evaporated *in vacuo.* Water phase was washed with ethyl acetate (20 mL) and acidified with 1M HCl to pH 3. The product was extracted two times with ethyl acetate (20 mL). Combined organic phases were washed with brine (10 mL), dried over anhydrous $Na_2SO_4$ and concentrated *in vacuo* to give compound SB27 as an off-white solid. Yield (141 mg, 100%). $^1$H NMR (400 MHz, DMSO-$d_6$) δ 12.10 (s, 1H), 11.97 (s, 1H), 8.22 (t, *J* = 5.6 Hz, 1H), 8.11 – 8.00 (m, 1H), 7.62 – 7.59 (m, 3H), 7.54 – 7.39 (m, 4H), 7.33 – 7.18 (m, 1H), 3.88 (s, 2H), 3.05 (q, *J* = 6.6 Hz, 2H), 2.08 (t, *J* = 7.4 Hz, 2H), 1.42 – 1.37 (m, 2H), 1.25 – 1.22 (m, 4H).

*Dicyclopentyl ((E)-3-(3-methoxy-4-((6-(2-((4-oxo-3-phenyl-4,5-dihydro-3H-pyrimido[5,4-b]indol-2-yl)thio)acetamido)hexanoyl)oxy)phenyl)acryloyl)glycyl-L-valyl-D-glutamate (**T4/N2**).*

Synthesized from **4** (141 mg, 0.303 mmol, 1.2 eq) and **N2** (156 mg, 0.253 mmol, 1 eq) using General procedure B. The crude product was purified flash $^1$H NMR (400 MHz, Chloroform-*d*) δ 10.65 (s, 1H), 8.05 (d, *J* = 8.0 Hz, 1H), 7.59 – 7.45 (m, 8H), 7.40 – 7.32 (m, 4H), 7.22 – 7.16 (m, 1H), 7.02 – 6.97 (m, 2H), 6.89 – 6.85 (m, 1H), 6.39 (d, *J* = 15.6 Hz, 1H), 5.17 – 5.10 (m, 2H), 4.51 – 4.41 (m, 2H), 4.14 – 4.05 (m, 2H), 3.88 – 3.82 (m, 2H), 3.72 (s, 3H), 3.29 (q, *J* = 6.5 Hz, 2H), 2.37 – 2.29 (m, 4H), 2.15 – 2.10 (m, 1H), 1.97 (s, 4H), 1.84 – 1.75 (m, 5H), 1.69 – 1.51 (m, 19H), 1.38 – 1.29 (m, 2H), 0.93 – 0.87 (m, 6H). $^{13}$C NMR (100 MHz, $CDCl_3$) δ 172.60, 171.58, 171.38, 171.28, 169.63, 168.76, 166.51, 155.85, 153.30, 151.14, 140.86, 140.72, 139.47, 138.20, 135.48, 133.61, 130.39, 129.90, 129.88, 129.15, 129.13, 128.19, 123.00, 121.01, 120.59, 120.47, 120.35, 120.22, 119.34, 113.11, 111.64, 78.67, 77.51, 77.24, 58.62, 55.81, 51.99, 43.48, 39.54, 36.07, 33.51, 32.68, 32.63, 32.57, 32.49, 30.96, 30.72, 30.64, 28.93, 26.86, 26.10, 24.32, 23.68, 23.63, 19.32, 17.89. HRMS *m/z* calculated for $C_{56}H_{68}O_{12}N_7S$: 1062.4641 $(M + H)^+$, found 1062.4635.

*2-(1-(2-(methylamino)-5-nitrophenyl)-1H-imidazol-4-yl)-5-(trifluoromethyl)phenyl 6-((tert-butoxycarbonyl)amino)hexanoate (**5**).*

Synthesized from **T1/2** (50 mg, 0.134 mmol, 1 eq) and SB41 (37 mg, 0.161 mmol, 1.2 eq) using General procedure C (1M HCl was not used for extraction). The crude product was purified by flash chromatography (hexane:ethyl acetate 1:1) to give compound SB50 as a white solid (35 mg, 44%). $^1$H NMR (400 MHz, ) δ 8.32 (d, *J* = 8.2 Hz, 1H), 8.22 (dd, *J* = 9.3, 2.7 Hz, 1H), 8.04 – 7.97 (m, 2H), 7.86 – 7.75 (m, 1H), 7.75 – 7.67 (m, 1H), 7.62 (d, *J* = 1.9 Hz, 1H), 6.86 (d, *J* = 9.4

Hz, 1H), 6.82 – 6.68 (m, 2H), 2.90 – 2.79 (m, 5H), 2.76 – 2.67 (m, 2H), 1.71 – 1.55 (m, 2H), 1.47 – 1.20 (m, 13H).

*2-(1-(2-(methylamino)-5-nitrophenyl)-1H-imidazol-4-yl)-5-(trifluoromethyl)phenyl 6-(2-((4-oxo-3-phenyl-4,5-dihydro-3H-pyrimido[5,4-b]indol-2-yl)thio)acetamido)hexanoate (**T1/2/T4**).*
Compound **5** (35 mg, 0.0594 mmol, 1 eq) was deprotected using general procedure A and coupled to **T4** (23 mg, 0.0653 mmol, 1.1 eq) using general procedure C (1M HCl was not used for extraction). The crude product was purified by flash chromatography (hexane:ethyl acetate 1:3) to give compound **T1/2/4** as a white solid (21 mg, 36%). $^{1}H$ NMR (400 MHz, DMSO-$d_6$) δ 12.16 (s, 1H), 8.37 (d, *J* = 8.2 Hz, 1H), 8.29 – 8.21 (m, 2H), 8.12 – 8.03 (m, 3H), 7.82 (d, *J* = 1.2 Hz, 1H), 7.75 (dd, *J* = 8.4, 1.9 Hz, 1H), 7.70 – 7.60 (m, 4H), 7.57 – 7.46 (m, 4H), 7.30 – 7.25 (m, 1H), 6.87 (d, *J* = 9.4 Hz, 1H), 6.83 – 6.74 (m, 1H), 3.93 (s, 2H), 3.10 (q, *J* = 6.7 Hz, 2H), 2.83 (d, *J* = 4.7 Hz, 3H), 2.64 (t, *J* = 7.5 Hz, 2H), 1.66 – 1.56 (m, 2H), 1.50 – 1.41 (m, 2H), 1.38 – 1.30 (m, 2H). $^{13}C$ NMR (100 MHz, DMSO) δ 171.93, 167.16, 155.42, 152.89, 150.73, 146.98, 139.40, 139.13, 137.70, 136.52, 135.92, 135.55, 131.09, 130.31, 130.01, 128.89, 127.71, 127.12, 124.32, 123.12, 123.08, 122.97, 121.51, 121.39, 121.27, 120.85, 120.76, 120.52, 119.73, 113.29, 110.48, 55.38, 39.22, 36.99, 33.99, 30.22, 29.27, 26.20, 24.15. HRMS *m/z* calculated for $C_{41}H_{36}O_6N_8F_3S$: 825.2425 $(M + H)^+$, found 825.2411.

*4-((6-(2-(1-(2-(methylamino)-5-nitrophenyl)-1H-imidazol-4-yl)-5-(trifluoromethyl)phenoxy)-6-oxohexyl)amino)-4-oxobutanoic acid (**6**).*
Compound **5** (110 mg, 0.187 mmol, 1 eq) was deprotected using general procedure A. The resulting intermediate was dissolved in DMF (1 mL). To an ice-chilled solution succinic anhydride (22 mg, 0.224 mmol, 1.2 eq), $Et_3N$ (101 µL, 0.746 mmol, 4 eq) and DMAP (catalytic

amount) were added and stirring continued overnight on room temperature. Subsequently DCM (20 mL) was added and the mixture was extracted twice with saturated NaHCO3. Combined water phases were acidified with 1 M HCl to pH 5. The product was extracted with ethyl acetate, dried over anhydrous $Na_2SO_4$ and concentrated *in vacuo to* produce the compound SB60. Yield (100 mg, 89%). $^1$H NMR (400 MHz, DMSO-$d_6$) δ 12.05 (s, 1H), 8.35 – 8.29 (m, 1H), 8.23 (dd, *J* = 9.3, 2.6 Hz, 1H), 8.05 – 7.99 (m, 2H), 7.82 – 7.73 (m, 2H), 7.73 – 7.67 (m, 1H), 7.66 – 7.58 (m, 1H), 6.91 – 6.82 (m, 1H), 6.81 – 6.71 (m, 1H), 2.98 (q, *J* = 6.4 Hz, 2H), 2.81 (d, *J* = 4.8 Hz, 3H), 2.72 – 2.66 (m, 2H), 2.43 – 2.34 (m, 2H), 2.27 (t, *J* = 6.8 Hz, 2H), 1.69 – 1.55 (m, 2H), 1.43 – 1.26 (m, 4H).

*Dicyclopentyl ((E)-3-(3-methoxy-4-((4-((6-(2-(1-(2-(methylamino)-5-nitrophenyl)-1H-imidazol-4-yl)-5-(trifluoromethyl)phenoxy)-6-oxohexyl)amino)-4-oxobutanoyl)oxy)phenyl)acryloyl)glycyl-L-valyl-D-glutamate (**T1/2/N2**).*

Synthesized from **6** (100 mg, 0.166 mmol, 1.1 eq) and N2 (93 mg, 0.151 mmol, 1.0 eq) using general procedure B. The crude product was purified by Isolera One flash chromatography (acetonitrile/0.1% TFA 20% → 100%) to obtain a compound **T1/2/N2** as a pale-yellow solid (25 mg, 14 %). $^1$H NMR (400 MHz, DMSO-$d_6$) δ 8.42 – 8.26 (m, 3H), 8.22 (dd, *J* = 9.3, 2.7 Hz, 1H), 8.02 (dd, *J* = 6.1, 2.0 Hz, 2H), 7.96 (d, *J* = 9.0 Hz, 1H), 7.88 (t, *J* = 5.6 Hz, 1H), 7.80 (d, *J* = 1.3 Hz, 1H), 7.74 – 7.66 (m, 1H), 7.63 (d, *J* = 1.8 Hz, 1H), 7.42 (d, *J* = 15.7 Hz, 1H), 7.33 (d, *J* = 1.8 Hz, 1H), 7.17 (dd, *J* = 8.3, 1.8 Hz, 1H), 7.09 (d, *J* = 8.2 Hz, 1H), 6.86 (d, *J* = 9.4 Hz, 1H), 6.80 – 6.70 (m, 2H), 5.11 – 4.94 (m, 2H), 4.32 – 4.15 (m, 2H), 3.91 (d, *J* = 5.6 Hz, 2H), 3.80 (s, 3H), 3.02 (q, *J* = 6.5 Hz, 2H), 2.81 (d, *J* = 4.8 Hz, 3H), 2.76 – 2.67 (m, 3H), 2.43 (t, *J* = 7.1 Hz, 2H), 2.35 – 2.27 (m, 2H), 2.04 – 1.89 (m, 2H), 1.79 (d, *J* = 8.2 Hz, 5H), 1.69 – 1.46 (m, 14H), 1.42 – 1.27 (m, 4H), 0.85 (t, *J* = 6.3 Hz, 6H). $^{13}$C NMR (100 MHz, DMSO) δ 172.18, 172.03, 171.77, 171.47,

171.13, 170.60, 169.20, 165.66, 161.28, 159.60, 151.52, 150.76, 146.99, 140.77, 140.70, 139.16, 138.88, 135.93, 135.55, 134.26, 131.10, 128.88, 124.35, 123.72, 123.09, 121.56, 121.54, 121.41, 120.56, 112.14, 112.11, 110.52, 108.02, 77.67, 76.93, 57.82, 56.27, 51.62, 38.76, 34.07, 32.62, 32.59, 32.48, 31.29, 30.41, 30.35, 30.23, 29.27, 26.37, 26.23, 24.19, 23.71, 23.66, 19.56, 18.33. HRMS *m/z* calculated for $C_{59}H_{72}O_{15}N_8F_3$: 1189.5064 $(M + H)^+$, found 1189.5062.

*2-(1-(2-(methylamino)-5-nitrophenyl)-1H-imidazol-4-yl)-5-(trifluoromethyl)phenyl 6-(4-((6-amino-2-butoxy-8-hydroxy-9H-purin-9-yl)methyl)benzamido)hexanoate (**T1/2/T7**).*

Compound **5** (95 mg, 0.161 mmol, 1 eq) was deprotected using general procedure A and coupled to **T7** (69 mg, 0.193 mmol, 1.2 eq) using general procedure B. The crude product was purified by flash chromatography (dichloromethane:methanol 15:1) to give compound **T1/2/T7** as a white solid (39 mg, 29%). $^1$H NMR (400 MHz, DMSO-$d_6$) δ 9.98 (s, 1H), 8.41 – 8.34 (m, 1H), 8.34 – 8.28 (m, 1H), 8.21 – 8.14 (m, 1H), 8.03 – 7.97 (m, 2H), 7.80 (d, *J* = 1.3 Hz, 1H), 7.77 – 7.72 (m, 2H), 7.69 (d, *J* = 8.9 Hz, 1H), 7.64 – 7.59 (m, 1H), 7.33 (d, *J* = 8.2 Hz, 2H), 6.79 (d, *J* = 9.4 Hz, 1H), 6.77 – 6.72 (m, 1H), 6.47 (s, 1H), 4.90 (s, 2H), 4.12 (t, *J* = 6.6 Hz, 2H), 2.77 (d, *J* = 4.8 Hz, 3H), 1.62 (m, 4H), 1.56 – 1.44 (m, 2H), 1.44 – 1.31 (m, 4H), 1.29 – 1.19 (m, 5H), 0.89 (t, *J* = 7.4 Hz, 3H). $^{13}$C NMR (100 MHz, DMSO) δ 172.02, 166.19, 160.57, 152.69, 150.72, 149.60, 148.29, 146.99, 140.51, 139.13, 135.93, 135.54, 130.03, 128.86, 127.84, 127.81, 127.79, 127.64, 127.11, 124.31, 121.54, 121.40, 110.44, 98.75, 66.30, 42.58, 34.07, 31.04, 30.19, 29.31, 29.24, 26.26, 24.21, 19.20, 14.16. HRMS *m/z* calculated for $C_{40}H_{42}F_3O_7N_{10}$: 831.3185 $(M + H)^+$, found 831.3186.

*Tert-butyl (2-(2-(2-(2-(2-naphthamido)benzo[d]thiazole-6-carboxamido)ethoxy)ethoxy)ethyl)carbamate (**7**).*

Synthesized from **RI** (120 mg, 0.344 mmol, 1 eq) and SG169 (155 mg, 0.447 mmol, 1.3 eq) using general procedure B. Yield (160 mg, 69%) $^{1}H$ NMR (400 MHz, DMSO-$d_6$) δ 13.21 (s, 1H), 8.93 – 8.81 (m, 1H), 8.60 (t, *J* = 5.6 Hz, 1H), 8.56 – 8.45 (m, 1H), 8.22 – 8.15 (m, 1H), 8.15 – 8.07 (m, 2H), 8.07 – 8.01 (m, 1H), 7.97 (dd, *J* = 8.4, 1.8 Hz, 1H), 7.89 – 7.78 (m, 1H), 7.77 – 7.56 (m, 2H), 6.78 (s, 1H), 3.63 – 3.35 (m, 10H), 3.06 (q, *J* = 6.0 Hz, 2H), 1.36 (s, 9H).

*2-(2-naphthamido)-N-(2-(2-(2-(2-((4-oxo-3-phenyl-4,5-dihydro-3H-pyrimido[5,4-b]indol-2-yl)thio)acetamido)ethoxy)ethoxy)ethyl)benzo[d]thiazole-6-carboxamide (**T4/RI**).*

Compound **7** (80 mg, 0.138 mmol, 1 eq) was deprotected using general procedure A and coupled to SB19 (53 mg, 0.152 mmol, 1.2 eq) using general procedure B. The crude product was purified by flash chromatography (dichloromethane:methanol 40:1) to give compound SB81 as a brown solid (55 mg, 49%). $^{1}H$ NMR (400 MHz, DMSO-$d_6$) δ 13.20 (s, 1H), 12.10 (s, 1H), 8.91 – 8.80 (m, 1H), 8.59 (t, *J* = 5.4 Hz, 1H), 8.53 (s, 1H), 8.32 (t, *J* = 5.7 Hz, 1H), 8.20 – 8.13 (m, 1H), 8.13 – 8.07 (m, 2H), 8.07 – 8.00 (m, 2H), 8.00 – 7.91 (m, 1H), 7.89 – 7.81 (m, 1H), 7.75 – 7.63 (m, 2H), 7.63 – 7.54 (m, 3H), 7.54 – 7.39 (m, 4H), 7.28 – 7.19 (m, 1H), 3.91 (s, 2H), 3.56 – 3.39 (m, 10H), 3.24 (q, *J* = 5.8 Hz, 2H). $^{13}C$ NMR (100 MHz, DMSO) δ 167.60, 166.41, 155.42, 152.82, 139.39, 137.70, 136.51, 135.34, 132.44, 130.33, 130.10, 130.01, 129.81, 129.10, 128.81, 128.22, 127.77, 127.59, 126.00, 124.79, 121.71, 120.86, 120.80, 120.62, 119.72, 113.29, 70.05, 69.95, 69.46, 69.42, 55.39, 36.89. HRMS *m/z* calculated for $C_{43}H_{38}O_6N_7S_2$: 812.2320 $(M + H)^+$, found 812.2312.

*Ethyl 6-(2-(2-naphthamido)benzo[d]thiazole-6-carboxamido)hexanoate (**8**).*

Synthesized from RI (900 mg, 2.583 mmol, 1 eq) and SG44 (607 mg, 3.100 mmol, 1.2 eq) using general procedure B. Yield (0.935 mg, 75%). $^1$H NMR (400 MHz, DMSO-$d_6$) δ 8.88 – 8.80 (m, 1H), 8.54 (t, *J* = 5.5 Hz, 1H), 8.50 – 8.43 (m, 1H), 8.20 (dd, *J* = 8.6, 1.8 Hz, 1H), 8.14 – 7.99 (m, 3H), 7.93 (dd, *J* = 8.5, 1.8 Hz, 1H), 7.81 – 7.74 (m, 1H), 7.71 – 7.57 (m, 2H), 4.11 – 3.97 (m, 2H), 3.32 – 3.18 (m, 2H), 2.35 – 2.23 (m, 2H), 1.68 – 1.45 (m, 4H), 1.45 – 1.20 (m, 2H), 1.20 – 1.11 (m, 3H).

*6-(2-(2-naphthamido)benzo[d]thiazole-6-carboxamido)hexanoic acid (**9**).*

To a stirring solution of compound **8** (0.935 mg, 1.912 mmol, 1 eq) in methanol (15 mL) 1M NaOH (15 mL) was added. The mixture was stirred at room temperature for 2.5 h. Subsequently methanol was evaporated *in vacuo* and water phase was washed with ethyl acetate (30 mL) and acidified with 1M HCl to pH 5. The product was extracted two times with ethyl acetate (20 mL). Yield (454 mg, 52%). $^1$H NMR (400 MHz, DMSO-$d_6$) δ 8.73 (s, 1H), 8.36 – 8.28 (m, 2H), 8.18 (d, *J* = 1.8 Hz, 1H), 8.08 – 7.98 (m, 1H), 7.98 – 7.86 (m, 2H), 7.77 – 7.67 (m, 1H), 7.60 – 7.47 (m, 1H), 7.47 – 7.39 (m, 1H), 3.28 – 3.21 (m, 3H), 2.24 – 2.09 (m, 2H), 1.62 – 1.44 (m, 4H), 1.41 – 1.21 (m, 2H).

*2-(1-(2-(methylamino)-5-nitrophenyl)-1H-imidazol-4-yl)-5-(trifluoromethyl)phenyl 6-(2-(2-naphthamido)benzo[d]thiazole-6-carboxamido)hexanoate (**T1/2/RI**).*

Synthesized from **9** (90 mg, 0.195 mmol, 1.1 eq) and T1/2 (66 mg, 0.177 mmol, 1 eq) using general procedure B. The crude product was purified by flash chromatography (dichloromethane:methanol 40:1) and Isolera One flash chromatography (acetonitrile/0.1% TFA 20% → 100%) to give compound **SB91** as a pale-yellow solid. Yield (20 mg, 14%). $^1$H NMR

(400 MHz, DMSO-$d_6$) δ 13.20 (s, 1H), 8.93 – 8.80 (m, 1H), 8.58 – 8.37 (m, 2H), 8.37 – 8.25 (m, 1H), 8.22 – 8.14 (m, 2H), 8.14 – 8.09 (m, 2H), 8.07 – 7.99 (m, 3H), 7.94 (dd, *J* = 8.5, 1.8 Hz, 1H), 7.90 – 7.74 (m, 2H), 7.74 – 7.60 (m, 4H), 6.87 – 6.69 (m, 2H), 3.30 – 3.22 (m, 2H), 2.87 – 2.70 (m, 5H), 1.79 – 1.65 (m, 2H), 1.64 – 1.50 (m, 2H), 1.49 – 1.33 (m, 2H). $^{13}$C NMR (100 MHz, DMSO) δ 172.06, 166.16, 150.74, 146.99, 139.15, 135.94, 135.54, 135.33, 132.45, 131.12, 130.59, 130.09, 129.80, 129.07, 128.88, 128.80, 128.22, 127.86, 127.58, 127.58, 127.54, 127.11, 125.96, 125.68, 124.81, 124.32, 123.13, 123.09, 121.56, 121.41, 121.34, 121.32, 110.46, 34.09, 30.23, 29.30, 26.32, 24.25. HRMS *m/z* calculated for $C_{42}H_{35}O_6N_7F_3S_2$: 822.2316 $(M + H)^+$, found 822.2307.

*Dicyclopentyl ((E)-3-(4-((6-(2-(2-naphthamido)benzo[d]thiazole-6-carboxamido)hexanoyl)oxy)-3-methoxyphenyl)acryloyl)glycyl-L-valyl-D-glutamate (**N2/RI**).*
Synthesized from **9** (110 mg, 0.239 mmol, 1.1 eq) and N2 (134 mg, 0.217 mmol, 1 eq) using general procedure B. The crude product was purified Isolera One flash chromatography (acetonitrile/0.1% TFA 20% → 100%) to obtain a compound **N2/RI** as a pale-yellow solid. Yield (42 mg, 18%). $^{1}$H NMR (400 MHz, DMSO-$d_6$) δ 13.19 (s, 1H), 8.83 (s, 1H), 8.49 (t, *J* = 5.7 Hz, 1H), 8.46 – 8.41 (m, 1H), 8.37 (d, *J* = 7.4 Hz, 1H), 8.31 (t, *J* = 5.8 Hz, 1H), 8.21 (dd, *J* = 8.6, 1.8 Hz, 1H), 8.13 – 7.85 (m, 5H), 7.78 – 7.69 (m, 1H), 7.69 – 7.57 (m, 2H), 7.42 (d, *J* = 15.7 Hz, 1H), 7.33 (d, *J* = 1.8 Hz, 1H), 7.19 – 7.06 (m, 2H), 6.77 (d, *J* = 15.8 Hz, 1H), 5.23 – 4.86 (m, 2H), 4.32 – 4.12 (m, 2H), 3.90 (d, *J* = 5.7 Hz, 2H), 3.80 (s, 3H), 2.60 (t, *J* = 7.2 Hz, 2H), 2.31 (t, *J* = 7.4 Hz, 2H), 2.04 – 1.86 (m, 3H), 1.84 – 1.36 (m, 24H), 0.92 – 0.73 (m, 6H). $^{13}$C NMR (100 MHz, DMSO) δ 172.17, 171.76, 171.53, 171.46, 169.21, 166.47, 165.68, 151.54, 140.72, 138.87, 134.99, 134.30, 132.66, 132.31, 129.64, 129.53, 128.36, 128.34, 128.27, 128.12, 127.16, 125.44, 125.40, 123.69, 122.61, 121.11, 120.58, 119.37, 112.10, 77.66, 76.92, 57.83, 56.25, 51.63,

33.66, 32.62, 32.59, 32.48, 31.29, 30.43, 29.36, 26.39, 26.30, 24.79, 23.71, 23.66, 19.56, 18.34. HRMS *m/z* calculated for $C_{57}H_{67}O_{12}N_6S$: 1059.4532 $(M + H)^+$, found 1059.4524.

#### 1.2 Synthesis of linkers

*Tert-Butyl (2-(2-(2-aminoethoxy)ethoxy)ethyl)carbamate (**10**).*

A solution of Boc2O (4.365 g, 20 mmol, 1 eq) in DCM (40 mL) was added dropwise to a stirring solution of 1,2-Bis(2-aminoethoxy)ethane (14.821 g, 100 mmol, 5 eq) in DCM (100 mL) at 0 °C. The resulting mixture was stirred at rt for 20 h. Subsequently, the solution was washed with water (3 × 100 mL) and brine (100 mL), dried over anhydrous $Na_2SO_4$ and concentrated in vacuo to give compound **10** as a colourless oil (4.370 g, yield: 88%). 1H NMR (400 MHz, DMSO-$d_6$) δ 6.72 (m, 1H), 3.53 - 3.44 (m, 4H), 3.41 - 3.30 (m, 4H), 3.05 (q, 2H), 2.64 (t, 2H), 1.36 (s, 9H).

*6-ethoxy-6-oxohexan-1-aminium chloride (**11**).*

To a suspension of 6-aminohexanoic acid (4.00 g, 30.9 mmol) in EtOH (22 mL), thionyl chloride (3.3 mL, 45.7 mmol) was added. The resulting mixture was refluxed for 3 h. After concentrating the mixture in vacuo, the resulting oily residue was coevaporated three times with diethyl ether to give compound **11** as a white powder. Yield (5.96 g, 100%). $^1$H NMR (400 MHz, DMSO-$d_6$) δ 7.98 (s, 3H), 4.05 (q, J = 7.1 Hz, 2H), 2.79 – 2.60 (m, 2H), 2.29 (t, J = 7.3 Hz, 2H), 1.64 – 1.43 (m, 4H), 1.43 – 1.25 (m, 2H), 1.18 (t, J = 7.1 Hz, 3H).

*6-((tert-butoxycarbonyl)amino)hexanoic acid (**12**).*

6-aminohexanoic acid (3.00 g, 22.9 mmol) was dissolved in water (8 mL) and 1 M NaOH (40 mL), while di-tert-butyl dicarbonate (6.49 g, 29.7 mmol) was dissolved in dioxane (15 mL). The reaction mixtures were combined on ice and stirred at room temperature overnight. Subsequently 1 M NaOH (4 mL) was added to increase the pH to 10. Dioxane was evaporated off *in vacuo* and the residual was washed with ether (20 mL). Water phase was acidified with 1 M HCl to pH 2 and extracted with ethyl acetate (3 × 70 mL). Combined organic layers were dried over $Na_2SO_4$ and concentrated *in vacuo* to produce compound **12** as a colorless oil (4.55 g, 86%). $^1$H NMR (400 MHz, DMSO-$d_6$) δ 11.98 (s, 1H), 6.77 (t, *J* = 5.8 Hz, 1H), 3.57 (s, 1H), 2.88 (q, *J* = 6.9 Hz, 2H), 2.18 (t, *J* = 7.4 Hz, 2H), 1.55 – 1.41 (m, 2H), 1.41 – 1.28 (m, 11H), 1.28 – 1.13 (m, 3H).

#### 1.3 LC-MS analysis of cell incubation media and lysate samples

##### 1.3.1 Liquid Chromatography with tandem Mass spectrometry (LC-MS/MS)

For LC-MS/MS quantitation of compounds in cell incubation media and lysate samples, chromatographic separation was conducted using a Vanquish Flex system (Thermo Fisher Scientific) equipped with a Kinetex PS C18 column (100 mm × 2.1 mm, 2.6 µm particle size; Phenomenex). The column temperature was maintained at 40 °C, and the injection volume was 10 µL. The mobile phase consisted of water/MeCN/formic acid (950:5:1, v/v/v) as solvent A and water/MeCN/formic acid (50:950:1, v/v/v) as solvent B. The following gradient was applied: 0–7.0 min, 5%–75% B; 7.0–8.0 min, 75% B; 8.0–8.1 min, 75%–5% B; 8.1–10.0 min, 5% B. The flow rate was kept constant at 0.50 mL/min throughout the analysis, and the autosampler temperature was maintained at 5 °C. MS/MS analysis was performed using a TSQ Fortis triple quadrupole mass spectrometer (Thermo Fisher Scientific) operated with a heated electrospray ionization source (H-ESI) in positive ionization mode and single-reaction monitoring (SRM). For all analytes, the detector dwell time was 100 ms, and the collision-induced dissociation (CID) gas pressure was 2 mTorr, except for compound **T4**, for which it was 2.5 mTorr. All other MS/MS parameters and chromatographic conditions for each compound are summarized in Table 1.3.1. Instrument control, data acquisition, and quantification were performed using Xcalibur software (Thermo Fisher Scientific).

Table 1.3.1. The optimized instrument conditions for analyzed compounds.

| Cpd | Retention time (min) | Precursor ion (m/z) | Product ion (m/z) | Ion source voltage (V) | Collision energy (V) | Vaporizer temperature (°C) | Sheath gas (AU) | Auxiliary gas (AU) | Sweep gas (AU) |
|---|---|---|---|---|---|---|---|---|---|
| **T7/RI** | 5.75 | 409.66 | 155.05 | 3500 | 30 | 200 | 50 | 10 | 1 |
| **T7** | 3.96 | 358.15 | 184.08 | 4000 | 20 | 200 | 55 | 10 | 1 |
| **T7-linker'** | 2.73 | 244.63 | 160.08, 284.08 | 3500 | 20 | 160 | 40 | 8 | 1 |
| **RI** | 5.72 | 349.06 | 155.05 | 4000 | 20 | 180 | 50 | 10 | 1 |
| **RI-linker** | 3.61 | 479.17 | 331.05, 374.09 | 3500 | 25 | 160 | 50 | 10 | 1 |
| **T4/T7** | 5.01 | 389.15 | 334.06 | 4000 | 15 | 200 | 50 | 10 | 1 |
| **T4** | 4.58 | 352.07 | 171.05, 199.05, 260.08 | 3500 | 25 | 150 | 50 | 10 | 1 |
| **T7-linker** | 2.57 | 222.62 | 284.08, 327.12 | 3500 | 15 | 160 | 55 | 12 | 1 |
| **T4-linker** | 2.7 | 438.16 | 334.06 | 3500 | 20 | 180 | 50 | 8 | 1 |

##### 1.3.2 Determination of extra- and intracellular concentration

PBMCs, isolated from healthy donors, were seeded (0.5 x $10^6$ cells/well) in 24-well plates in 700 µL of growth medium and treated with the compounds (1 µM) or the corresponding vehicle (0.1% DMSO). After 18 h, the supernatants and cells were washed with ice-cold PBS and scraped, collected, and centrifuged (1800 rpm, 5 minutes). The supernatants were stored at -80 °C, while the cell pellets were further washed with ice-cold PBS and centrifuged (1800 rpm, 5 minutes). Then, 1 mL of the ice-cold extraction solvent MeCN/MeOH (v/v = 1:1) was added and the cell suspension was incubated at 4 °C for 30 minutes. Subsequently, cells were transferred to cold micro centrifuge tubes and supernatants were harvested (12,000 g, 15 min, 4 °C) and stored at -80 °C. Samples were analysed with LC-MS.

#### 2. Supplementary Figures

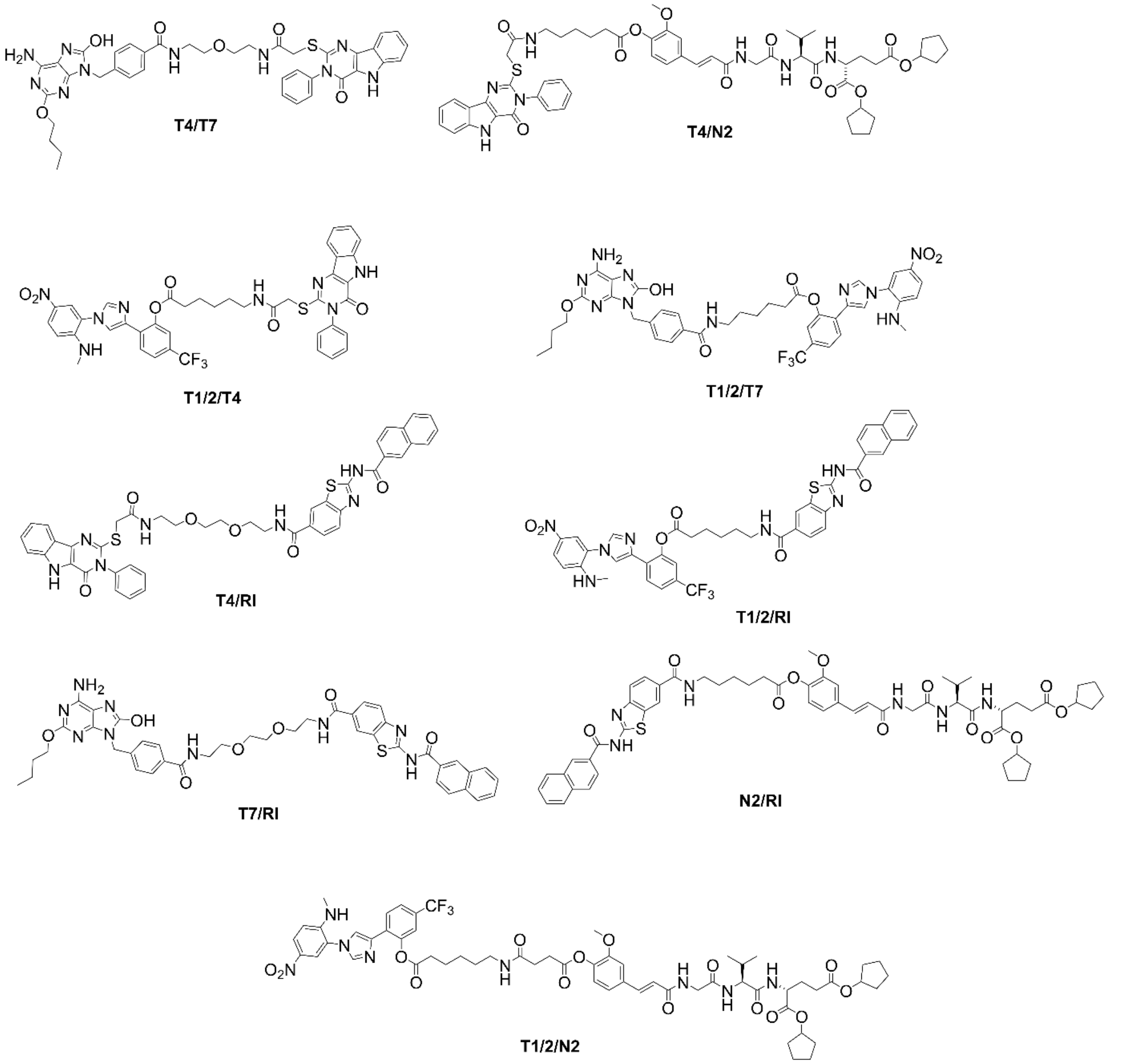

**Fig. S1**. Chemical structures of synthesized conjugated PRR ligands.

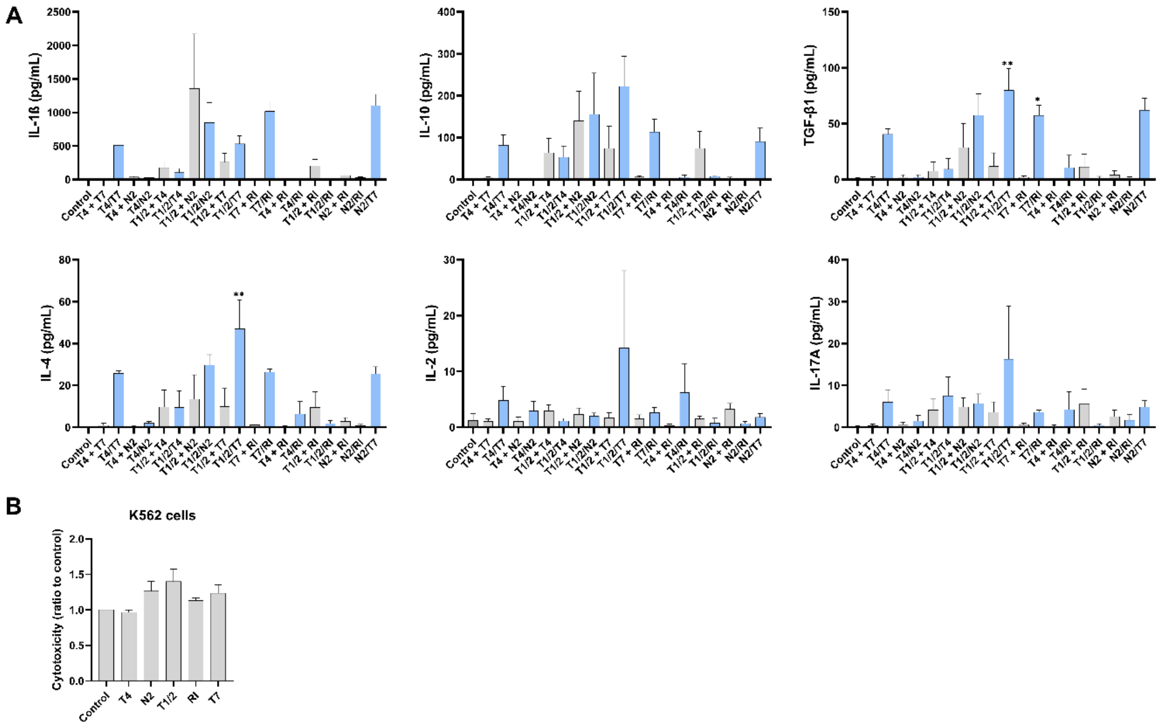

**Fig. S2**. (A) Cytokine release from human PBMCs after 18 h stimulation with conjugated agonists, unlinked mixtures (both 1 µM), **N2/T7** (1 µM, positive control), or vehicle (0.1% DMSO). Data are mean ± SEM of three independent experiments. One-way ANOVA with Bonferroni's test compared unlinked mixtures to conjugates. *, $p < 0.05$; **, $p < 0.01$ vs. vehicle. (B) PBMC cytotoxicity against K562 cells after 18 h treatment with single agonists (1 µM) or vehicle, followed by 4 h coincubation with K562 cells. Each experiment was conducted in duplicate and repeated three times. Data are relative to the negative control (NT, 0.1% DMSO) and shown as mean ± SEM of three independent experiments. **, $p < 0.01$ vs. controls (ANOVA with Bonferroni's test).

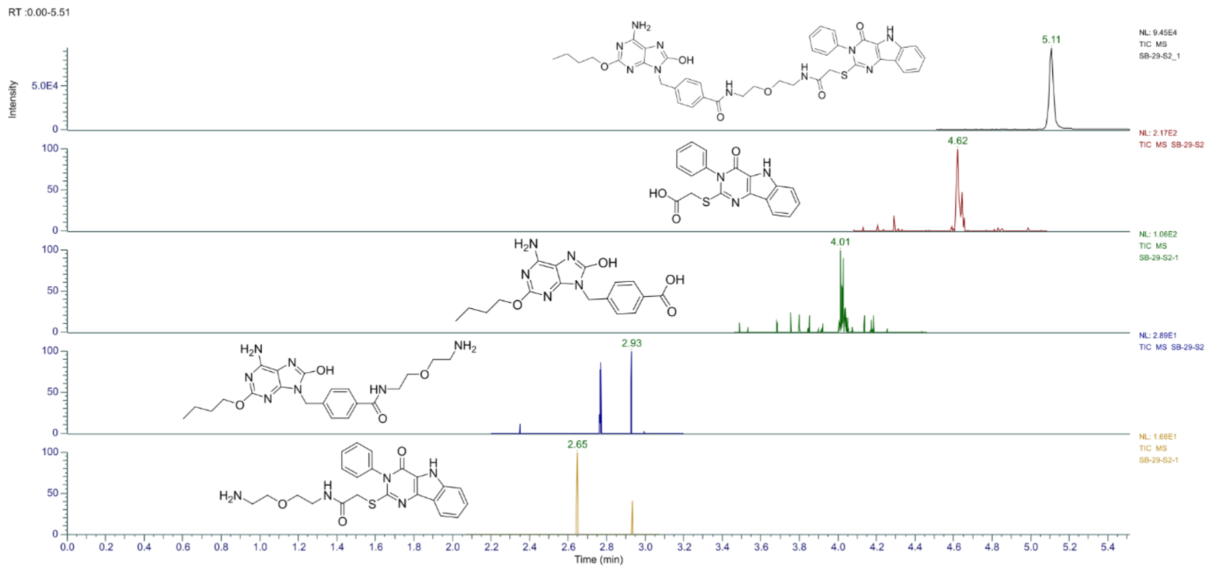

**Fig. S3.** Extracted-ion chromatograms of compound **T4/T7** and its detected metabolites in PBMC lysates following overnight stimulation with compound **T4/T7**.

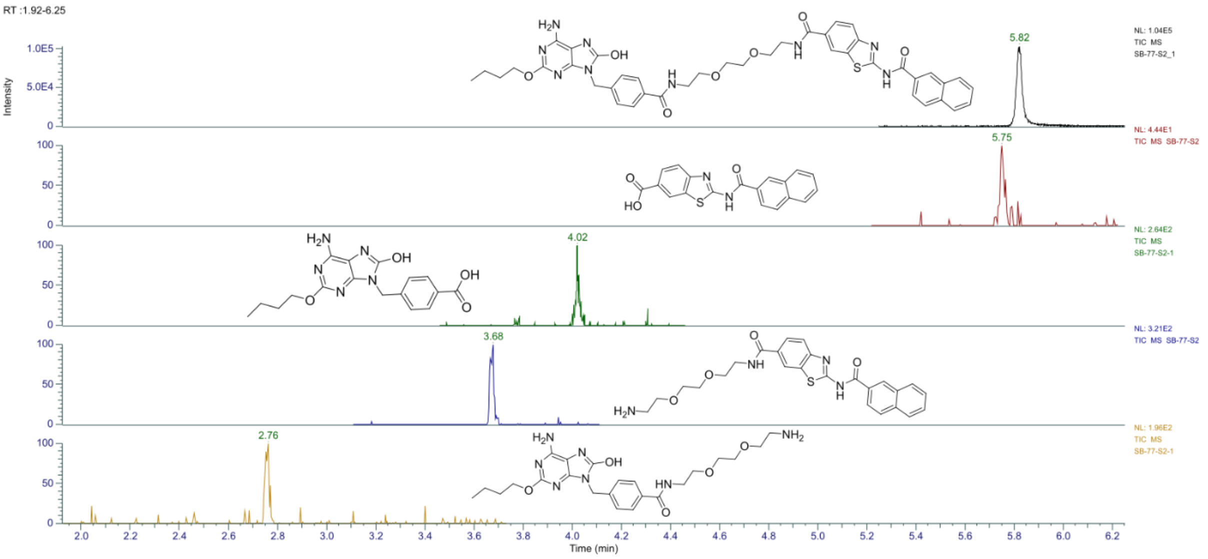

**Fig. S4.** Extracted-ion chromatograms of compound **T7/RI** and its detected metabolites in PBMC lysates following overnight stimulation with compound **T7/RI**.

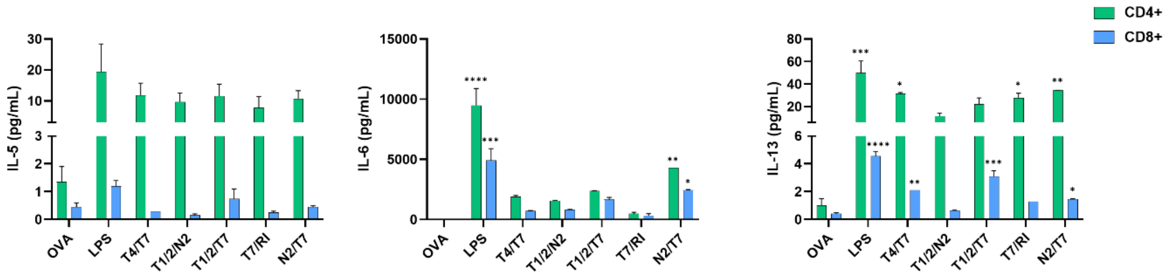

**Fig. S5**. Cytokine concentrations in BMDC-T-cell coculture supernatants following the 72 h coincubation. BMDCs from C57BL/6 mice were treated for 18 h with compounds (1 µM), LPS (1 µg/mL), or vehicle (0.1% DMSO) in the presence of OVA (50 µg/mL). CFSE-labeled OVA-specific $CD4^+$ or $CD8^+$ T cells (isolated from OT-II or OT-I mouse splenocytes, respectively) were added to the treated and washed BMDCs and cocultured for 72 h. Data are mean ± SEM of duplicates of two independent experiments. *, $p < 0.05$, **, $p < 0.01$, ***, $p < 0.001$, ****, $p < 0.0001$ versus vehicle-treated control. Statistical significance was determined using one-way ANOVA with post-hoc Dunnett's test comparing conjugates versus OVA.

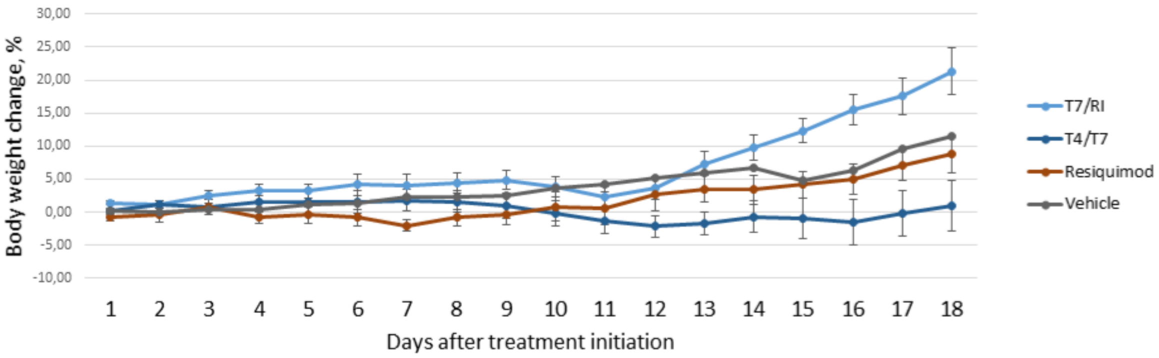

**Fig. S6**. The dynamics of body weight change values for B16F10 tumors bearing animals of groups treated with vehicle, **T4/T7**, **T7/RI**, and resiquimod. The data are presented as mean ±SEM; p-values are not depicted.

### 3. Supplementary Schemes

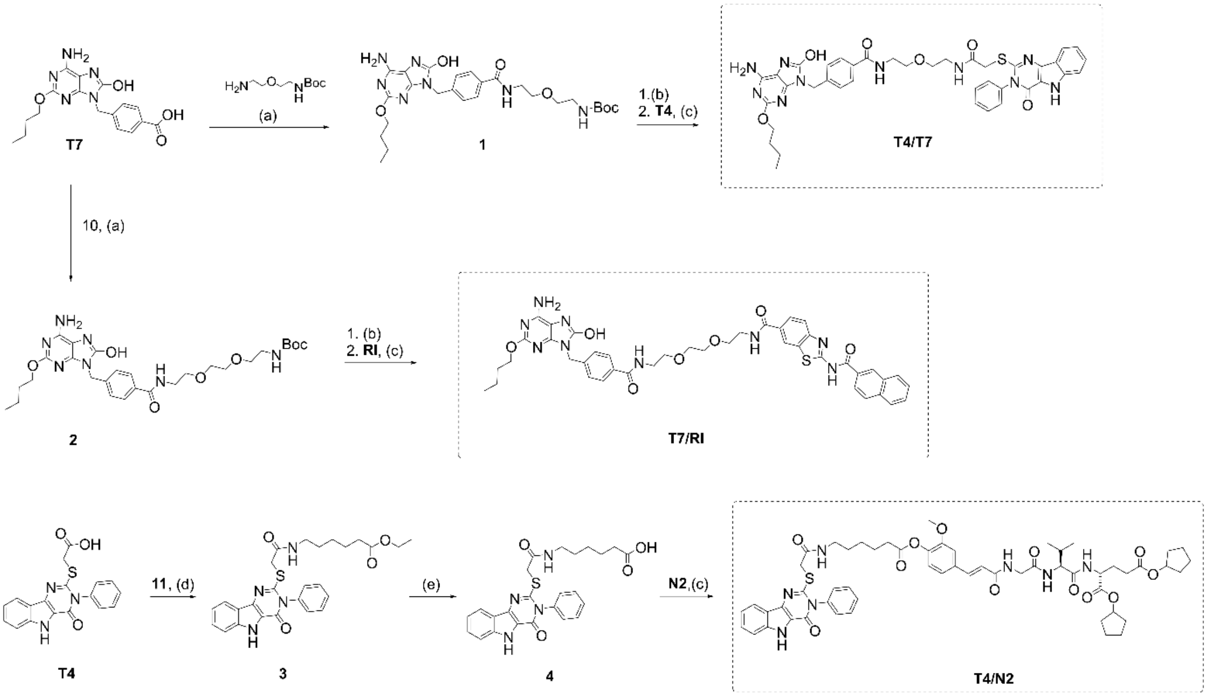

**Scheme S1**: Synthesis of conjugates **T4/T7**, **T7/RI** and **T4/N2**. Reagents and conditions: (a) DIPEA, COMU, DCM DMSO, rt.; (b) TFA, DCM (1:5), rt.; (c) DIPEA, COMU, DMF, rt.; (d) DIPEA, HATU, DMF, rt.; (e) NaOH, MeOH, rt.

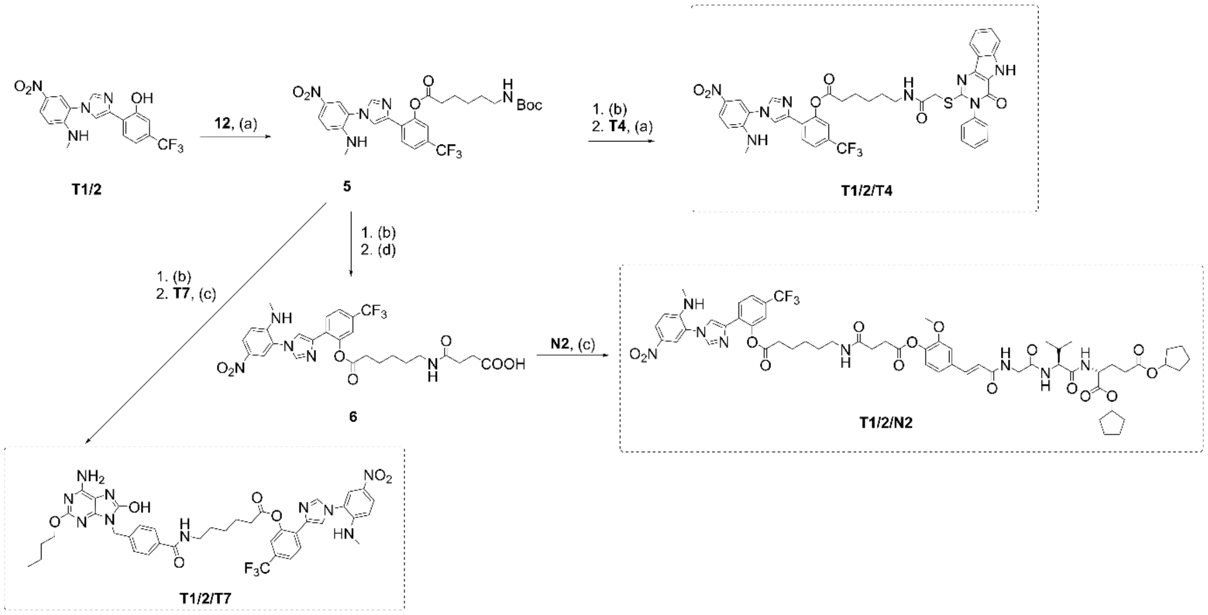

**Scheme S2**: Synthesis of conjugates **T1/2/T4**, **T1/2/T7** and **T1/2/N2**. Reagents and conditions: (a) DIPEA, HATU, DMF, rt.; (b) TFA, DCM (1:5), rt.; (c) DIPEA, COMU, DMF, rt.; (d) succinic anhydride, $Et_3N$, DMAP, DMF, rt.

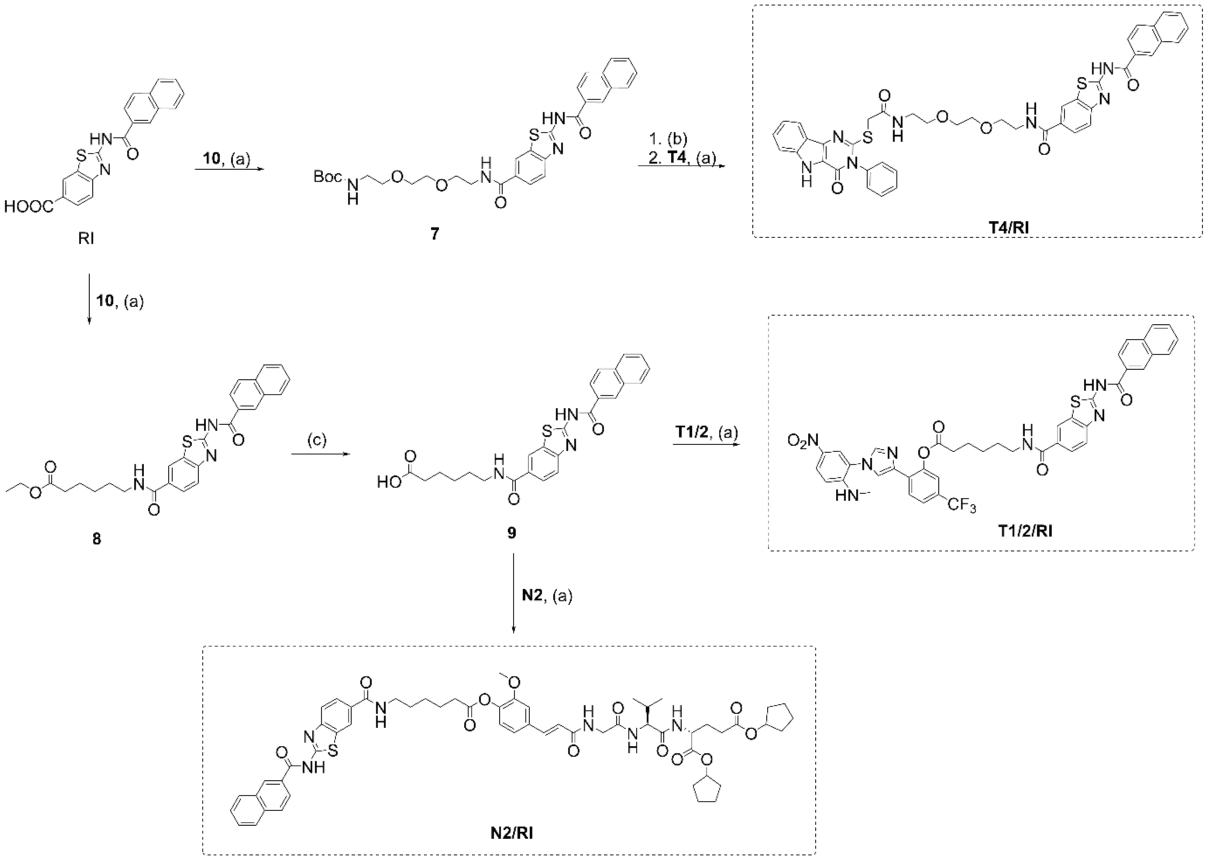

**Scheme S3**: Synthesis of conjugates **T4/RI**, **T1/2/RI** and **N2/RI**. Reagents and conditions: (a) DIPEA, COMU, DMF, rt.; (b) TFA, DCM (1:5), rt.; (c) NaOH, MeOH, rt.

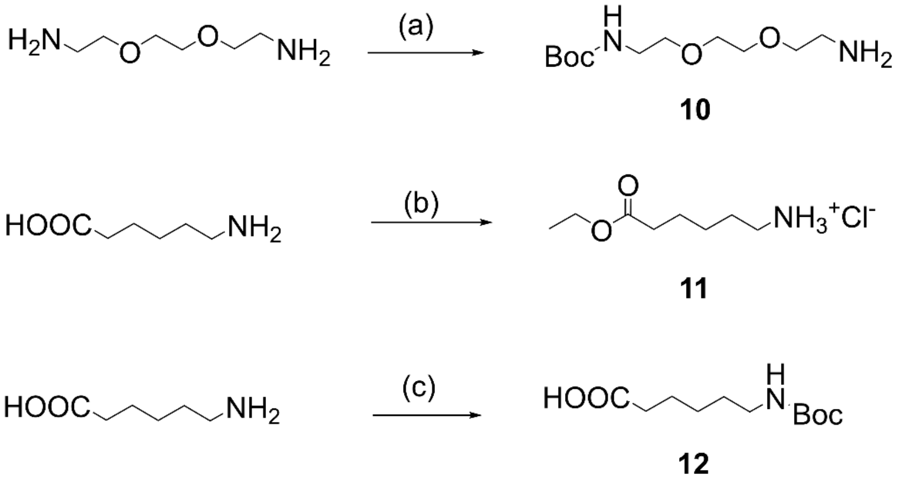

**Scheme S4**: Synthesis of linkers **10**, **11** and **12**. Reagents and conditions: (a) $Boc_2O$, DCM, rt.; (b) $SOCl_2$, EtOH, reflux, (c) $Boc_2O$, NaOH/$H_2O$, dioxane, rt.

### 4. Supplementary Tables

**Table S1.** Cytokine concentrations after treating human PBMCs with conjugates and unlinked mixtures of agonists.

| | Average ± SEM [pg/mL] | | | | | | | | | | | | |
|---|---|---|---|---|---|---|---|---|---|---|---|---|---|
| | **IL-4** | **IL-2** | **IP-10** | **IL-1β** | **TNF-α** | **MCP-1** | **IL-17A** | **IL-6** | **IL-10** | **IFN-γ** | **IL-12p70** | **IL-8** | **TGF-β1** |
| **Control** | 0.3 ± 0.2 | 1.2 ± 0.8 | 21 ± 3.9 | 0 ± 0 | 0.8 ± 0.3 | 35.1 ± 20.8 | 0.2 ± 0.1 | 15.2 ± 3.2 | 0.7 ± 0.3 | 0.3 ± 0.3 | 0.3 ± 0.2 | 45.1 ± 5.8 | 0.7 ± 0.4 |
| **T1/2** | 26 ± 9.4 | 4.5 ± 3.7 | 106.4 ± 2 | 188.9 ± 80.8 | 1497.8 ± 1058.4 | 9000 ± 0 | 9 ± 4.7 | 6858.2 ± 2141.8 | 81.9 ± 51.5 | 52.9 ± 27.5 | 15.6 ± 7.6 | 10000 ± 0 | 32 ± 15.2 |
| **T4** | 0.7 ± 0.2 | 0.6 ± 0.4 | 43.4 ± 10.6 | 4.3 ± 2.2 | 2.3 ± 1 | 5.9 ± 1.7 | 0.6 ± 0.3 | 15.6 ± 6.3 | 3.5 ± 1.6 | 1.7 ± 0.9 | 1 ± 0.4 | 111.3 ± 37.9 | 3.6 ± 1.9 |
| **T7** | 0.3 ± 0.2 | 0.9 ± 0.4 | 79.2 ± 16.2 | 9 ± 5.6 | 25.6 ± 18.1 | 84.4 ± 51.2 | 1.3 ± 0.5 | 276.6 ± 97.1 | 4.6 ± 2.4 | 3.2 ± 1 | 1.5 ± 0.4 | 528.9 ± 272.7 | 3.8 ± 1 |
| **N2** | 1.7 ± 0.8 | 1 ± 0.6 | 104.1 ± 2.2 | 51.9 ± 8 | 94.6 ± 23.4 | 338.6 ± 104.6 | 2.5 ± 0.5 | 940.8 ± 70.2 | 1.8 ± 0.2 | 4.2 ± 0.3 | 1.5 ± 0 | 8764.3 ± 3587.7 | 7.7 ± 1.4 |
| **RI** | 0.2 ± 0.2 | 3.2 ± 1.3 | 28.3 ± 15.4 | 1.9 ± 1.9 | 3.4 ± 2.4 | 57.5 ± 24.7 | 0.1 ± 0.1 | 28.4 ± 5.6 | 0.7 ± 0.4 | 0.4 ± 0.4 | 0.1 ± 0.1 | 129.5 ± 44.1 | 0.2 ± 0.2 |
| **T1/2+T4** | 9.8 ± 8.2 | 3 ± 0.9 | 29.3 ± 24.7 | 176.2 ± 94.2 | 405.2 ± 239.3 | 7109.8 ± 1257.2 | 4.1 ± 2.6 | 6169.2 ± 2830.8 | 63.6 ± 33.9 | 55.5 ± 40.7 | 7.2 ± 4.9 | 6193.1 ± 2703.5 | 7.7 ± 7.7 |
| **T1/2+T7** | 9.9 ± 8.6 | 1.7 ± 0.9 | 32.1 ± 28.2 | 259.8 ± 131.5 | 651.2 ± 364.9 | 5705.8 ± 1841.8 | 3.6 ± 2.5 | 6203.6 ± 2796.4 | 74 ± 52.7 | 31.3 ± 28.8 | 6.9 ± 5.7 | 5697.6 ± 2499.5 | 11.9 ± 11.9 |
| **T1/2+N2** | 13.6 ± 11.5 | 2.4 ± 1.1 | 37 ± 31.3 | 1358.7 ± 814.2 | 1431.1 ± 945.6 | 7538.3 ± 1461.7 | 4.9 ± 2.2 | 8384.8 ± 615.2 | 140.2 ± 70.7 | 56.3 ± 29.8 | 10.4 ± 6.9 | 7495.1 ± 3018.7 | 28.6 ± 21.4 |
| **T1/2+RI** | 9.5 ± 7.4 | 1.6 ± 0.4 | 35.1 ± 30.5 | 207.5 ± 93.1 | 524.6 ± 352.6 | 7712.8 ± 1152.7 | 5.5 ± 3.5 | 6633.8 ± 2366.2 | 74.4 ± 40.3 | 42.1 ± 25.1 | 7.1 ± 4.5 | 6602.9 ± 2653.7 | 11.3 ± 11.3 |
| **T4+T7** | 1.1 ± 0.8 | 1.1 ± 0.5 | 25.3 ± 22 | 11.2 ± 2.7 | 15 ± 3.9 | 311.3 ± 168.4 | 0.4 ± 0.4 | 205.8 ± 20 | 5.1 ± 1.3 | 1.6 ± 1.1 | 0.3 ± 0.3 | 481.5 ± 97.5 | 1.2 ± 1.2 |
| **T4+N2** | 0.5 ± 0.3 | 1.1 ± 0.7 | 31.2 ± 30.8 | 39.3 ± 6.6 | 44.9 ± 21.8 | 747.9 ± 267.9 | 0.6 ± 0.6 | 576 ± 183.9 | 1.7 ± 0.3 | 2 ± 2 | 0.5 ± 0.5 | 3687 ± 1076.3 | 2.2 ± 2.2 |
| **T4+RI** | 0.4 ± 0.2 | 0.3 ± 0.3 | 18.4 ± 17 | 1 ± 1 | 2.1 ± 1.7 | 17.8 ± 14.6 | 0.3 ± 0.3 | 27.2 ± 17.2 | 0.7 ± 0.7 | 0.6 ± 0.6 | 0.2 ± 0.2 | 109.2 ± 62.8 | 0.3 ± 0.3 |
| **T7+RI** | 1.1 ± 0.2 | 1.6 ± 0.6 | 45 ± 17.7 | 14.6 ± 6.1 | 15.9 ± 5.2 | 814.5 ± 755.5 | 0.6 ± 0.3 | 328.3 ± 129.9 | 6.4 ± 2.7 | 2.7 ± 1.2 | 1 ± 0.5 | 498.5 ± 234.6 | 2.2 ± 1.1 |
| **N2+RI** | 2.8 ± 1.8 | 3.2 ± 1 | 36.9 ± 28.5 | 64.9 ± 19.4 | 72.2 ± 35.2 | 1888.7 ± 1065.2 | 2.6 ± 1.6 | 1125.7 ± 366.7 | 3.9 ± 1.5 | 7.5 ± 1.5 | 2.1 ± 1.5 | 8789 ± 2211 | 4 ± 4 |
| **T4/T7** | 25.8 ± 1.1 | 4.9 ± 2.5 | 1595.8 ± 218 | 513.2 ± 39.4 | 1320.3 ± 367.1 | 9000 ± 0 | 6 ± 2.9 | 8933.6 ± 36.4 | 81.9 ± 24.8 | 1061.7 ± 120.6 | 82.4 ± 23.4 | 4476.2 ± 2013 | 40.6 ± 4.6 |
| **T4/N2** | 1.9 ± 0.9 | 3 ± 1.6 | 34 ± 31.5 | 27.6 ± 6.5 | 38.8 ± 18.2 | 1435.6 ± 880.1 | 1.5 ± 1.3 | 484.7 ± 175.9 | 2.3 ± 0.5 | 3.1 ± 2.3 | 0.8 ± 0.8 | 5473.7 ± 2849.9 | 2.2 ± 2.2 |
| **T1/2/T4** | 9.4 ± 8 | 1.2 ± 0.4 | 29.6 ± 27.1 | 105.8 ± 56.2 | 190.5 ± 141.2 | 6117.8 ± 2570.4 | 7.5 ± 4.4 | 6059.6 ± 2940.4 | 53.5 ± 26.5 | 12.2 ± 7.4 | 6.1 ± 4.2 | 5039.3 ± 1861.2 | 9.4 ± 9.4 |
| **T1/2/N2** | 29.7 ± 5 | 2 ± 0.6 | 106.8 ± 9.2 | 848.9 ± 298.5 | 4768.4 ± 2311.5 | 7885.4 ± 608.1 | 5.7 ± 2.3 | 9000 ± 0 | 156.1 ± 97.5 | 198.1 ± 67.4 | 28.8 ± 4.2 | 8939.6 ± 762.1 | 57.6 ± 19.2 |

| | | | | | | | | | | | | | |
|---|---|---|---|---|---|---|---|---|---|---|---|---|---|
| **T1/2/T7** | 47 ± 13.9 | 14.2 ± 13.8 | 379.8 ± 142.4 | 533.9 ± 118.8 | 3045 ± 2094.4 | 8212.1 ± 366.8 | 16.3 ± 12.7 | 9000 ± 0 | 222.3 ± 71.6 | 579.4 ± 398.1 | 58.9 ± 13.8 | 8048.5 ± 1951.5 | 79.9 ± 19.2 |
| **T7/RI** | 26.5 ± 1.4 | 2.7 ± 0.9 | 1974.8 ± 284.1 | 1019.1 ± 147 | 2037.7 ± 483 | 8910.8 ± 89.2 | 3.6 ± 0.4 | 9000 ± 0 | 113.4 ± 31.1 | 3335.1 ± 582.8 | 95.5 ± 18 | 5132.5 ± 2443.8 | 57.3 ± 9.2 |
| **T4/RI** | 6.3 ± 6 | 6.3 ± 5 | 35.4 ± 33.8 | 9.1 ± 6.1 | 6.9 ± 6.4 | 32 ± 17.4 | 4.3 ± 4.3 | 26.8 ± 12.5 | 5.6 ± 4.9 | 5.4 ± 5.4 | 4.5 ± 4.5 | 91.7 ± 27 | 10.8 ± 10.8 |
| **T1/2/RI** | 1.8 ± 1.3 | 0.9 ± 0.8 | 34.6 ± 33.7 | 4.5 ± 3.6 | 6.5 ± 2.2 | 2062.7 ± 1461.7 | 0.4 ± 0.4 | 279.1 ± 84.6 | 7.8 ± 1.8 | 1.3 ± 1.2 | 0.6 ± 0.6 | 1230.7 ± 504.1 | 1.4 ± 1.4 |
| **N2/RI** | 0.7 ± 0.6 | 0.7 ± 0.4 | 32.4 ± 30.7 | 29 ± 11.2 | 42.2 ± 24.2 | 279.8 ± 164.7 | 1.7 ± 1.5 | 434.6 ± 257.1 | 1.3 ± 0.7 | 1.4 ± 1.4 | 0.6 ± 0.6 | 1438.9 ± 549.6 | 1.3 ± 1.3 |
| **N2/T7** | 25.7 ± 3.2 | 1.8 ± 0.6 | 1765.2 ± 259.5 | 1106.8 ± 166.3 | 2391.5 ± 594.1 | 8814.4 ± 110.7 | 4.8 ± 1.6 | 9000 ± 0 | 90.3 ± 32.1 | 2149.4 ± 515.1 | 146.6 ± 55.5 | 8568.5 ± 1431.5 | 62 ± 10.9 |

**Table S2A.** Contributions of individual protein analytes to the two principal component axes.

| **Loadings** | | |
|---|---|---|
| **Variable** | **PC1** | **PC2** |
| **IL-4** | 0.967 | -0.125 |
| **IL-2** | 0.631 | -0.294 |
| **IP-10** | 0.617 | 0.753 |
| **IL-1β** | 0.794 | 0.130 |
| **TNF-α** | 0.858 | -0.151 |
| **MCP-1** | 0.894 | -0.088 |
| **IL-17A** | 0.811 | -0.442 |
| **IL-6** | 0.930 | -0.076 |
| **IL-10** | 0.941 | -0.233 |
| **IFN-γ** | 0.613 | 0.744 |
| **IL-12p70** | 0.841 | 0.494 |
| **IL-8** | 0.648 | -0.418 |
| **TGF-β1** | 0.966 | -0.006 |

**Table S2B.** Contributions of samples to the two principal component axes.

| **Samples** | **PC1** | **PC2** |
|---|---|---|
| **NT** | -2.494 | 0.485 |
| **T1/2** | 2.607 | -1.774 |
| **T4** | -2.418 | 0.514 |
| **T7** | -2.262 | 0.437 |
| **N2** | -1.433 | -0.364 |
| **RI** | -2.352 | 0.345 |
| **T1/2 + T4** | 0.405 | -0.805 |
| **T1/2 + T7** | 0.361 | -0.652 |
| **T1/2 + N2** | 2.684 | -0.963 |
| **T1/2 + RI** | 0.670 | -0.910 |
| **T4 + T7** | -2.356 | 0.419 |
| **T4 + N2** | -2.059 | 0.142 |
| **T4 + RI** | -2.554 | 0.532 |
| **T7 + RI** | -2.211 | 0.383 |
| **N2 + RI** | -1.151 | -0.635 |
| **T4/T7** | 4.202 | 2.275 |
| **T4/N2** | -1.657 | -0.244 |
| **T1/2/T4** | 0.193 | -0.861 |
| **T1/2/N2** | 4.534 | -1.310 |
| **T1/2/T7** | 7.212 | -2.276 |
| **T7/RI** | 5.849 | 4.864 |
| **T4/RI** | -1.417 | -0.223 |
| **T1/2/RI** | -2.139 | 0.340 |
| **N2/RI** | -2.216 | 0.280 |

**Table S3.** Percentage of dead K562 cells, dead PBMCs and ratio of dead K562 cells compared to untreated control.

| | **Average ± SEM** | | |
|---|---|---|---|
| | **% dead K562** | **dead K562, ratio to NT** | **% dead PBMCs** |
| **Control** | 24.2 ± 2.1 | 1.0 ± 0.0 | 2.23 ± 0.32 |
| **T4** | 22.0 ± 2.8 | 0.94 ± 0.03 | 1.79 ± 0.74 |
| **N2** | 25.8 ± 1.8 | 1.25 ± 0.13 | 1.05 ± 0.23 |
| **T1/2** | 29.3 ± 5.3 | 1.38 ± 0.15 | 1.36 ± 0.56 |
| **RI** | 30.4 ± 3.3 | 1.11 ± 0.03 | 1.27 ± 0.30 |
| **T7** | 25.9 ± 4.2 | 1.23 ± 0.11 | 1.76 ± 0.21 |
| **T4 + T7** | 30.6 ± 1.0 | 1.14 ± 0.12 | 1.35 ± 0.19 |
| **T4/T7** | 77.3 ± 2.5 | 3.74 ± 0.23 | 1.18 ± 0.31 |
| **T4 + N2** | 30.9 ± 3.4 | 1.0 ± 0.0 | 1.28 ± 0.33 |
| **T4/N2** | 28.3 ± 2.9 | 1.01 ± 0.16 | 1.27 ± 0.17 |
| **T1/2 + T4** | 43.4 ± 4.9 | 1.40 ± 0.31 | 8.71 ± 4.10 |
| **T1/2/T4** | 26.3 ± 3.5 | 1.26 ± 0.13 | 2.02 ± 0.35 |
| **T1/2 + N2** | 33.4 ± 4.3 | 1.60 ± 0.14 | 2.92 ± 1.86 |
| **T1/2/N2** | 35.7 ± 3.2 | 2.11 ± 0.18 | 2.85 ± 1.58 |
| **T1/2 + T7** | 36.6 ± 5.7 | 1.73 ± 0.14 | 1.42 ± 0.53 |
| **T1/2/T7** | 64.8 ± 7.0 | 3.12 ± 0.30 | 1.36 ± 0.41 |
| **T7 + RI** | 31.2 ± 1.2 | 1.19 ± 0.10 | 1.04 ± 0.13 |
| **T7/RI** | 79.1 ± 1.7 | 3.84 ± 0.30 | 1.45 ± 0.34 |
| **T4/RI** | 36.8 ± 2.5 | 1.17 ± 0.05 | 1.11 ± 0.02 |
| **T4/RI** | 25.2 ± 2.0 | 0.88 ± 0.08 | 2.27 ± 0.76 |
| **T1/2 + RI** | 45.3 ± 0.1 | 1.45 ± 0.17 | 9.30 ± 4.30 |
| **T1/2/RI** | 38.0 ± 0.8 | 1.35 ± 0.17 | 1.55 ± 0.20 |
| **N2 + RI** | 37.4 ± 0.2 | 1.19 ± 0.13 | 1.46 ± 0.31 |
| **N2/RI** | 32.4 ± 2.2 | 1.13 ± 0.06 | 1.20 ± 0.12 |
| **N2/T7** | 77.6 ± 3.5 | 3.75 ± 0.25 | 1.29 ± 0.28 |
| **IL-2** | 75.5 ± 3.9 | 3.18 ± 0.40 | 1.67 ± 0.31 |

**Table S4A.** Extracellular and intracellular concentrations of **T4/T7**, **T7/RI** and single agonists after treatment of PBMCs (all at 1 μM).

| **Compounds** | **Extracellular concentration (ng/mL)** | **Intracellular concentration (ng/mL)** |
|---|---|---|
| **T4** | 252.77 | 0.95 |
| **T7** | 331.18 | 0.90 |
| **RI** | 302.32 | 1.86 |
| **T4/T7** | 667.22 | 52.44 |
| **T7/RI** | 209.44 | 126.21 |

**Table S4B.** Intracellular concentrations of **T4/T7** and detected degradation products/metabolites after treatment of PBMCs with 1 μM **T4/T7**.

| **Compounds** | **Structures** | **Intracellular concentration (ng/mL)** |
|---|---|---|
| **T4/T7** | 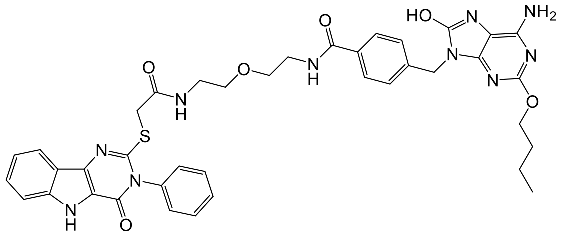 | 52.44 |
| **T4** | 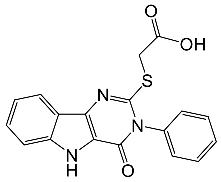 | 0.59 |
| **T7** | 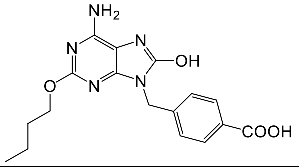 | 0.34 |
| **T4-linker** | 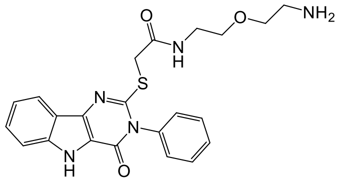 | 0.80 |
| **T7-linker** | 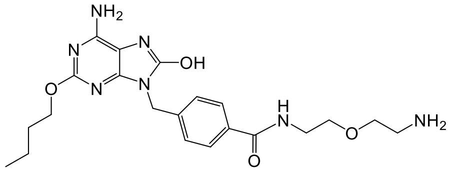 | 0.33 |

**Table S4C.** Intracellular concentrations of **T7/RI** and detected degradation products/metabolites after treatment of PBMCs with 1 µM **T7/RI**.

| **Compounds** | **Structures** | **Intracellular concentration (ng/mL)** |
|---|---|---|
| **T7/RI** | 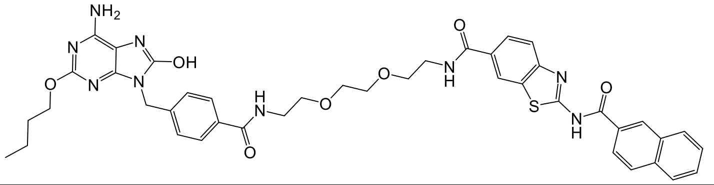 | 126.21 |
| **T7** | 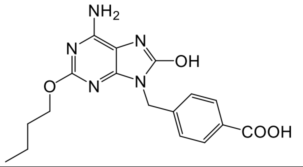 | 0.36 |
| **RI** | 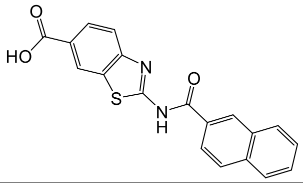 | 0.14 |
| **T7-linker'** | 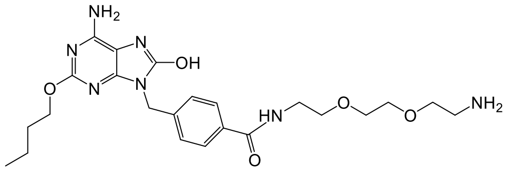 | 0.61 |
| **RI-linker** | 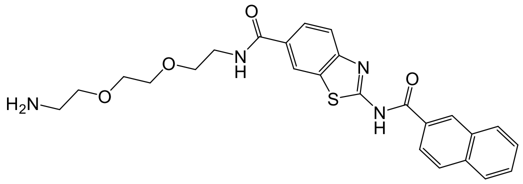 | nd |

**Table S5A**: Cytokine concentrations after treatment of human PBMCs with conjugates **T4/T7** and **T7/RI** and the corresponding antagonists.

| | Average ± SEM [pg/mL] | | | | | | | |
|---|---|---|---|---|---|---|---|---|
| | **IP-10** | **IL-1b** | **TNF-α** | **MCP-1** | **IL-6** | **IL-10** | **IFN-γ** | **IL-8** |
| **NT** | 50.23 ± 24.06 | 1.35 ± 0 | 0.22 ± 0.11 | 547.01 ± 104.2 | 71.02 ± 8.21 | 0.65 ± 0.18 | 0.23 ± 0.06 | 347.69 ± 51.88 |
| **T4/T7** | 700.01 ± 149.69 | 1.35 ± 0 | 311.34 ± 303.24 | 2867.75 ± 287.03 | 133.49 ± 39.62 | 0.84 ± 0.2 | 3.45 ± 1.37 | 420.57 ± 66.55 |
| **T4/T7+TAK242** | 11.88 ± 4.97 | 1.35 ± 0 | 0.14 ± 0.08 | 112.51 ± 37.81 | 12.69 ± 7.52 | 0.08 ± 0.01 | 0.06 ± 0.02 | 20.63 ± 2.99 |
| **T4/T7+M5049** | 34.43 ± 20.19 | 2.39 ± 1.04 | 0.23 ± 0.07 | 337.07 ± 10.33 | 49.8 ± 15.07 | 0.69 ± 0.16 | 0.26 ± 0.16 | 296.4 ± 38.6 |
| **T4/T7+TAK242+M5049** | 1.06 ± 0.35 | 1.35 ± 0 | 0.03 ± 0.01 | 8.8 ± 1.99 | 1.53 ± 0.8 | 0.07 ± 0 | 0.04 ± 0.01 | 5.63 ± 1.52 |
| **T7/RI** | 938.2 ± 166.38 | 150.49 ± 77 | 171.72 ± 52.23 | 6829.18 ± 897.04 | 2440.01 ± 854.67 | 66.8 ± 22.58 | 95.49 ± 51.03 | 1043.24 ± 169.12 |
| **T7/RI+M5049** | 31.65 ± 5.79 | 1.79 ± 0.44 | 0.47 ± 0.11 | 489.25 ± 43.03 | 74.02 ± 19.06 | 1.33 ± 0.87 | 0.28 ± 0.08 | 248.59 ± 79.24 |
| **T7/RI+MRT67307** | 5.56 ± 0.61 | 1.35 ± 0 | 0.04 ± 0.01 | 105.38 ± 8.71 | 39.66 ± 6.46 | 0.18 ± 0.05 | 0.08 ± 0.02 | 168.77 ± 37.88 |
| **T7/RI+M5049+MRT67307** | 7.04 ± 1.67 | 1.35 ± 0 | 0.04 ± 0.01 | 66.4 ± 9.53 | 66.94 ± 14.34 | 0.25 ± 0.1 | 0.15 ± 0.06 | 313.17 ± 78.07 |

**Table S5B**: Percentage of dead K562 cells, dead PBMCs and ratio of dead K562 cells compared to untreated control after pretreatment of human PBMCs with conjugates **T4/T7** and **T7/RI** and the corresponding antagonists.

| | Average ±SEM | | |
|---|---|---|---|
| | **% dead K562** | **dead K562, ratio to NT** | **% dead PBMCs** |
| **NT** | 56.4 ± 7.5 | 1.0 ± 0.0 | 1.6 ± 0.4 |
| **IL-2** | 88.9 ± 0.8 | 1.6 ± 0.1 | 1.6 ± 0.1 |
| **T4/T7** | 84.2 ± 1.3 | 1.5 ± 0.1 | 1.4 ± 0.2 |
| **T4/T7+TAK242** | 37.8 ± 4.9 | 0.7 ± 0.1 | 3.2 ± 0.5 |
| **T4/T7+M5049** | 47.4 ± 1.3 | 0.8 ± 0.0 | 5.1 ± 0.5 |
| **T4/T7+TAK242+M5049** | 16.7 ± 1.2 | 0.3 ± 0.0 | 6.5 ± 0.5 |
| **T7/RI** | 80.7 ± 1.8 | 1.5 ± 0.1 | 1.3 ± 0.1 |
| **T7/RI+M5049** | 7.7 ± 0.3 | 0.1 ± 0.0 | 2.9 ± 0.7 |
| **T7/RI+MRT67307** | 45.4 ± 2.9 | 0.8 ± 0.0 | 6.0 ± 0.6 |
| **T7/RI+M5049+MRT67307** | 5.0 ± 2.1 | 0.1 ± 0.0 | 6.1 ± 0.3 |

**Table S6A**: Proliferation indexes of $CD4^+$ and $CD8^+$ T cells.

| | Average ± SEM | |
|---|---|---|
| | **$CD4^+$ (OT-II) proliferation index** | **$CD8^+$ (OT-I) proliferation index** |
| **w/o OVA** | 1.1 ± 0.0 | 1.1 ± 0.0 |
| **OVA** | 1.4 ± 0.1 | 2.0 ± 0.1 |
| **LPS** | 1.4 ± 0.0 | 2.4 ± 0.1 |
| **T4/T7** | 2.1 ± 0.1 | 3.2 ± 0.1 |
| **T1/2/N2** | 2.3 ± 0.0 | 3.6 ± 0.0 |
| **T1/2/T7** | 2.2 ± 0.0 | 3.5 ± 0.1 |
| **T7/RI** | 2.1 ± 0.0 | 3.5 ± 0.1 |
| **N2/T7** | 2.0 ± 0.1 | 3.2 ± 0.1 |

**Table S6B**: Cytokine concentrations in BMDC-T-cell coculture.

| | | **Average ± SEM** | | | | | | | | | | | |
|---|---|---|---|---|---|---|---|---|---|---|---|---|---|
| | | **IL-2** | **IL-4** | **IL-5** | **IL-6** | **IL-9** | **IL-10** | **IL-13** | **IL-17A** | **IL-17F** | **IL-22** | **IFN-γ** | **TNF-α** |
| **CD4+** | **OVA only** | 41.8 ± 3.9 | 0.7 ± 0.4 | 1.3 ± 0.5 | 11 ± 1.1 | 2.7 ± 1.5 | 0 ± 0 | 1 ± 0.5 | 6 ± 3.5 | 0.9 ± 0 | 2.2 ± 0.4 | 33.9 ± 1.7 | 13.1 ± 3.5 |
| | **LPS** | 873.5 ± 102.7 | 3.7 ± 0.2 | 19.3 ± 9.1 | 9490.7 ± 1406.1 | 520.2 ± 47.2 | 25.7 ± 4.4 | 49.7 ± 10.7 | 1655.4 ± 217.3 | 693.4 ± 68.5 | 5923.9 ± 784 | 6377 ± 1114.1 | 218.3 ± 26.5 |
| | **T4/T7** | 790.8 ± 11.1 | 5.8 ± 1.1 | 11.8 ± 3.9 | 1910.9 ± 97.6 | 349.9 ± 34.2 | 5.8 ± 0 | 31.5 ± 0.9 | 1213.6 ± 221.3 | 436.9 ± 62.2 | 3485.4 ± 174.5 | 2265.7 ± 151.3 | 176.5 ± 6.2 |
| | **T1/2/N2** | 1108 ± 83.6 | 2.4 ± 0.3 | 9.7 ± 2.8 | 1560 ± 39.9 | 368.5 ± 32.5 | 3.3 ± 0.3 | 11.8 ± 2.5 | 431.7 ± 4.7 | 199.2 ± 6.4 | 2446.9 ± 47 | 1095.1 ± 44 | 137 ± 7.8 |
| | **T1/2/T7** | 1052 ± 127.5 | 3.8 ± 1.3 | 11.5 ± 3.9 | 2399.8 ± 4 | 354.9 ± 11.8 | 10 ± 0.3 | 22 ± 5.8 | 1010.3 ± 53.9 | 415.2 ± 11 | 3557.6 ± 421.9 | 2268.4 ± 55.6 | 145.2 ± 12.2 |
| | **T7/RI** | 476.7 ± 89.1 | 39.1 ± 4.8 | 7.7 ± 3.7 | 491.9 ± 117.6 | 334.8 ± 9.2 | 2.3 ± 2.3 | 27.9 ± 4.1 | 925.1 ± 177.3 | 439.5 ± 105.5 | 1874.4 ± 229.3 | 1041.4 ± 132.8 | 140.7 ± 24.1 |
| | **N2/T7** | 722.9 ± 84 | 5 ± 1.4 | 10.7 ± 2.6 | 4304.1 ± 0.6 | 328.1 ± 4.5 | 14.4 ± 0.2 | 34.4 ± 0.2 | 1336.2 ± 2.4 | 623.2 ± 63.1 | 4058.6 ± 428.9 | 2673.2 ± 559.8 | 165.6 ± 9.9 |
| **CD8+** | **OVA only** | 123.2 ± 30.7 | 0.2 ± 0.2 | 0.5 ± 0.2 | 2.2 ± 0.2 | 0.8 ± 0.2 | 0 ± 0 | 0.4 ± 0.1 | 0.3 ± 0 | 0.7 ± 0 | 0.5 ± 0.2 | 15.9 ± 8 | 5.7 ± 0.8 |
| | **LPS** | 36.6 ± 6.6 | 0.8 ± 0.1 | 1.2 ± 0.2 | 4925.2 ± 965.8 | 33.8 ± 4.9 | 8.8 ± 0.5 | 4.5 ± 0.4 | 403.4 ± 43.8 | 157.9 ± 23.3 | 826.5 ± 19.1 | 3326.4 ± 263.9 | 172.1 ± 11.8 |
| | **T4/T7** | 117.8 ± 29.4 | 0 ± 0 | 0.3 ± 0 | 732.8 ± 22.5 | 5.5 ± 0.1 | 0 ± 0 | 2.1 ± 0 | 148.4 ± 23 | 80.5 ± 5.7 | 339.6 ± 21.4 | 434.1 ± 31.3 | 121.8 ± 5.3 |
| | **T1/2/N2** | 81.7 ± 6.6 | 0.1 ± 0.1 | 0.2 ± 0 | 815.2 ± 34.1 | 6.1 ± 0.5 | 0 ± 0 | 0.7 ± 0.1 | 33.3 ± 0.4 | 12.6 ± 5.6 | 86.4 ± 6.6 | 276 ± 1.3 | 152.9 ± 4 |
| | **T1/2/T7** | 74.5 ± 29.6 | 0.1 ± 0.1 | 0.8 ± 0.3 | 1664 ± 185.9 | 13.9 ± 0.9 | 4.4 ± 2.7 | 3.1 ± 0.4 | 240.7 ± 8.1 | 107.2 ± 2 | 546.6 ± 40.5 | 748.6 ± 126.4 | 143.2 ± 2.3 |
| | **T7/RI** | 141.5 ± 5.9 | 0.1 ± 0 | 0.3 ± 0.1 | 328.8 ± 162.6 | 2 ± 1.1 | 0 ± 0 | 1.3 ± 0 | 80.3 ± 7.3 | 32.4 ± 1.9 | 161.3 ± 45.5 | 345.8 ± 89.7 | 98.5 ± 10.1 |
| | **N2/T7** | 60.9 ± 3.3 | 0.1 ± 0 | 0.5 ± 0 | 2441.4 ± 58.4 | 11.7 ± 0.1 | 3.7 ± 0.5 | 1.4 ± 0.1 | 236.7 ± 15.6 | 100 ± 10.3 | 433.2 ± 7.3 | 917.4 ± 75.1 | 131.9 ± 7.9 |

**Table S7**. Concentration of OVA-specific IgG, IgG1 and IgG2a antibody responses.

| | **Average ± SEM [AU/mL]** | | |
|---|---|---|---|
| | **Total IgG** | **IgG1** | **IgG2a** |
| **OVA** | 10.6 ± 0.6 | 161.6 ± 49.2 | 6.6 ± 1.2 |
| **OVA + ALUM** | 113.6 ± 49.2 | 922.8 ± 273.4 | 8.0 ± 1.2 |
| **OVA + T4/T7** | 1449.2 ± 231.4 | 10506.8 ± 2228.4 | 1106.0 ± 306 |
| **OVA + T1/2/N2** | 237.8 ± 18.9 | 2084.4 ± 409.2 | 49.0 ± 29.1 |
| **OVA + T1/2/T7** | 339.4 ± 52.6 | 2886.0 ± 554.6 | 12.8 ± 4.0 |
| **OVA + T7/RI** | 1289.4 ± 154.7 | 8488.6 ± 2916.2 | 1435.2 ± 420.1 |

**Table S8A**. Tumor volumes of B16F10 tumors.

| | Average ± SEM tumor volume (mm³) | | | | | | | | | |
|---|---|---|---|---|---|---|---|---|---|---|
| **Days after treatment initiation (DPT)** | **0** | **2** | **4** | **6** | **8** | **10** | **12** | **14** | **16** | **18** |
| **Vehicle** | 102.4 ± 11.3 | 133.5 ± 14.3 | 191.2 ± 22.6 | 341 ± 54.0 | 499.8 ± 59.2 | 701.9 ± 79.7 | 1086.7 ± 103.3 | 1454.5 ± 146.2 | 1886.1 ± 155.5 | 2287.6 ± 154.6 |
| **T4/T7** | 100.3 ± 9.1 | 130.5 ± 17.6 | 169.1 ± 24.6 | 295 ± 55.8 | 385.9 ± 78.1 | 501.4 ± 100.9 | 684 ± 149.8 | 910.6 ± 219.3 | 1079.7 ± 279.5 | 1132.4 ± 199.6 |
| **T7/RI** | 105.3 ± 15.4 | 129.2 ± 17.6 | 162.3 ± 21.4 | 282.3 ± 48.1 | 397.1 ± 70.8 | 575.8 ± 96.4 | 893.4 ± 138.2 | 1013.4 ± 141.3 | 1548 ± 215.9 | 2006.3 ± 295.8 |
| **Resiquimod** | 102.1 ± 9.6 | 136.1 ± 12.2 | 177.6 ± 16.8 | 232.3 ± 27.6 | 290.2 ± 25.9 | 351.2 ± 36.4 | 505.5 ± 59.2 | 723.5 ± 76.5 | 1038.9 ± 104.1 | 1236.2 ± 156.1 |

**Table S8B**. Calculated TGI and T/C values.

| | **TGI, %** | | | **T/C, %** | | |
|---|---|---|---|---|---|---|
| **Groups** | **maximal** | **DPT 16** | **DPT 18** | **lowest** | **DPT 16** | **DPT 18** |
| **T4/T7** | 58.95 (DPT 18) | 50.25 | 58.95 | 49.5 (DPT 18) | 57.25 | 49.50 |
| **T7/RI** | 32.1 (DPT 14) | 25.96 | 30.62 | 69.7 (DPT 16) | 69.67 | 87.70 |
| **Resiquimod** | 56.3 (DPT 18) | 48.99 | 56.27 | 54.0 (DPT 18) | 55.08 | 54.04 |

**Table S8C.** Survival of mice.

| **Groups** | **vehicle** | **T4/T7** | **T7/RI** | **resiquimod** |
|---|---|---|---|---|
| **Median survival [days]** | 18 | 22 | 20 | 24 |

### 5. HPLC chromatograms/traces of final compounds

Compound **T4/T7**

**Chromatogram and Results**

**Injection Details**

| | | | |
|---|---|---|---|
| *Injection Name:* | T4/T7 | *Run Time (min):* | 12,50 |
| *Vial Number:* | GB3 | *Injection Volume:* | |
| *Injection Type:* | | *Channel:* | UV_VIS_2 |
| *Calibration Level:* | | *Wavelength:* | 254 |
| *Instrument Method:* | Method2021 | *Bandwidth:* | 16 |
| *Processing Method:* | Quantitative | *Dilution Factor:* | 1,0000 |
| *Injection Date/Time:* | | *Sample Weight:* | 1,0000 |

**Chromatogram**

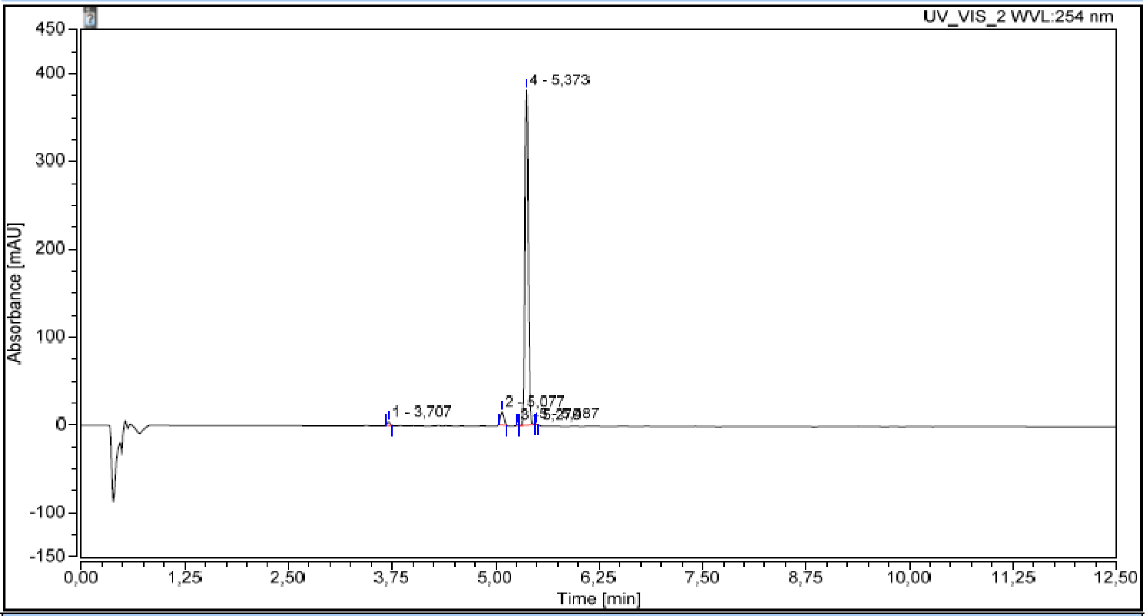

**Integration Results**

| No. | Peak Name | Retention Time min | Area mAU*min | Height mAU | Relative Area % | Relative Height % | Amount n.a. |
|---|---|---|---|---|---|---|---|
| 1 | | 3,707 | 0,162 | 4,452 | 0,87 | 1,11 | n.a. |
| 2 | | 5,077 | 0,710 | 15,064 | 3,83 | 3,74 | n.a. |
| 3 | | 5,270 | 0,025 | 0,857 | 0,13 | 0,21 | n.a. |
| 4 | | 5,373 | 17,606 | 381,538 | 95,03 | 94,72 | n.a. |
| 5 | | 5,487 | 0,024 | 0,904 | 0,13 | 0,22 | n.a. |
| Total: | | | 18,527 | 402,815 | 100,00 | 100,00 | |

Compound **T4/N2**

#### Chromatogram and Results

**Injection Details**

| | | | |
|---|---|---|---|
| *Injection Name:* | T4/N2 | *Run Time (min):* | 12,50 |
| *Vial Number:* | BB6 | *Injection Volume:* | |
| *Injection Type:* | | *Channel:* | UV_VIS_2 |
| *Calibration Level:* | | *Wavelength:* | 254 |
| *Instrument Method:* | Method2021 | *Bandwidth:* | 16 |
| *Processing Method:* | Quantitative | *Dilution Factor:* | 1,0000 |
| *Injection Date/Time:* | | *Sample Weight:* | 1,0000 |

**Chromatogram**

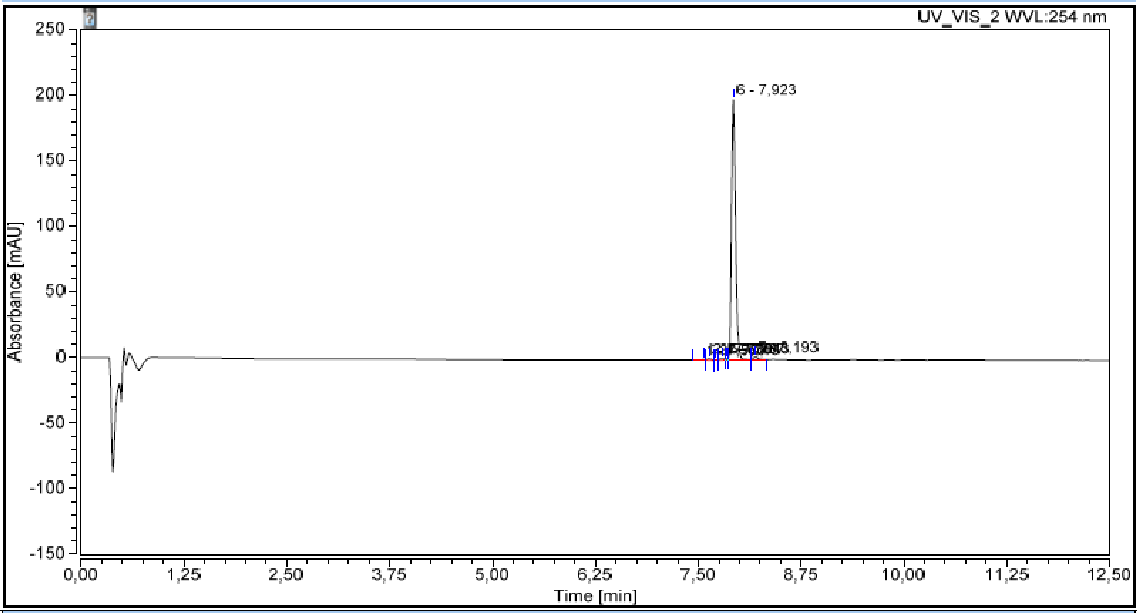

**Integration Results**

| No. | Peak Name | Retention Time min | Area mAU*min | Height mAU | Relative Area % | Relative Height % | Amount n.a. |
|---|---|---|---|---|---|---|---|
| 1 | | 7,560 | 0,013 | 0,242 | 0,12 | 0,12 | n.a. |
| 2 | | 7,630 | 0,060 | 1,149 | 0,52 | 0,56 | n.a. |
| 3 | | 7,713 | 0,008 | 0,190 | 0,07 | 0,09 | n.a. |
| 4 | | 7,787 | 0,074 | 1,102 | 0,65 | 0,54 | n.a. |
| 5 | | 7,843 | 0,026 | 0,991 | 0,23 | 0,48 | n.a. |
| 6 | | 7,923 | 11,184 | 198,616 | 97,17 | 97,17 | n.a. |
| 7 | | 8,193 | 0,144 | 2,121 | 1,25 | 1,04 | n.a. |
| **Total:** | | | **11,510** | **204,410** | **100,00** | **100,00** | |

Compound **T1/2/T4**

#### Chromatogram and Results

**Injection Details**

| | | | |
|---|---|---|---|
| *Injection Name:* | T1/2/T4 | *Run Time (min):* | 12,00 |
| *Vial Number:* | BB4 | *Injection Volume:* | |
| *Injection Type:* | | *Channel:* | UV_VIS_2 |
| *Calibration Level:* | | *Wavelength:* | 254 |
| *Instrument Method:* | Method2021_12 | *Bandwidth:* | 16 |
| *Processing Method:* | Quantitative | *Dilution Factor:* | 1,0000 |
| *Injection Date/Time:* | | *Sample Weight:* | 1,0000 |

**Chromatogram**

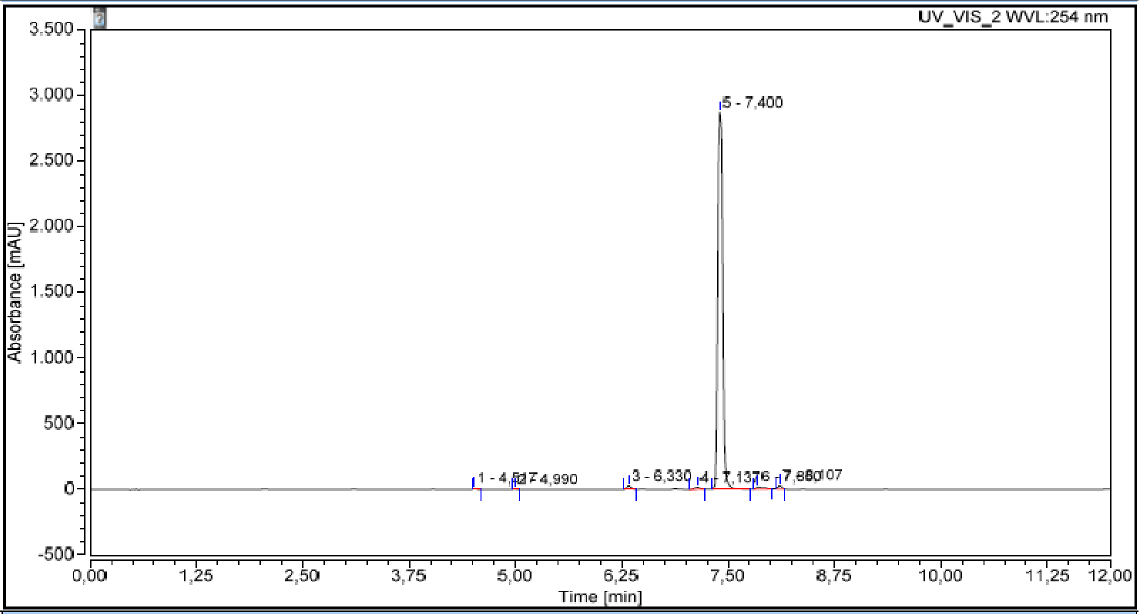

**Integration Results**

| No. | Peak Name | Retention Time min | Area mAU*min | Height mAU | Relative Area % | Relative Height % | Amount n.a. |
|---|---|---|---|---|---|---|---|
| 1 | | 4,517 | 0,310 | 7,433 | 0,15 | 0,25 | n.a. |
| 2 | | 4,990 | 0,110 | 2,537 | 0,05 | 0,09 | n.a. |
| 3 | | 6,330 | 1,573 | 28,336 | 0,76 | 0,95 | n.a. |
| 4 | | 7,137 | 1,030 | 14,844 | 0,50 | 0,50 | n.a. |
| 5 | | 7,400 | 201,222 | 2873,035 | 97,04 | 96,69 | n.a. |
| 6 | | 7,850 | 1,832 | 18,820 | 0,88 | 0,63 | n.a. |
| 7 | | 8,107 | 1,292 | 26,451 | 0,62 | 0,89 | n.a. |
| **Total:** | | | **207,369** | **2971,456** | **100,00** | **100,00** | |

Compound **T1/2/N2**

#### Chromatogram and Results

**Injection Details**

| | | | |
|---|---|---|---|
| *Injection Name:* | T1/2/N2 | *Run Time (min):* | 12,50 |
| *Vial Number:* | BB5 | *Injection Volume:* | |
| *Injection Type:* | | *Channel:* | UV_VIS_2 |
| *Calibration Level:* | | *Wavelength:* | 254 |
| *Instrument Method:* | Method2021 | *Bandwidth:* | 16 |
| *Processing Method:* | Quantitative | *Dilution Factor:* | 1,0000 |
| *Injection Date/Time:* | | *Sample Weight:* | 1,0000 |

**Chromatogram**

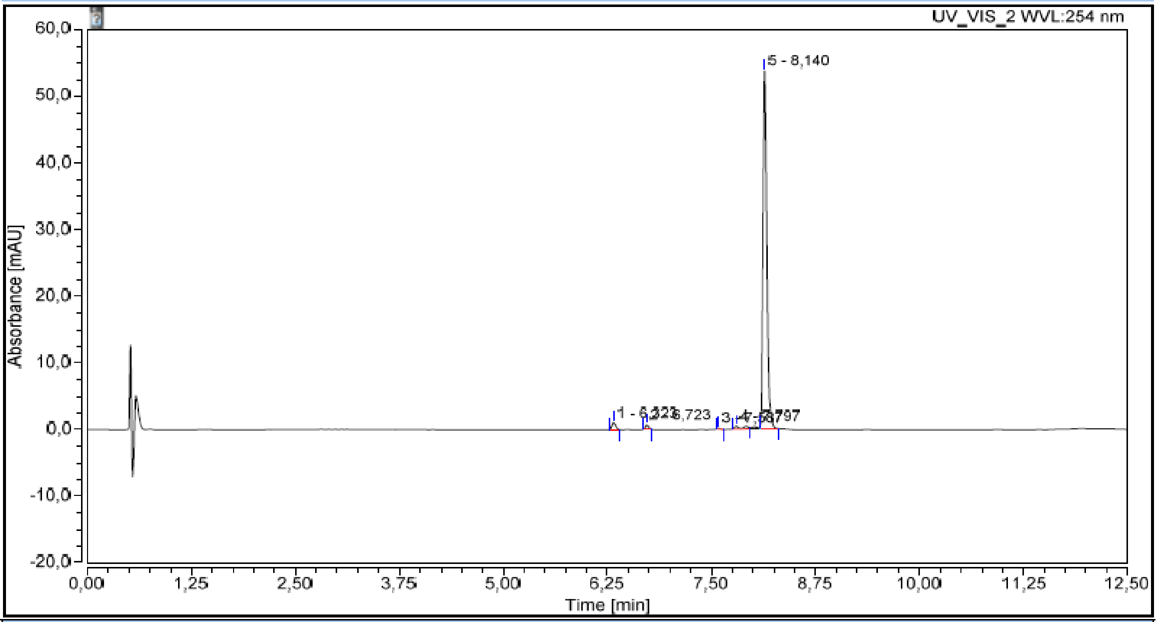

**Integration Results**

| No. | Peak Name | Retention Time<br>min | Area<br>mAU*min | Height<br>mAU | Relative Area<br>% | Relative Height<br>% | Amount<br>n.a. |
|---|---|---|---|---|---|---|---|
| 1 | | 6,323 | 0,057 | 1,065 | 1,86 | 1,90 | n.a. |
| 2 | | 6,723 | 0,031 | 0,683 | 1,02 | 1,22 | n.a. |
| 3 | | 7,587 | 0,005 | 0,141 | 0,18 | 0,25 | n.a. |
| 4 | | 7,797 | 0,056 | 0,495 | 1,83 | 0,88 | n.a. |
| 5 | | 8,140 | 2,919 | 53,793 | 95,11 | 95,76 | n.a. |
| Total: | | | 3,069 | 56,177 | 100,00 | 100,00 | |

Compound **T1/2/T7**

#### Chromatogram and Results

**Injection Details**

| | | | |
|---|---|---|---|
| *Injection Name:* | **T1/2/T7** | *Run Time (min):* | **12,00** |
| *Vial Number:* | **BB2** | *Injection Volume:* | |
| *Injection Type:* | | *Channel:* | **UV_VIS_2** |
| *Calibration Level:* | | *Wavelength:* | **254** |
| *Instrument Method:* | **Method2021_12** | *Bandwidth:* | **16** |
| *Processing Method:* | **Quantitative(1)** | *Dilution Factor:* | **1,0000** |
| *Injection Date/Time:* | | *Sample Weight:* | **1,0000** |

**Chromatogram**

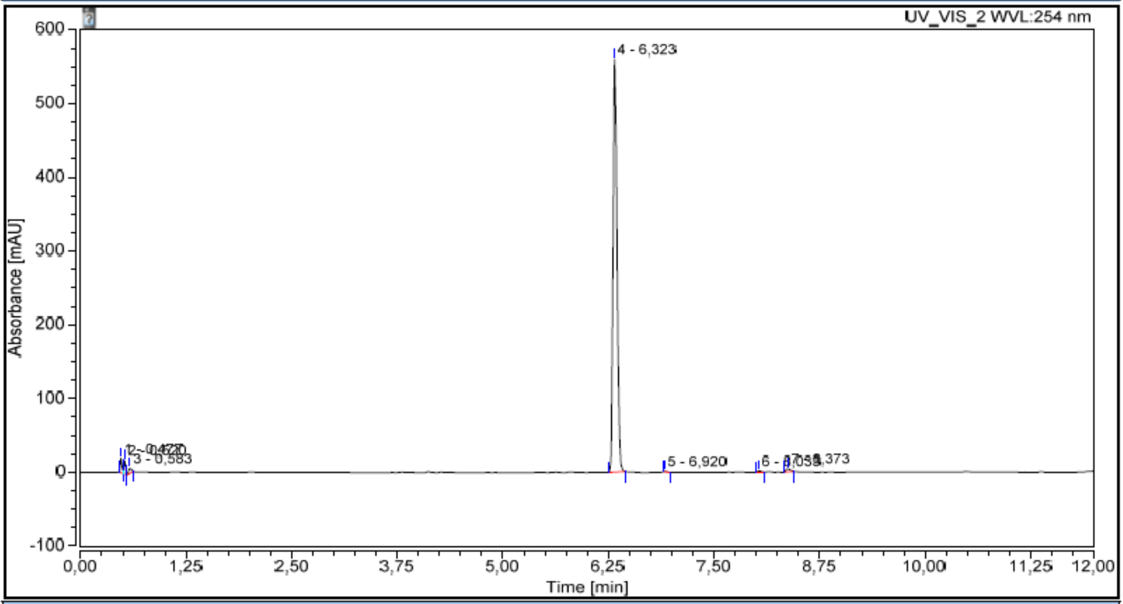

**Integration Results**

| No. | Peak Name | Retention Time<br>min | Area<br>mAU*min | Height<br>mAU | Relative Area<br>% | Relative Height<br>% | Amount<br>n.a. |
|---|---|---|---|---|---|---|---|
| 1 | | 0,477 | 0,446 | 19,223 | 1,38 | 3,15 | n.a. |
| 2 | | 0,520 | 0,466 | 18,287 | 1,44 | 3,00 | n.a. |
| 3 | | 0,583 | 0,300 | 6,747 | 0,93 | 1,11 | n.a. |
| 4 | | 6,323 | 30,759 | 559,644 | 95,09 | 91,66 | n.a. |
| 5 | | 6,920 | 0,029 | 0,763 | 0,09 | 0,12 | n.a. |
| 6 | | 8,033 | 0,090 | 1,626 | 0,28 | 0,27 | n.a. |
| 7 | | 8,373 | 0,257 | 4,243 | 0,79 | 0,69 | n.a. |
| **Total:** | | | **32,347** | **610,532** | **100,00** | **100,00** | |

Compound **T7/RI**

#### Chromatogram and Results

**Injection Details**

| | | | |
|---|---|---|---|
| *Injection Name:* | **T7/RI** | *Run Time (min):* | **8,00** |
| *Vial Number:* | **RD2** | *Injection Volume:* | **5,00** |
| *Injection Type:* | **Unknown** | *Channel:* | **UV_VIS_1** |
| *Calibration Level:* | | *Wavelength:* | **210** |
| *Instrument Method:* | **Method2021** | *Bandwidth:* | **8** |
| *Processing Method:* | **Quantitative** | *Dilution Factor:* | **1,0000** |
| *Injection Date/Time:* | | *Sample Weight:* | **1,0000** |

**Chromatogram**

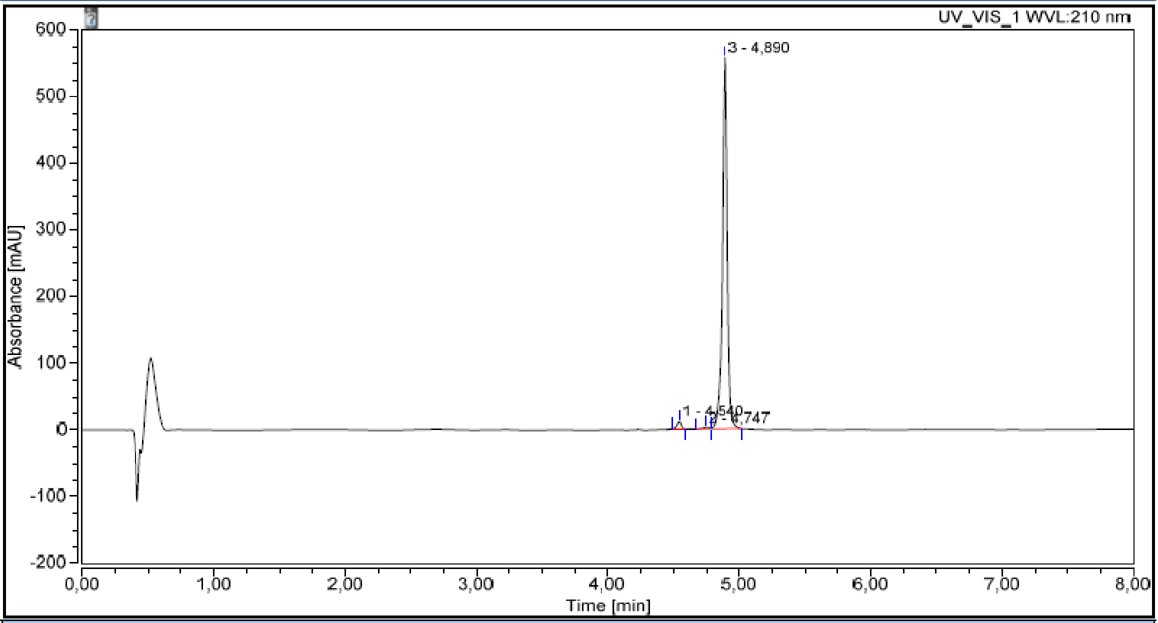

**Integration Results**

| No. | Peak Name | Retention Time min | Area mAU*min | Height mAU | Relative Area % | Relative Height % | Amount n.a. |
|---|---|---|---|---|---|---|---|
| 1 | | 4,540 | 0,490 | 12,510 | 1,97 | 2,19 | n.a. |
| 2 | | 4,747 | 0,146 | 2,320 | 0,59 | 0,41 | n.a. |
| 3 | | 4,890 | 24,201 | 557,444 | 97,44 | 97,41 | n.a. |
| Total: | | | **24,837** | **572,274** | **100,00** | **100,00** | |

Compound **T4/RI**

#### Chromatogram and Results

**Injection Details**

| | | | |
|---|---|---|---|
| *Injection Name:* | **T4/RI** | *Run Time (min):* | **12,50** |
| *Vial Number:* | **GB4** | *Injection Volume:* | |
| *Injection Type:* | | *Channel:* | **UV_VIS_2** |
| *Calibration Level:* | | *Wavelength:* | **254** |
| *Instrument Method:* | **Method2021** | *Bandwidth:* | **16** |
| *Processing Method:* | **Quantitative** | *Dilution Factor:* | **1,0000** |
| *Injection Date/Time:* | | *Sample Weight:* | **1,0000** |

**Chromatogram**

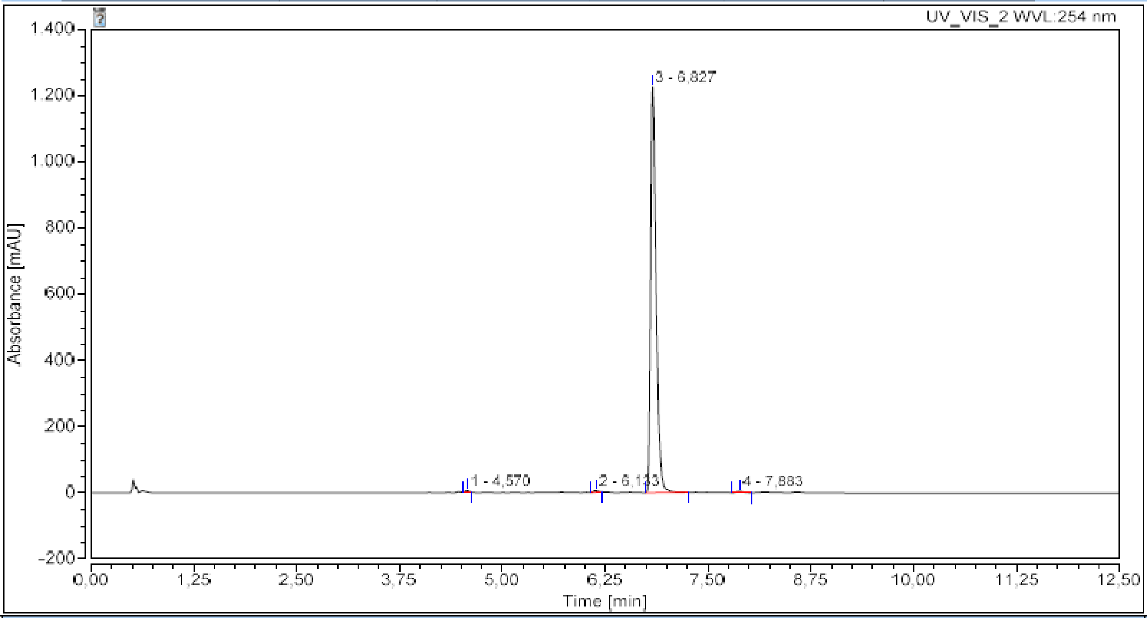

**Integration Results**

| No. | Peak Name | Retention Time min | Area mAU*min | Height mAU | Relative Area % | Relative Height % | Amount n.a. |
|---|---|---|---|---|---|---|---|
| 1 | | 4,570 | 0,267 | 5,407 | 0,28 | 0,43 | n.a. |
| 2 | | 6,133 | 0,324 | 5,158 | 0,34 | 0,41 | n.a. |
| 3 | | 6,827 | 94,781 | 1228,734 | 99,01 | 98,78 | n.a. |
| 4 | | 7,883 | 0,360 | 4,642 | 0,38 | 0,37 | n.a. |
| **Total:** | | | **95,732** | **1243,941** | **100,00** | **100,00** | |

Compound **T1/2/RI**

#### Chromatogram and Results

**Injection Details**

| | | | |
|---|---|---|---|
| *Injection Name:* | T1/2/RI | *Run Time (min):* | 12,50 |
| *Vial Number:* | GB6 | *Injection Volume:* | |
| *Injection Type:* | | *Channel:* | UV_VIS_2 |
| *Calibration Level:* | | *Wavelength:* | 254 |
| *Instrument Method:* | Method2021 | *Bandwidth:* | 16 |
| *Processing Method:* | Quantitative | *Dilution Factor:* | 1,0000 |
| *Injection Date/Time:* | | *Sample Weight:* | 1,0000 |

**Chromatogram**

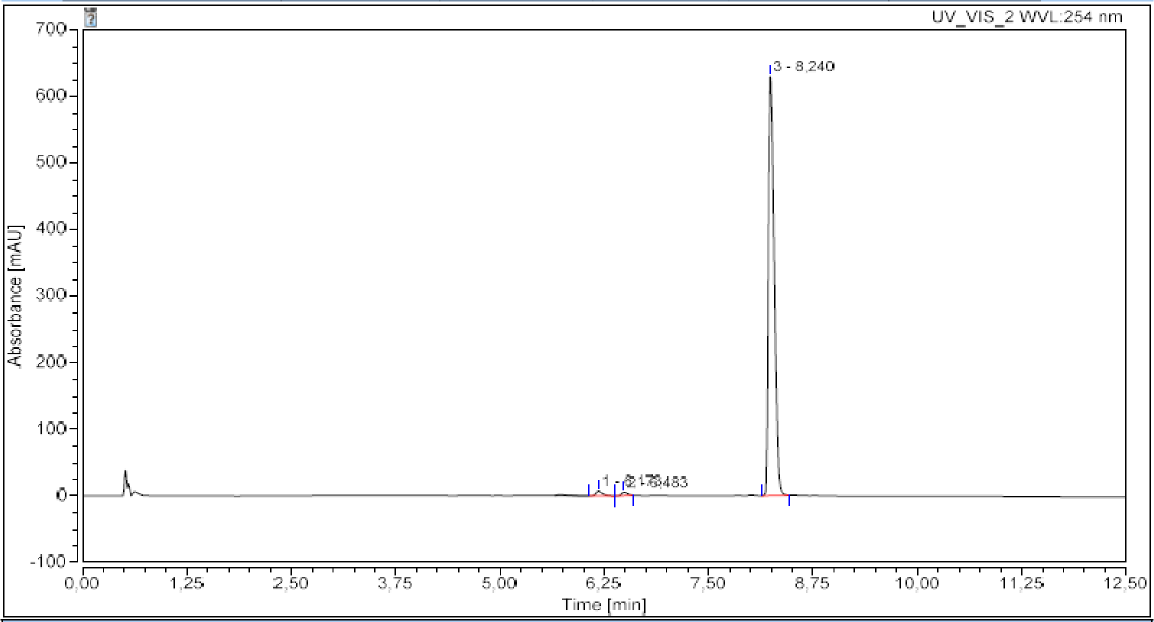

**Integration Results**

| No. | Peak Name | Retention Time min | Area mAU*min | Height mAU | Relative Area % | Relative Height % | Amount n.a. |
|---|---|---|---|---|---|---|---|
| 1 | | 6,173 | 0,709 | 7,762 | 1,34 | 1,21 | n.a. |
| 2 | | 6,483 | 0,445 | 5,486 | 0,84 | 0,85 | n.a. |
| 3 | | 8,240 | 51,653 | 630,243 | 97,82 | 97,94 | n.a. |
| Total: | | | 52,807 | 643,491 | 100,00 | 100,00 | |

Compound **N2/RI**

#### Chromatogram and Results

**Injection Details**

| | | | |
|---|---|---|---|
| *Injection Name:* | N2/RI | *Run Time (min):* | 12,50 |
| *Vial Number:* | BB3 | *Injection Volume:* | |
| *Injection Type:* | | *Channel:* | UV_VIS_2 |
| *Calibration Level:* | | *Wavelength:* | 254 |
| *Instrument Method:* | Method2021 | *Bandwidth:* | 16 |
| *Processing Method:* | Quantitative | *Dilution Factor:* | 1,0000 |
| *Injection Date/Time:* | | *Sample Weight:* | 1,0000 |

**Chromatogram**

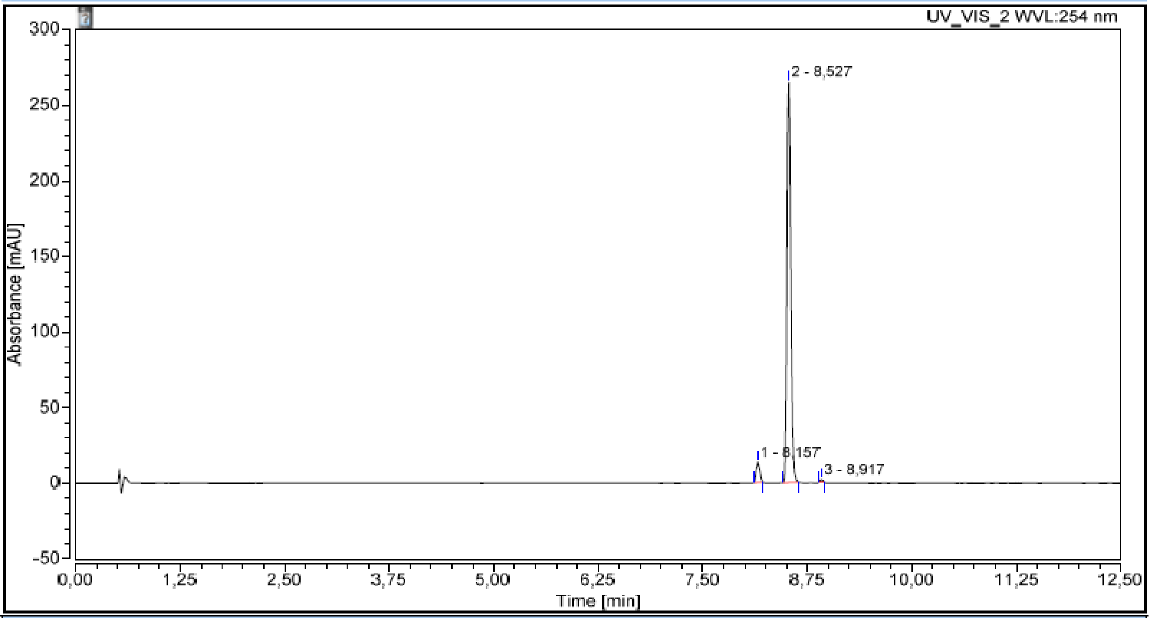

**Integration Results**

| No. | Peak Name | Retention Time min | Area mAU*min | Height mAU | Relative Area % | Relative Height % | Amount n.a. |
|---|---|---|---|---|---|---|---|
| 1 | | 8,157 | 0,680 | 13,456 | 4,52 | 4,80 | n.a. |
| 2 | | 8,527 | 14,315 | 264,958 | 95,04 | 94,59 | n.a. |
| 3 | | 8,917 | 0,066 | 1,692 | 0,44 | 0,60 | n.a. |
| Total: | | | 15,062 | 280,105 | 100,00 | 100,00 | |

### 6. HRMS spectra of final compounds

Compound **T4/T7**

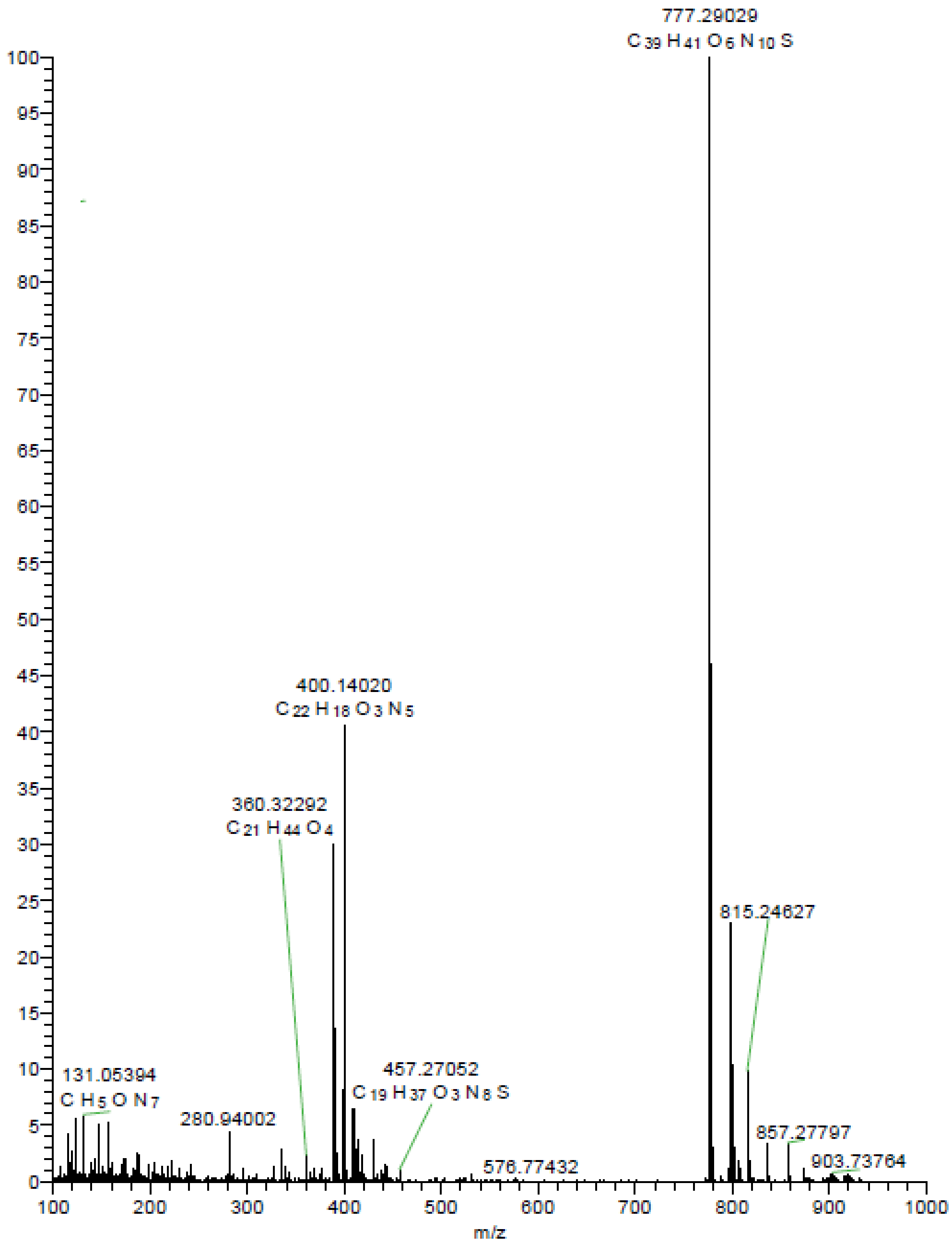

Elemental composition search on mass 777.29029

m/z= 772.29029-782.29029

| m/z | Theo. Mass | Delta (ppm) | RDB equiv. | Composition |
|---|---|---|---|---|
| 777.29029 | 777.29258 | -2.94 | 24.5 | $C_{39}H_{41}O_6N_{10}S$ |

Compound **T4/N2**

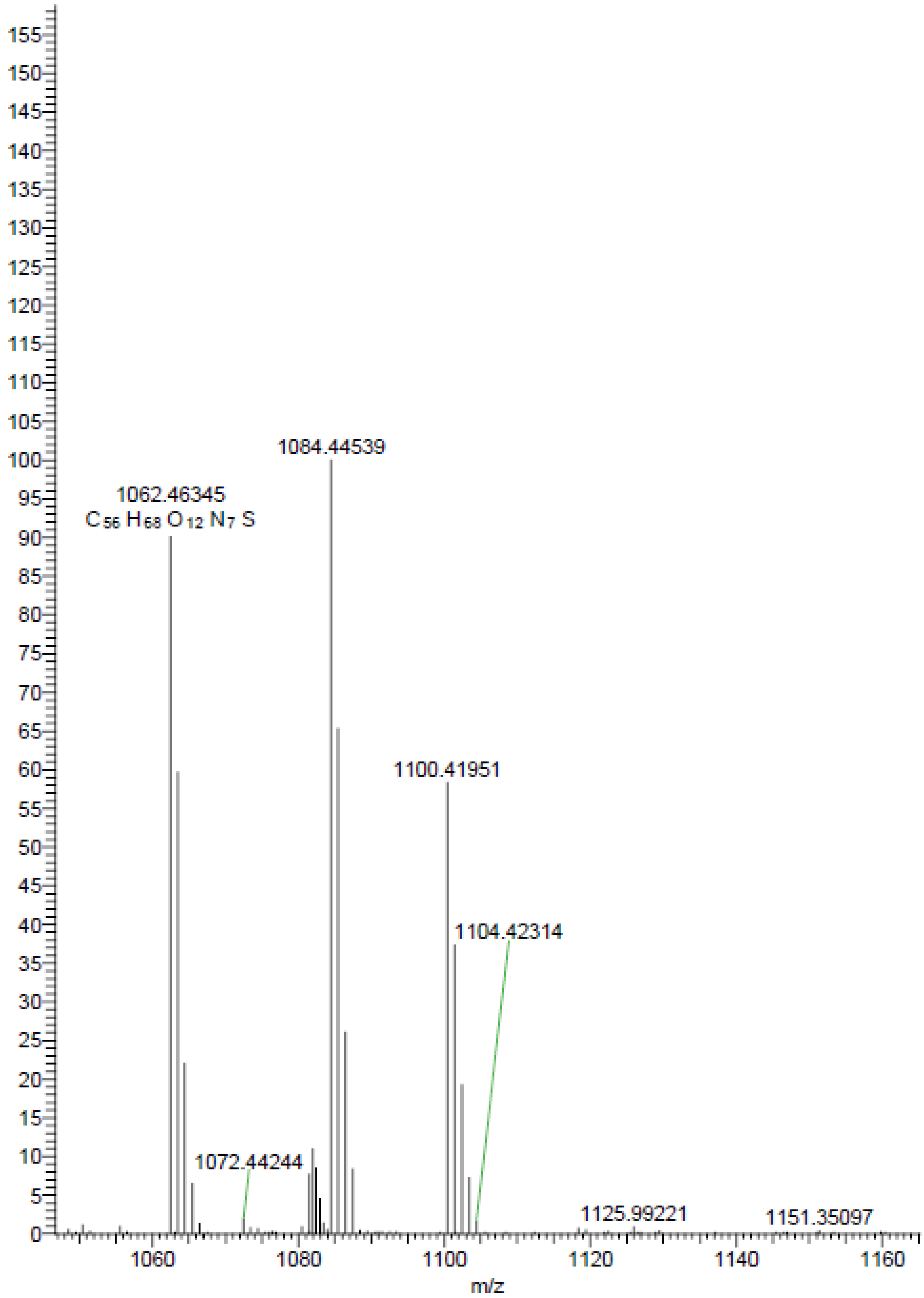

Elemental composition search on mass 1062.46345

m/z= 1057.46345-1067.46345

| m/z | Theo. Mass | Delta (ppm) | RDB equiv. | Composition |
|---|---|---|---|---|
| 1062.46345 | 1062.46412 | -0.63 | 26.5 | $C_{56}H_{68}O_{12}N_{7}S$ |

Compound **T1/2/T4**

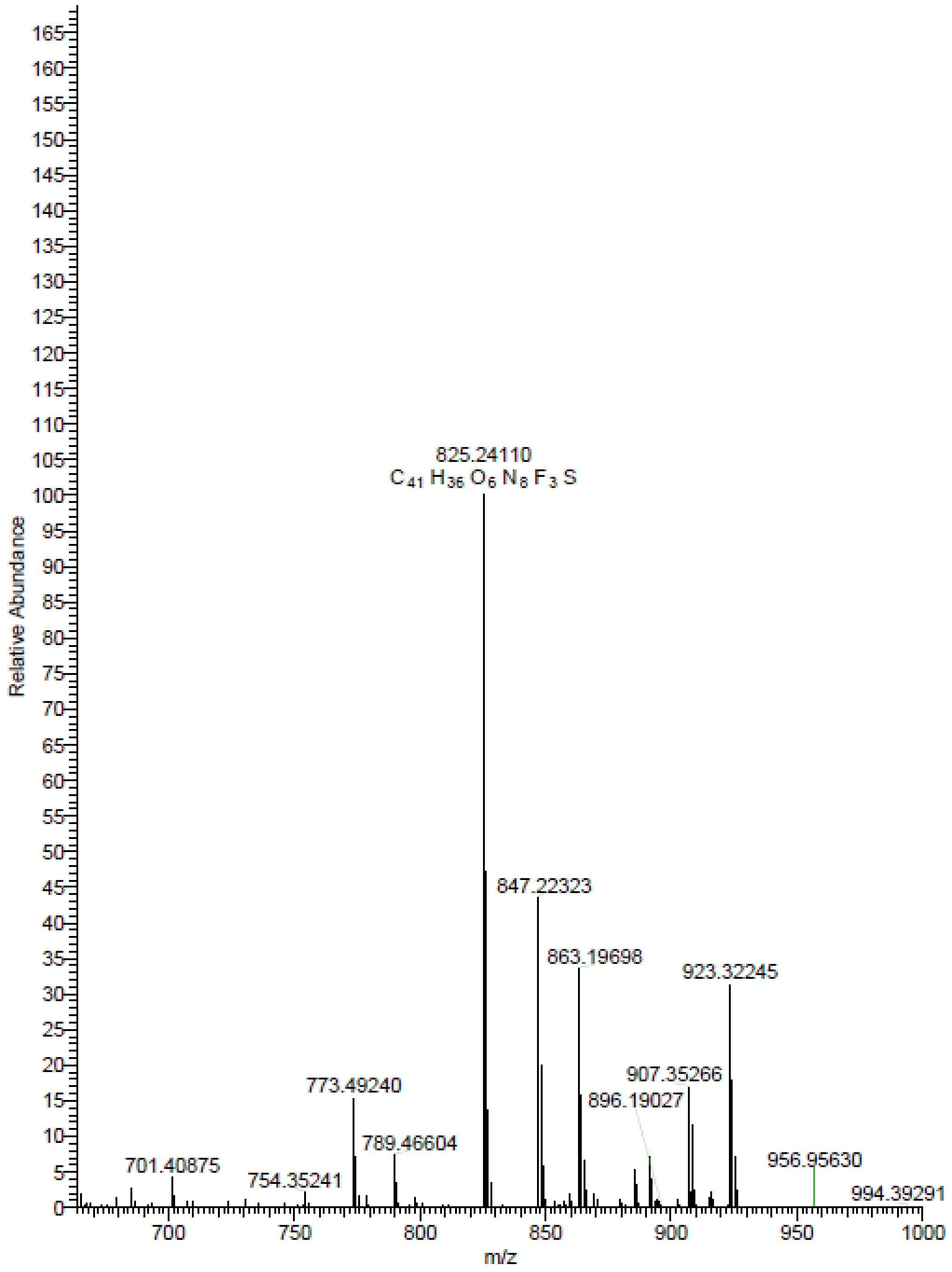

Elemental composition search on mass 825.24110

m/z= 820.24110-830.24110

| m/z | Theo. Mass | Delta (ppm) | RDB equiv. | Composition |
|---|---|---|---|---|
| 825.24110 | 825.24251 | -1.71 | 26.5 | $C_{41}H_{36}O_6N_8F_3S$ |

Compound **T1/2/N2**

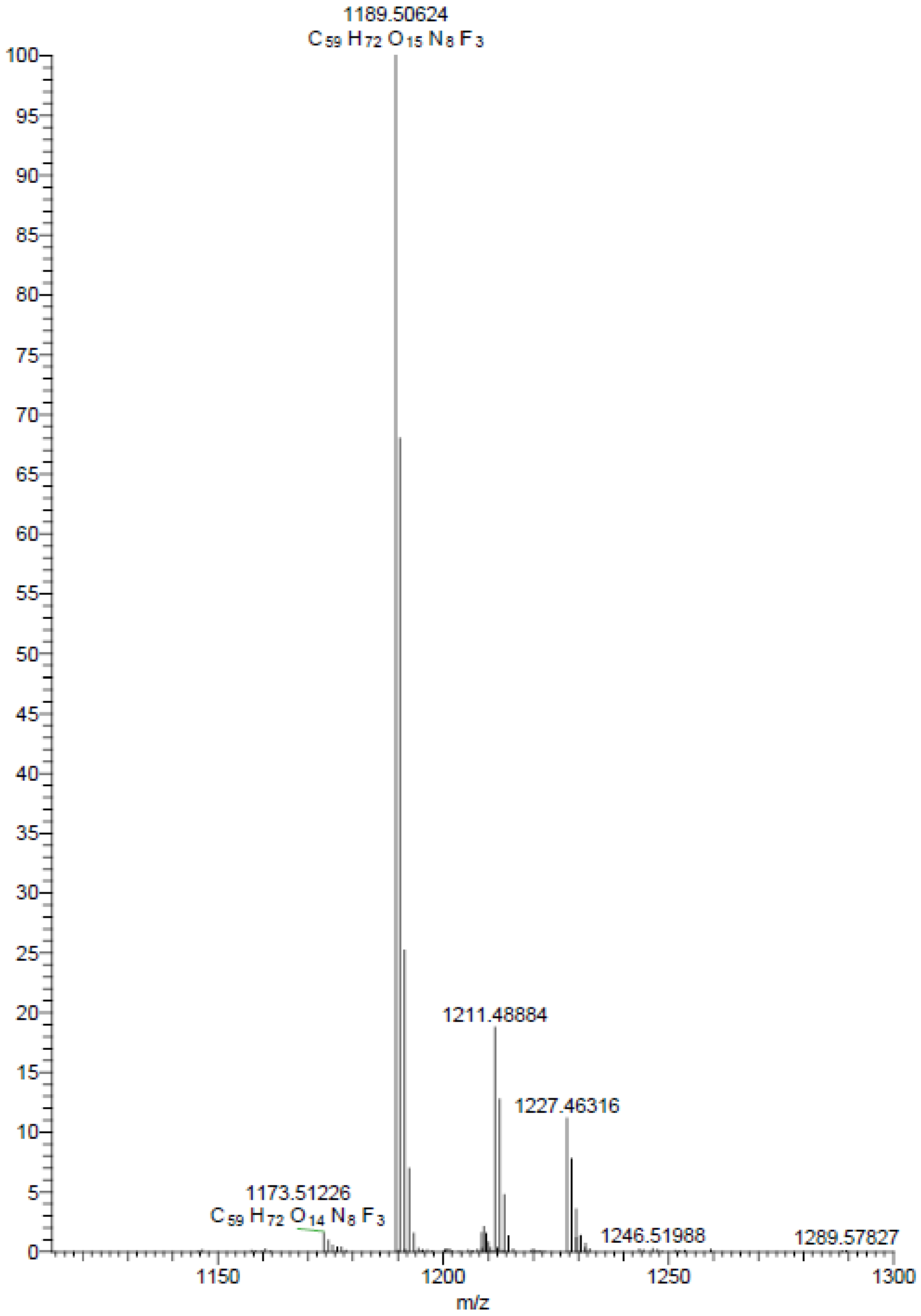

Elemental composition search on mass 1189.50624

m/z= 1184.50624-1194.50624

| m/z | Theo. Mass | Delta (ppm) | RDB equiv. | Composition |
|---|---|---|---|---|
| 1189.50624 | 1189.50637 | -0.11 | 26.5 | $C_{59}H_{72}O_{15}N_8F_3$ |

Compound **T1/2/T7**

SB-67 #177-361 RT: 0.78-1.59 AV: 185 NL: 6.66E6
T: FTMS + p ESI Full ms [150.0000-1200.0000]

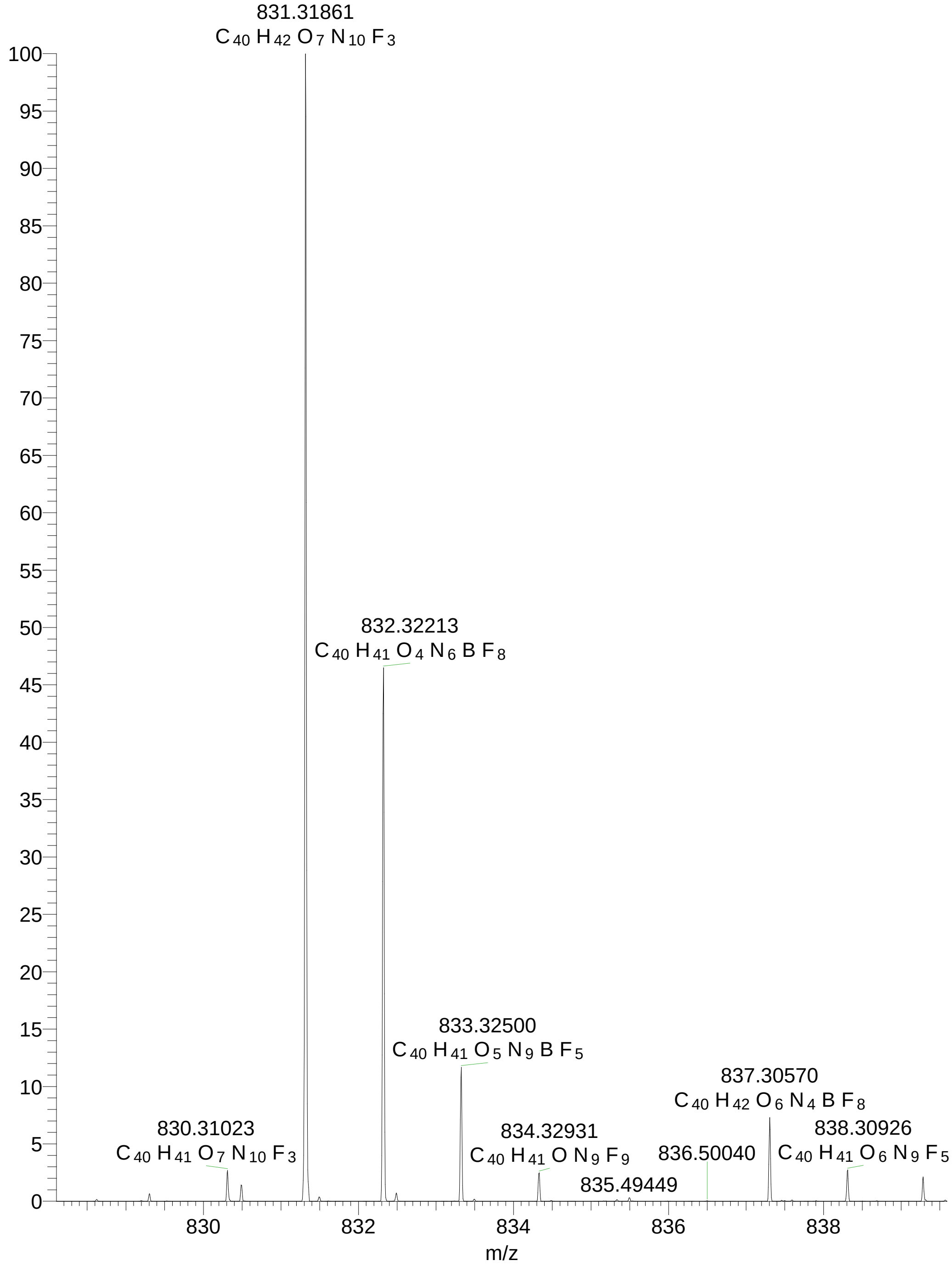

Elemental composition search on mass 831.31861

m/z= 826.31861-836.31861

| m/z | Theo. Mass | Delta (ppm) | RDB equiv. | Composition |
|---|---|---|---|---|
| 831.3186 | 831.3185 | 0.19 | 23.5 | $C_{40}H_{42}O_7N_{10}F_3$ |
| | [illegible] | [illegible] | [illegible] | [illegible] |

Compound **T7/RI**

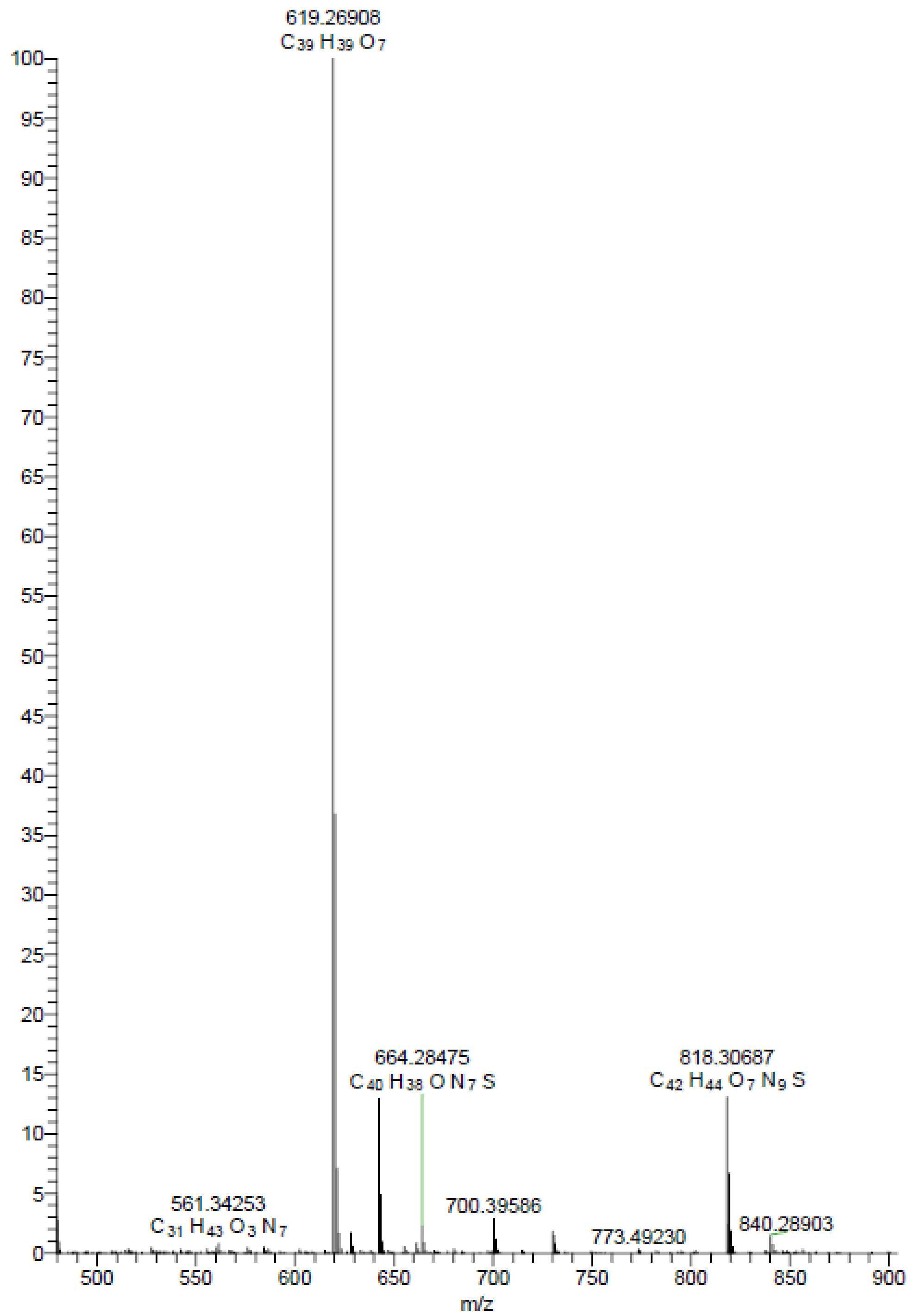

Elemental composition search on mass 818.30687

m/z= 813.30687-823.30687

| m/z | Theo. Mass | Delta (ppm) | RDB equiv. | Composition |
|---|---|---|---|---|
| 818.30687 | 818.30789 | -1.25 | 25.5 | $C_{42}H_{44}O_7N_9S$ |

Compound **T4/RI**

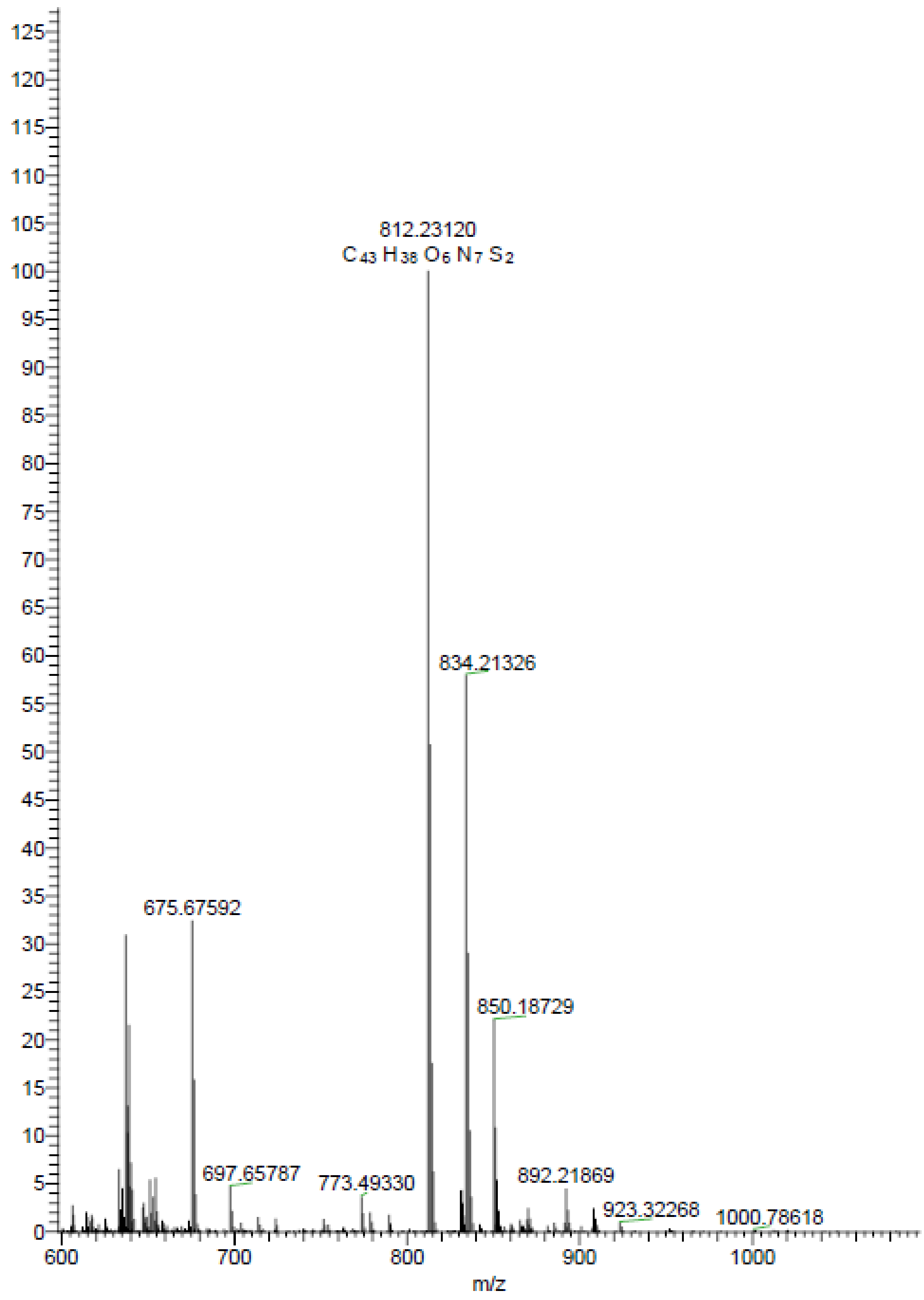

Elemental composition search on mass 812.23120

m/z= 807.23120-817.23120

| m/z | Theo. Mass | Delta (ppm) | RDB equiv. | Composition |
|---|---|---|---|---|
| 812.23120 | 812.23195 | -0.92 | 28.5 | $C_{43}H_{38}O_6N_7S_2$ |

Compound **T1/2/RI**

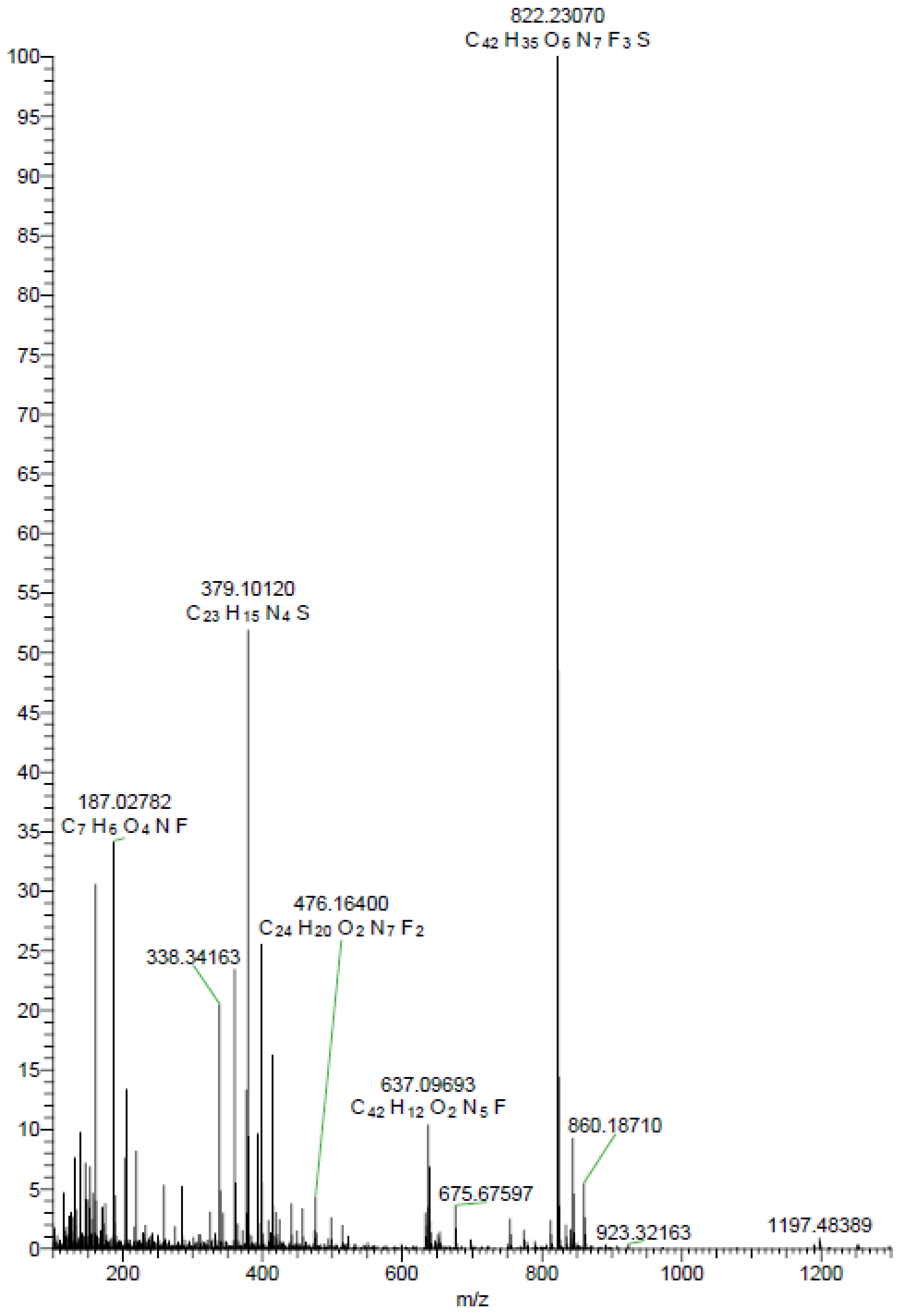

Elemental composition search on mass 822.23070

m/z= 817.23070-827.23070

| m/z | Theo. Mass | Delta (ppm) | RDB equiv. | Composition |
|---|---|---|---|---|
| 822.23070 | 822.23161 | -1.11 | 27.5 | $C_{42}H_{35}O_6N_7F_3S$ |

Compound **N2/RI**

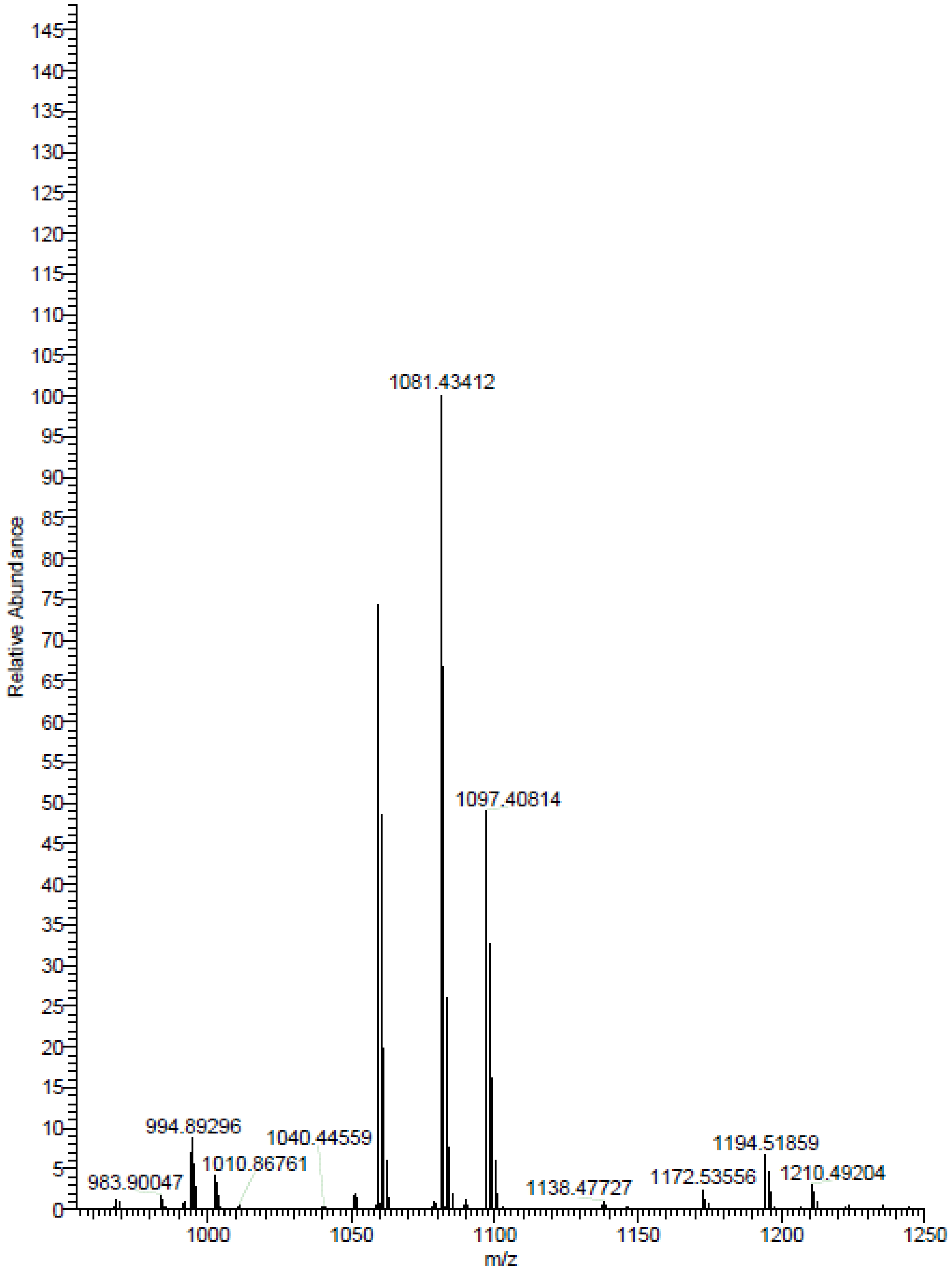

Elemental composition search on mass 1059.45239

m/z= 1054.45239-1064.45239

| m/z | Theo. Mass | Delta (ppm) | RDB equiv. | Composition |
|---|---|---|---|---|
| 1059.45239 | 1059.45322 | -0.78 | 27.5 | $C_{57} H_{67} O_{12} N_{6} S$ |

### 7. 1H and 13C NMR spectra of final compounds

Compound **T4/T7:** $^{1}H$, 400 MHz, DMSO-$d_6$

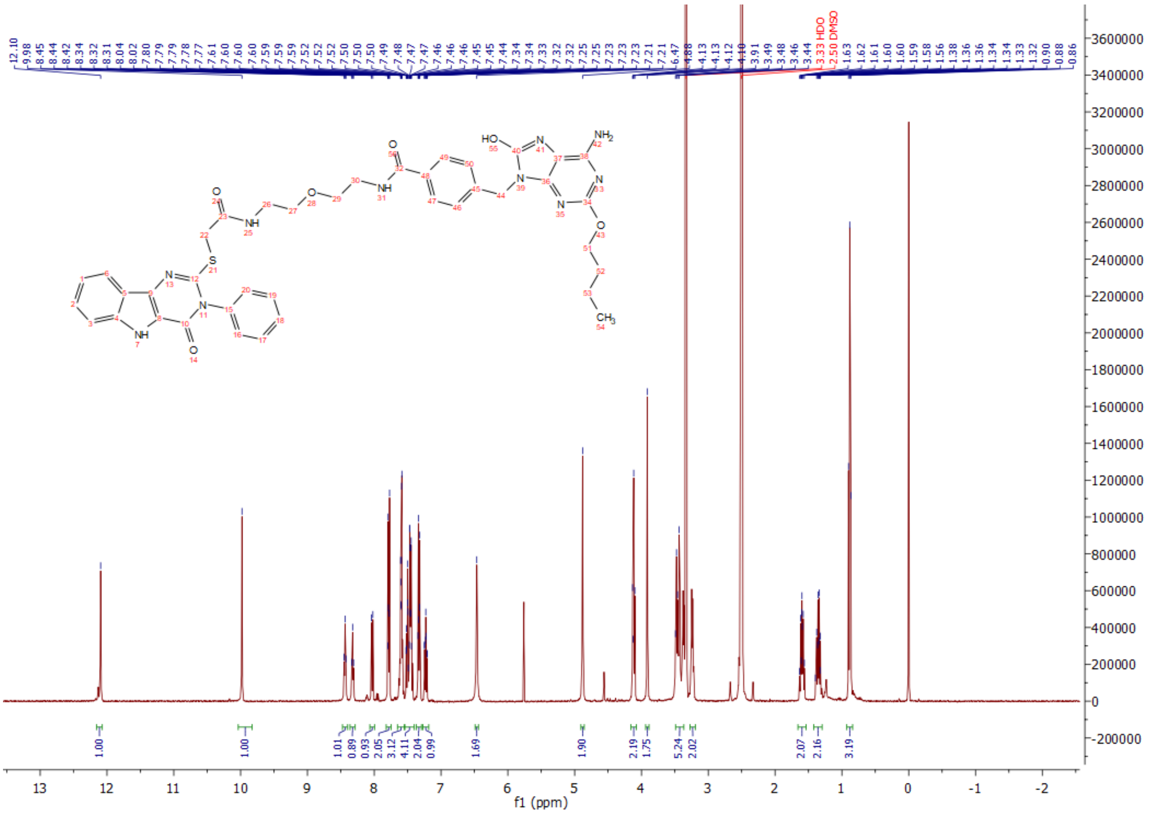

Compound **T4/T7:** $^{13}C$, 100 MHz, DMSO-$d_6$

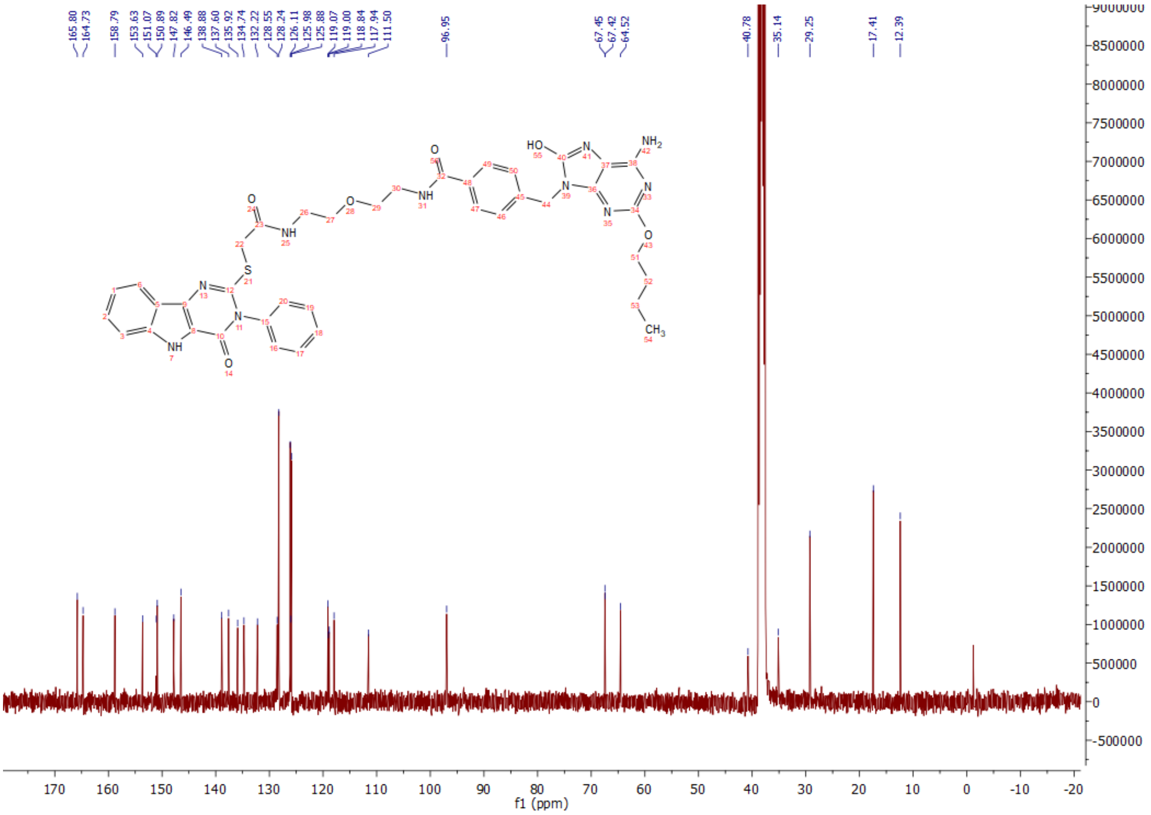

Compound **T4/N2:** $^{1}H$, 400 MHz, $CDCl_3$

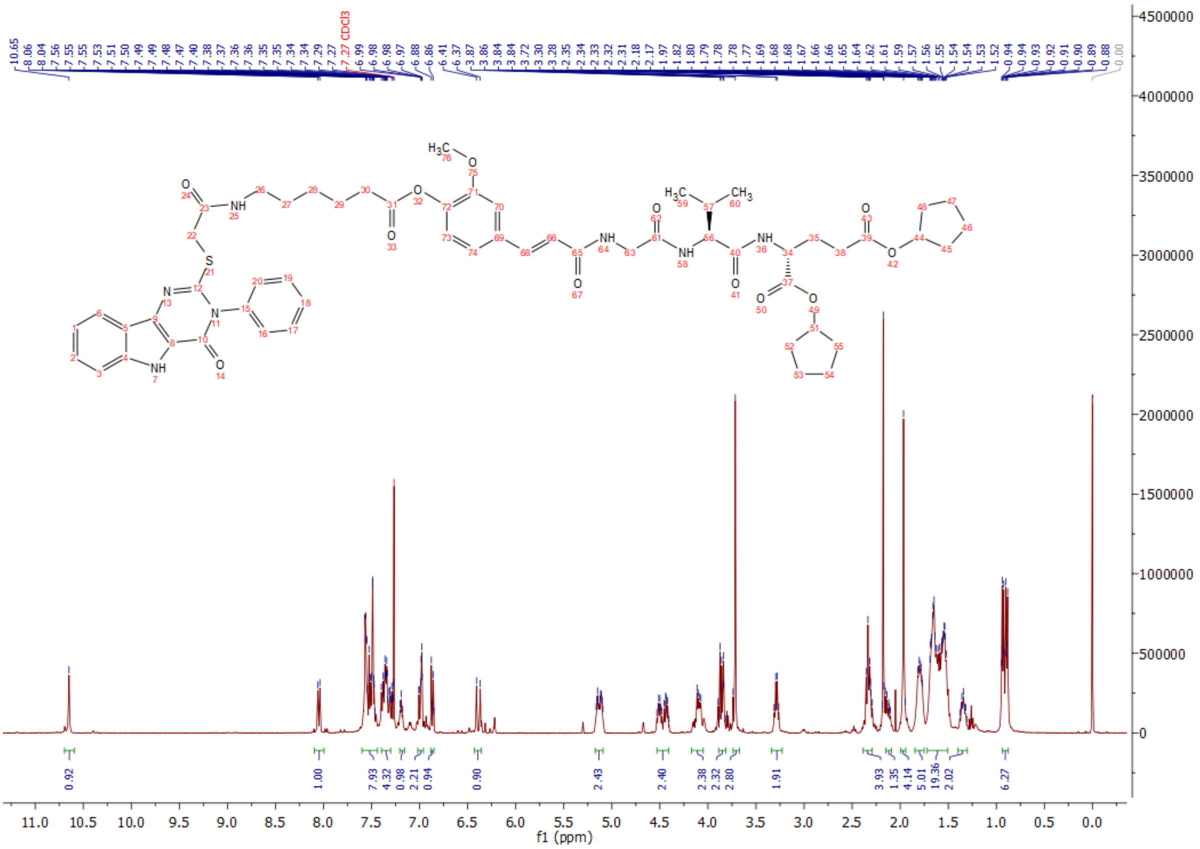

Compound **T4/N2:** $^{13}C$, 100 MHz, $CDCl_3$

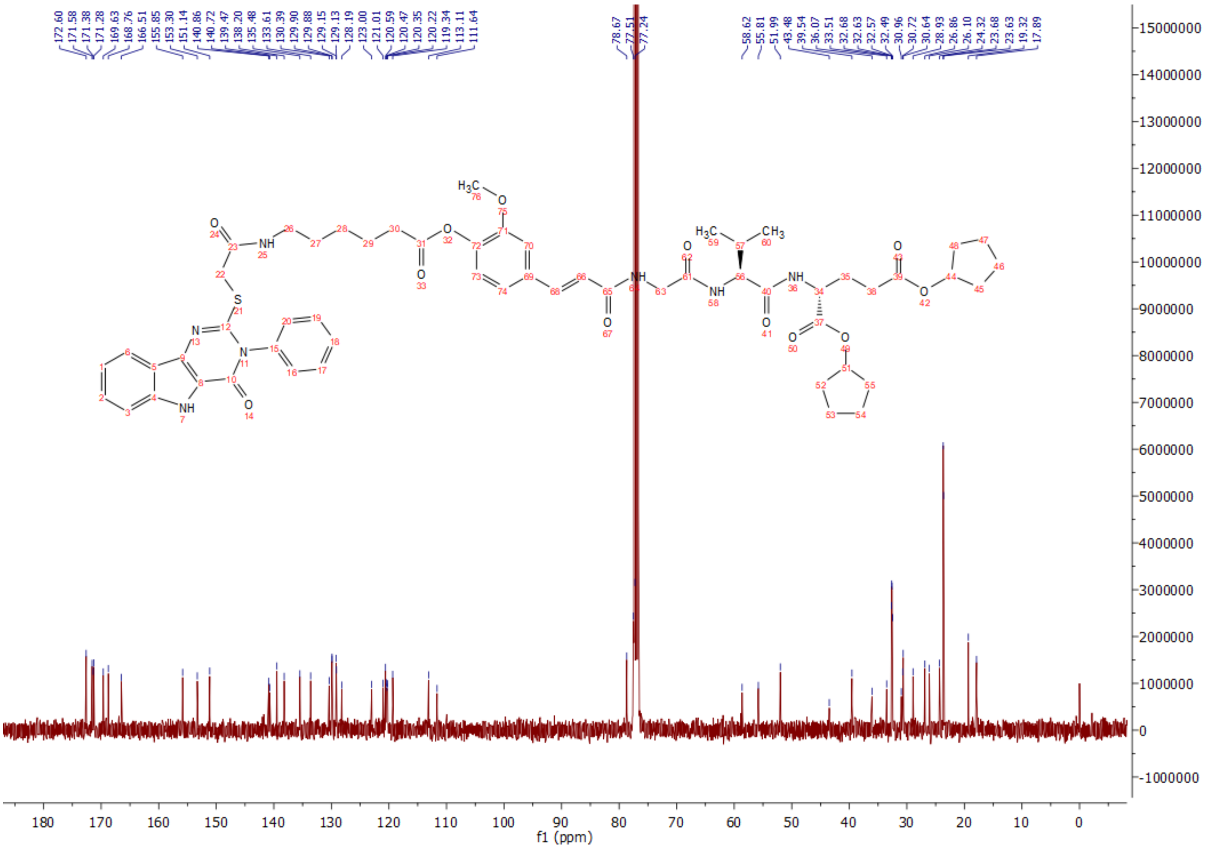

Compound **T1/2/T4:** $^{1}H$, 400 MHz, DMSO-$d_6$

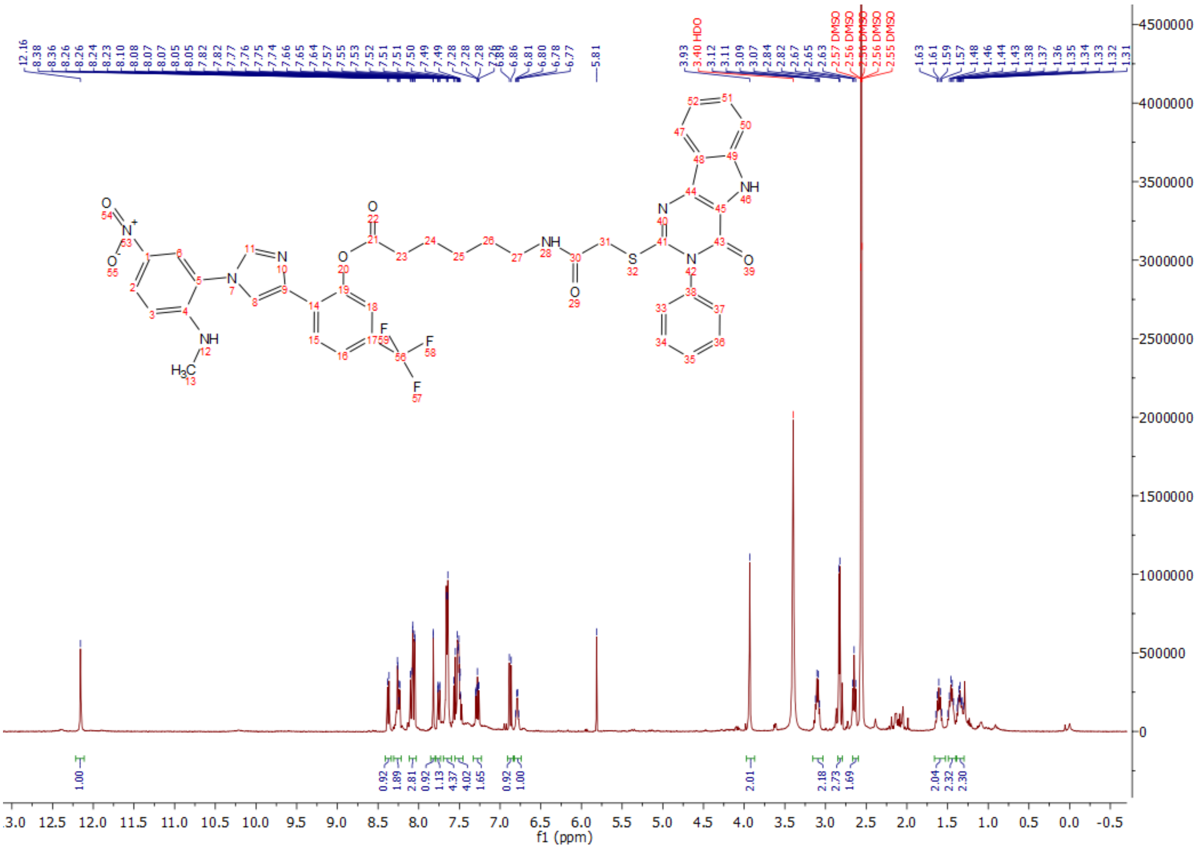

Compound **T1/2/T4:** $^{13}C$, 100 MHz, DMSO-$d_6$

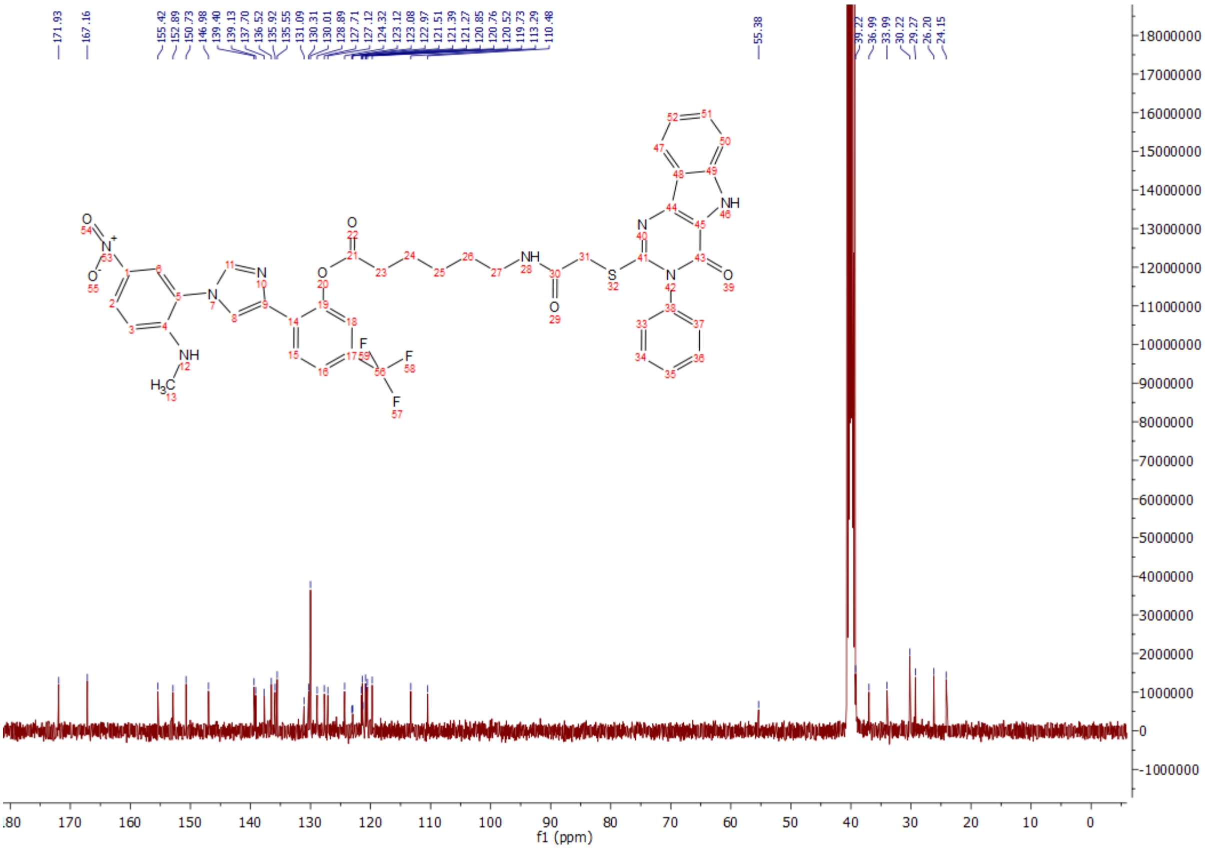

Compound **T1/2/N2:** $^{1}H$, 400 MHz, DMSO-d$_6$

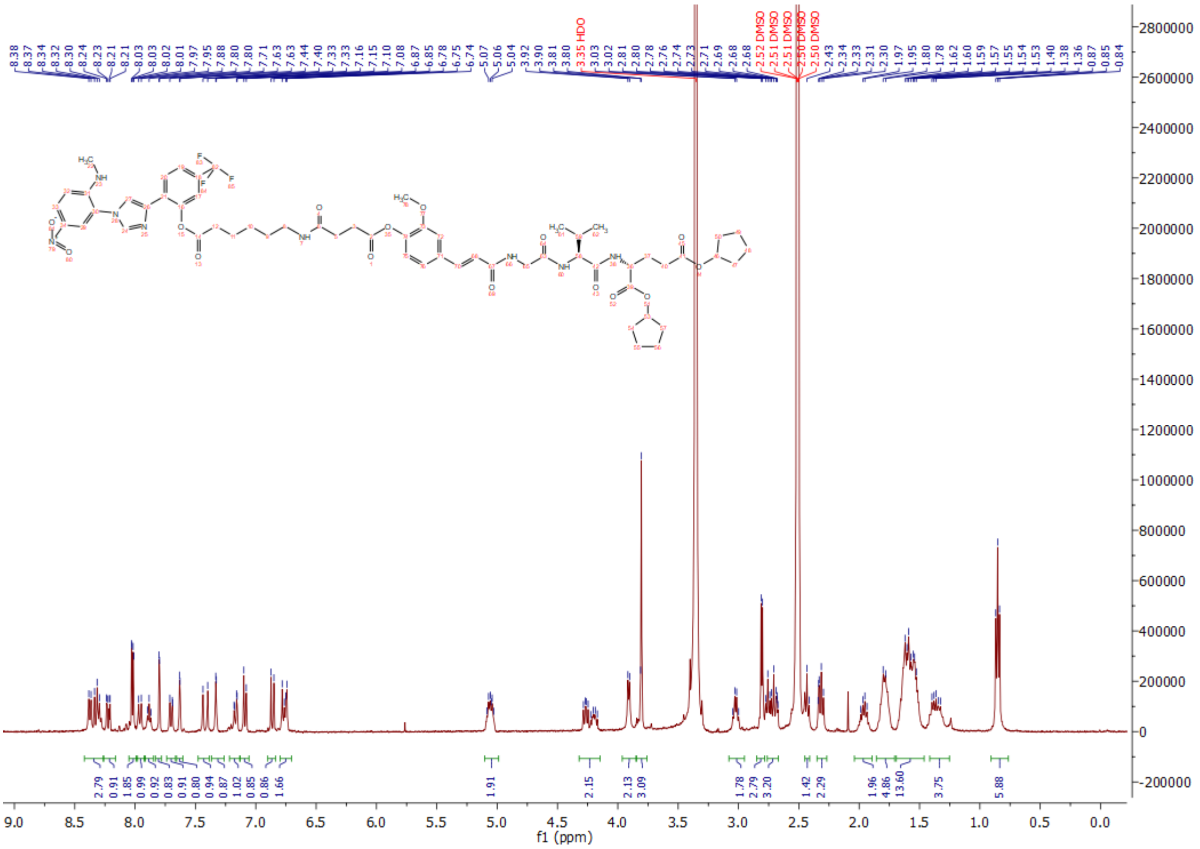

Compound **T1/2/N2:** $^{13}C$, 100 MHz, DMSO-d$_6$

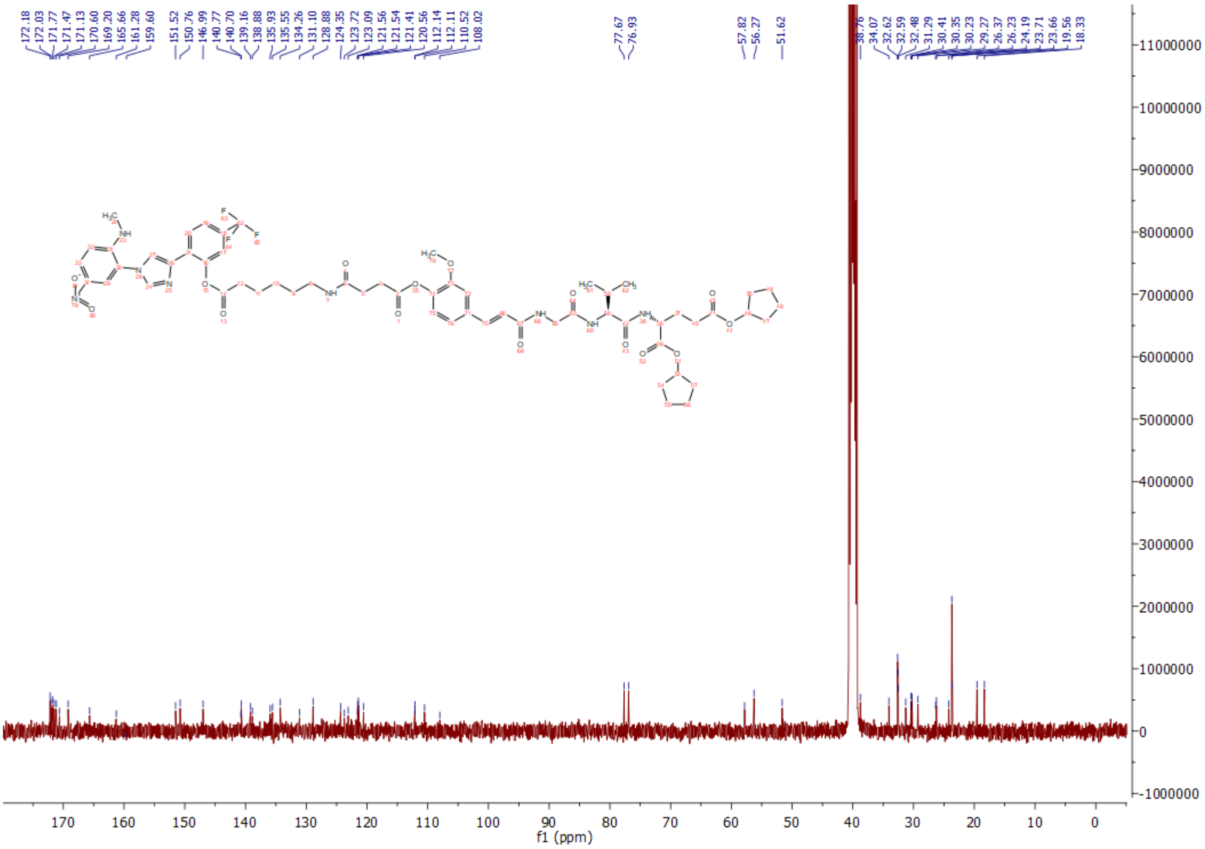

Compound **T1/2/T7:** $^{1}H$, 400 MHz, DMSO-d$_6$

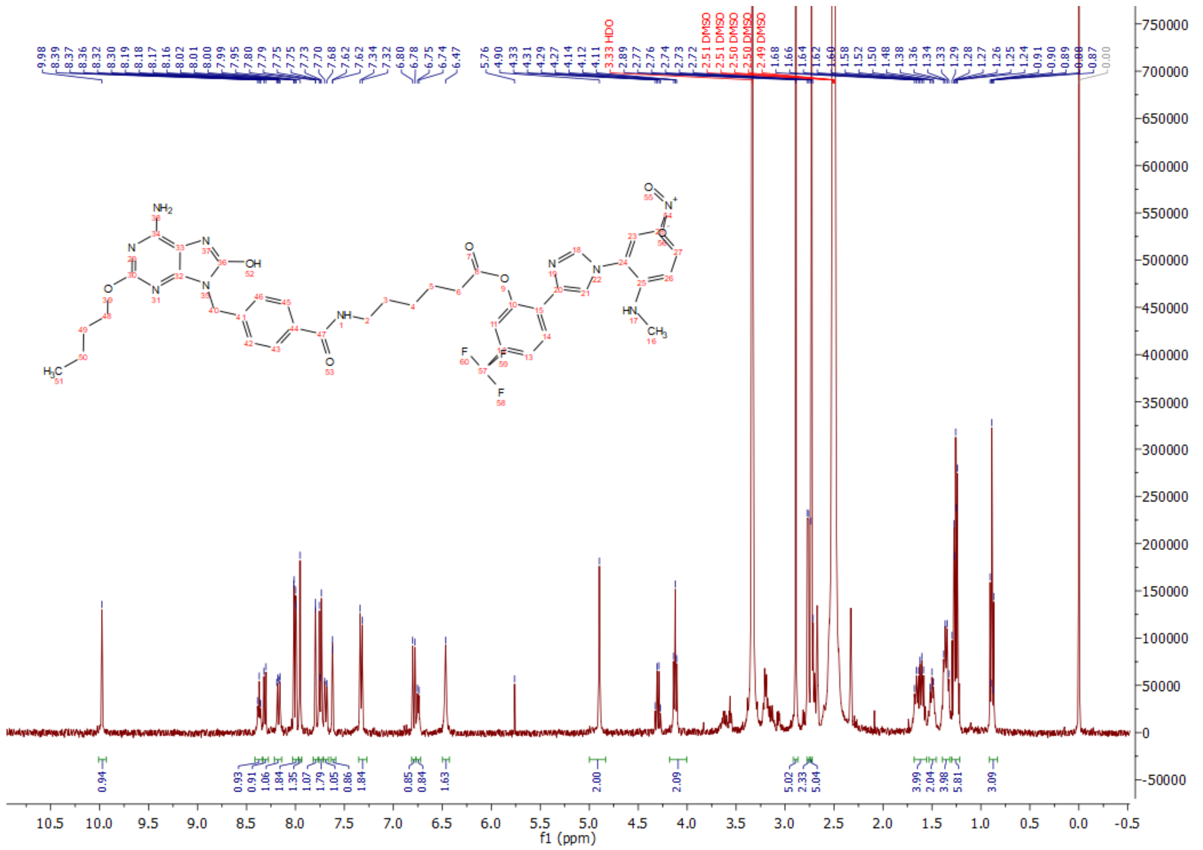

Compound **T1/2/T7:** $^{1}H$, 100 MHz, DMSO-d$_6$

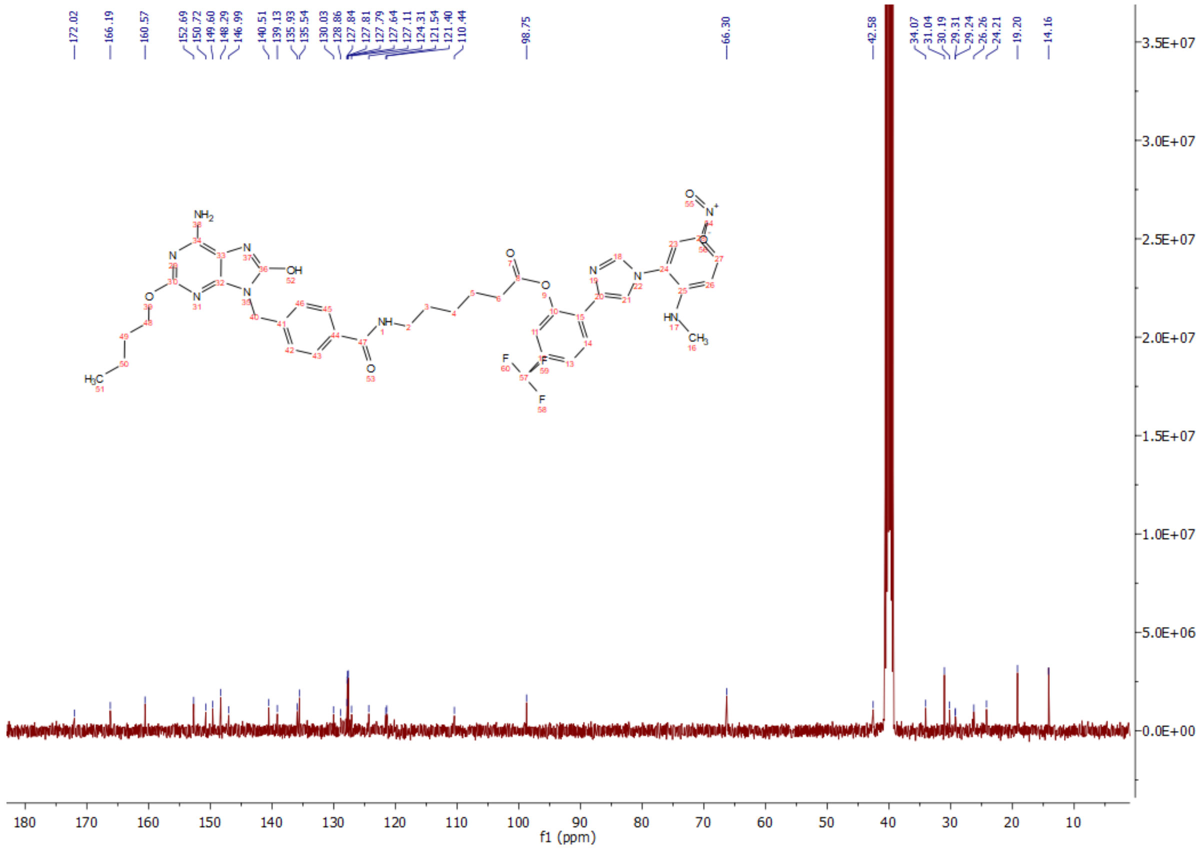

Compound **T7/RI**: $^{1}H$, 400 MHz, DMSO-$d_6$

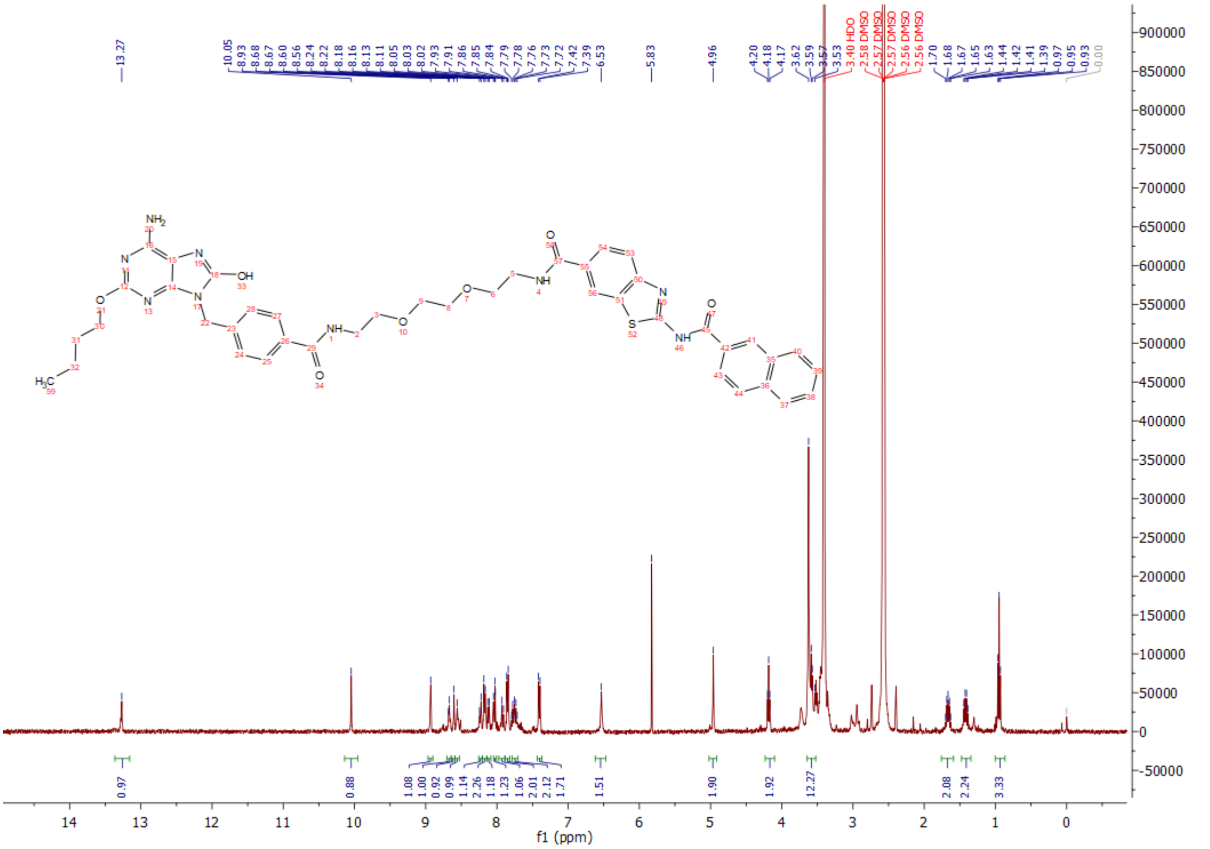

Compound **T7/RI**: $^{13}C$, 100 MHz, DMSO-$d_6$

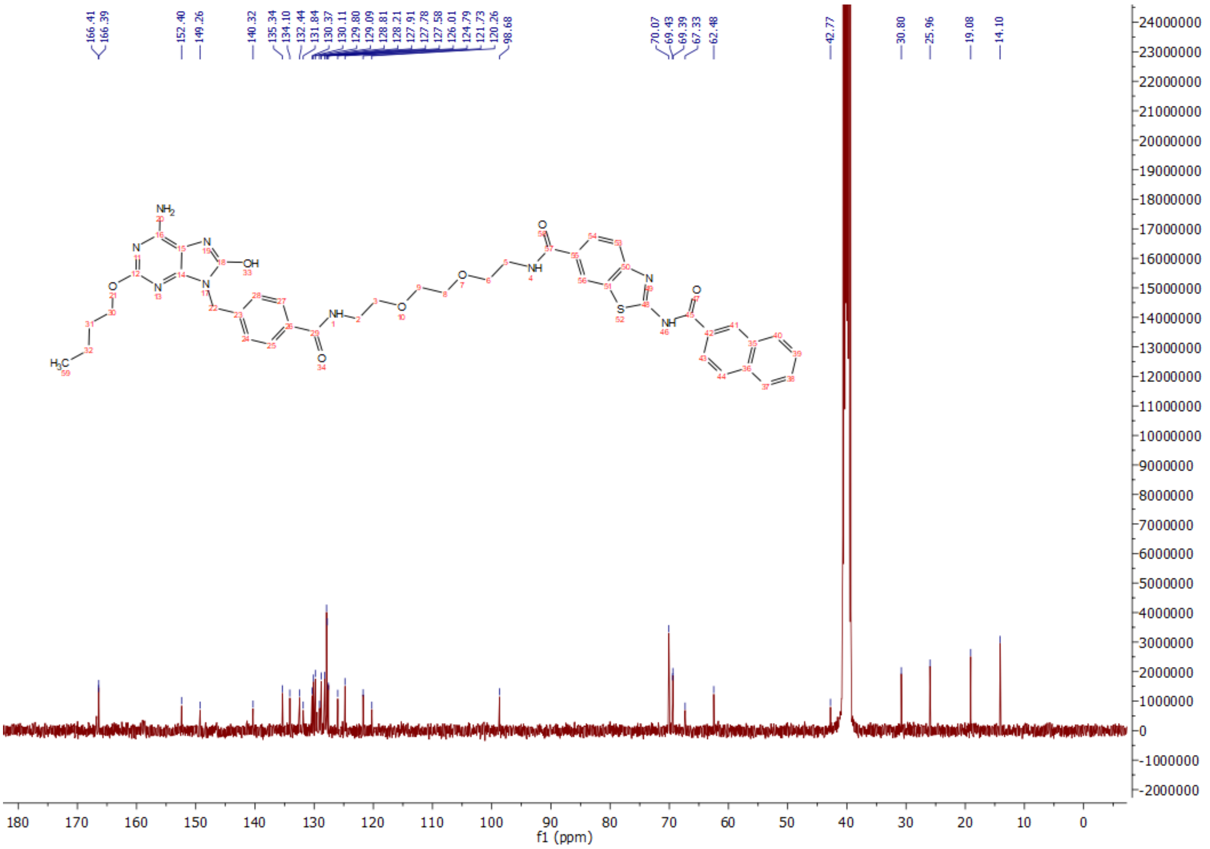

Compound **T4/RI**: $^{1}H$, 400 MHz, DMSO-$d_6$

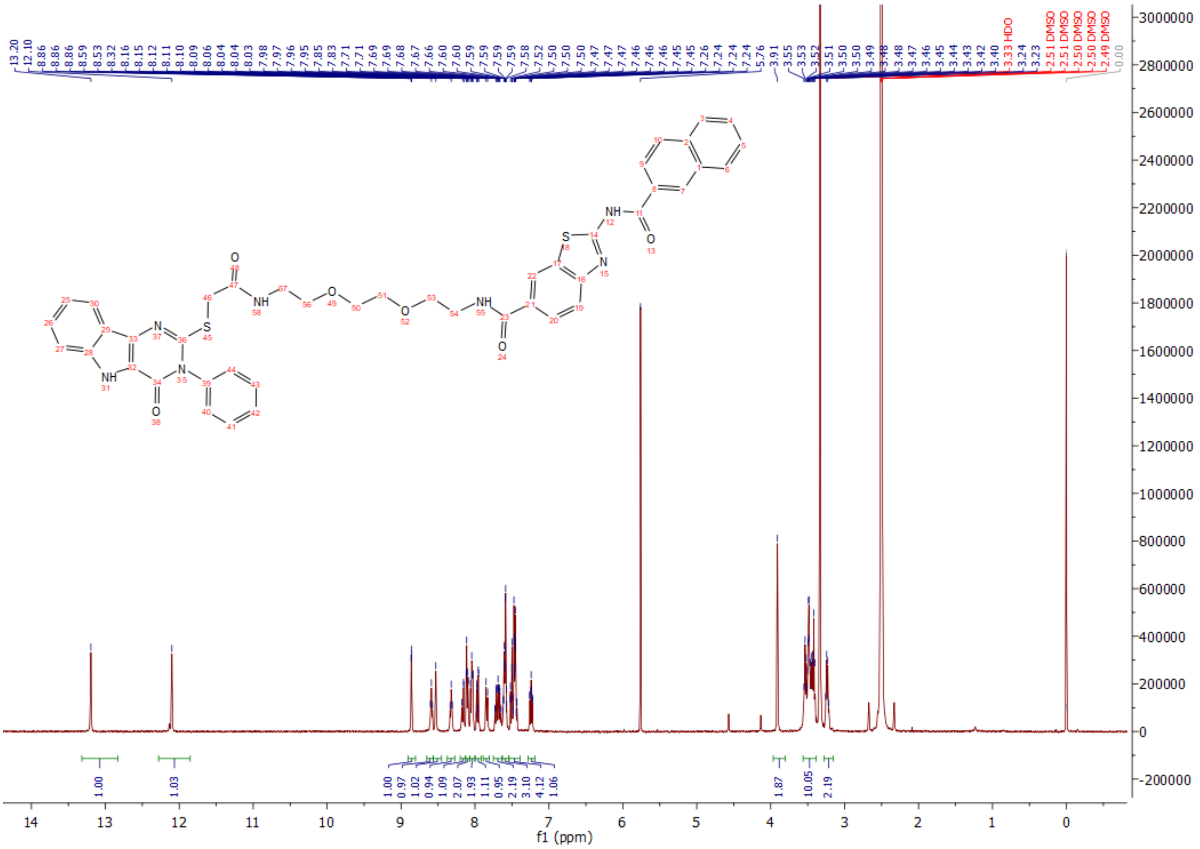

Compound **T4/RI**: $^{13}C$, 100 MHz, DMSO-$d_6$

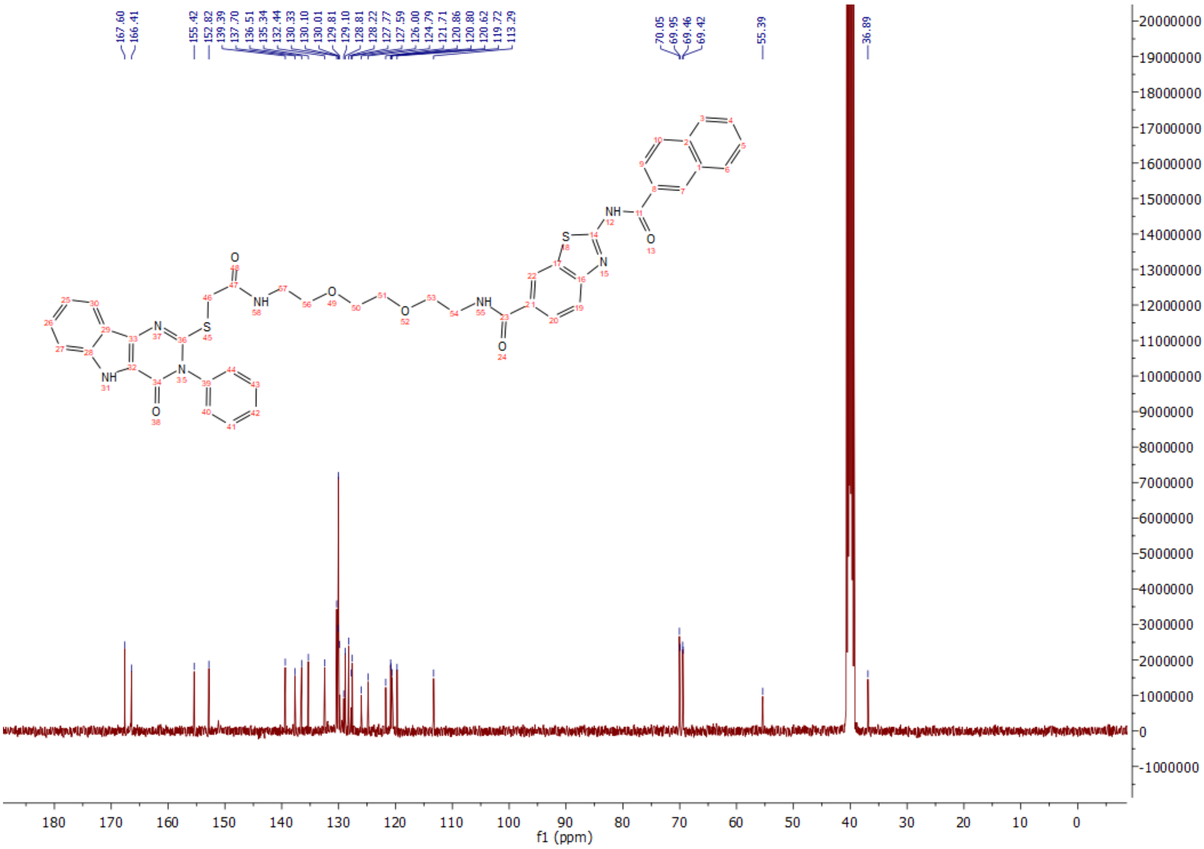

Compound **T1/2/RI**: $^{1}H$, 400 MHz, DMSO-d$_6$

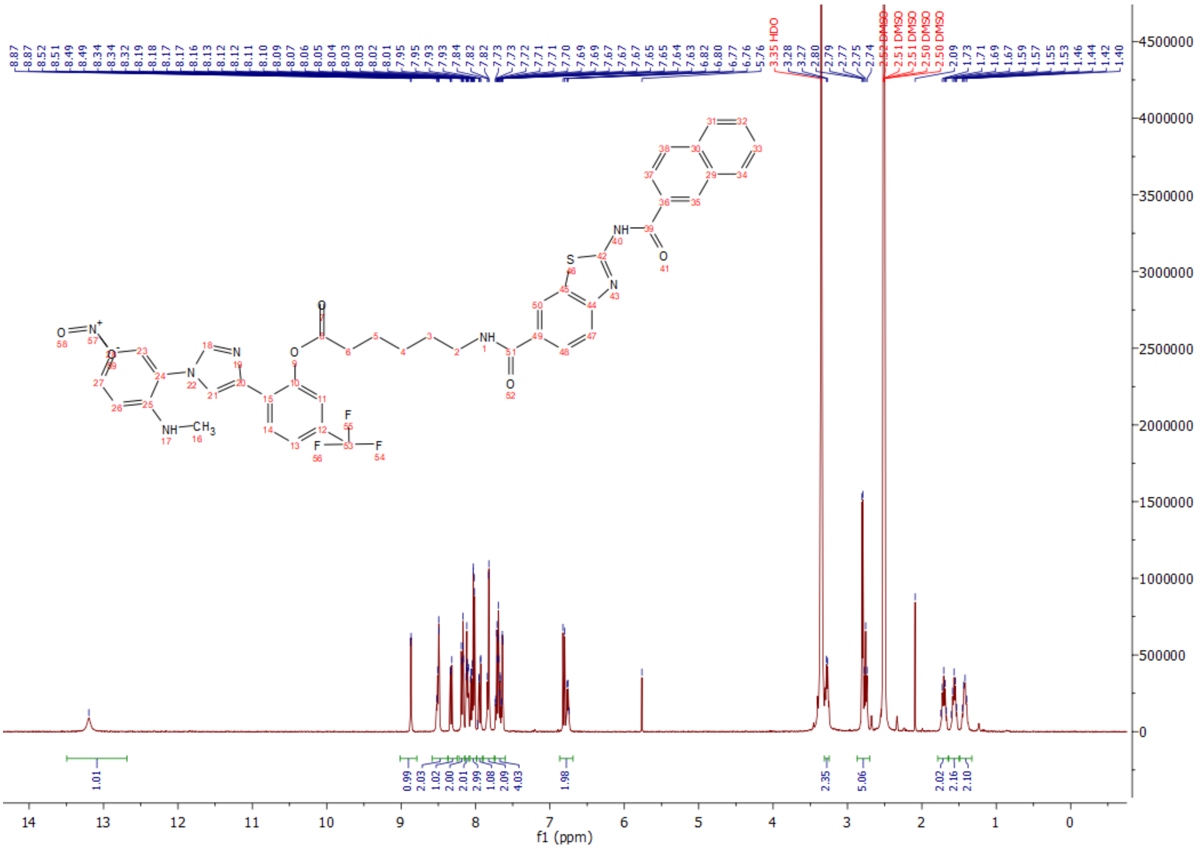

Compound **T1/2/RI**: $^{13}C$, 100 MHz, DMSO-d$_6$

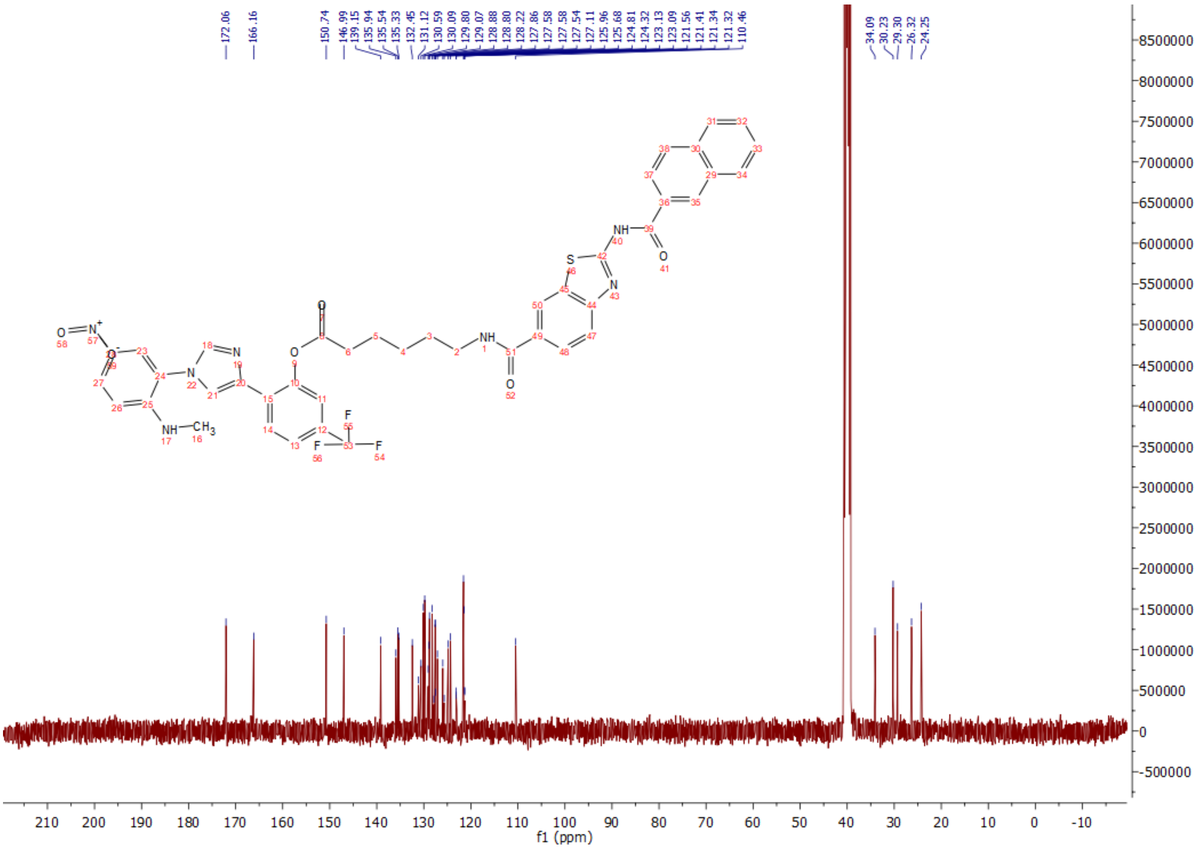

Compound **N2/RI**: $^{1}H$, 400 MHz, DMSO-$d_6$

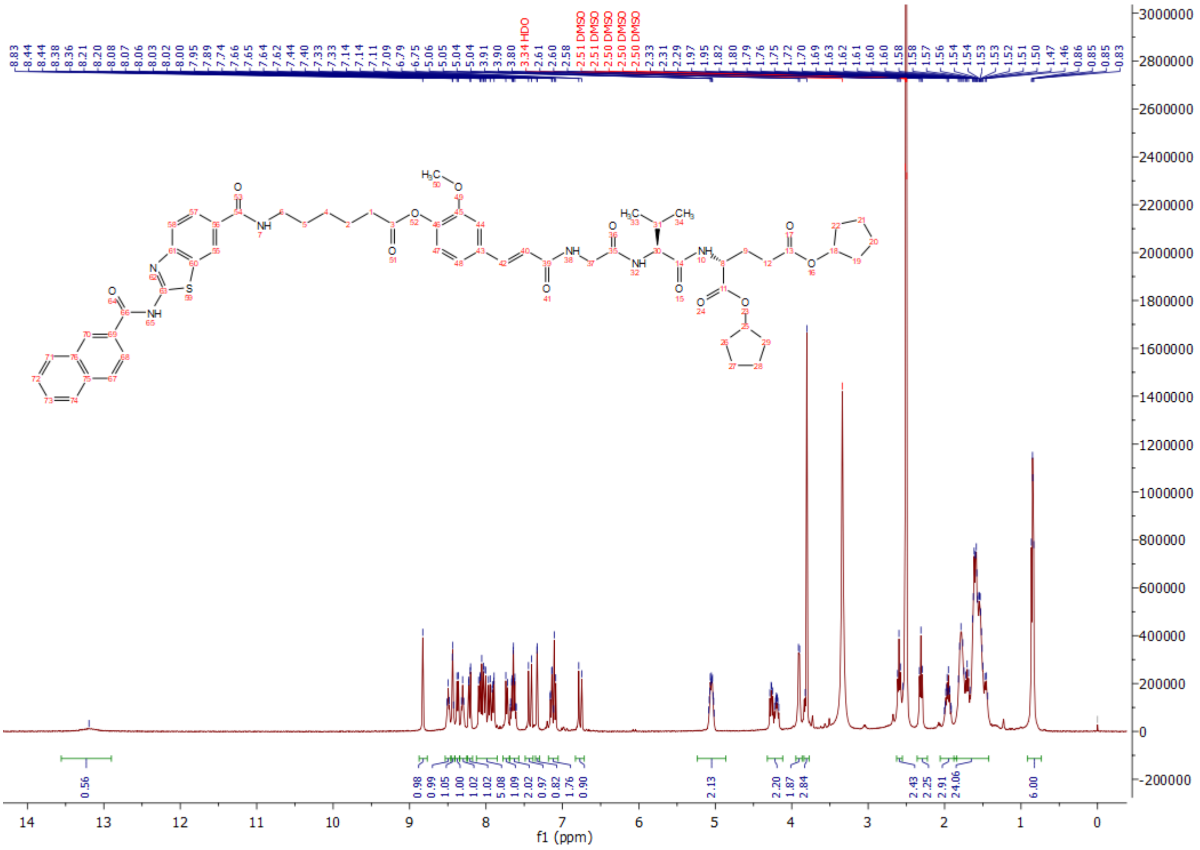

Compound **N2/RI**: $^{13}C$, 100 MHz, DMSO-$d_6$

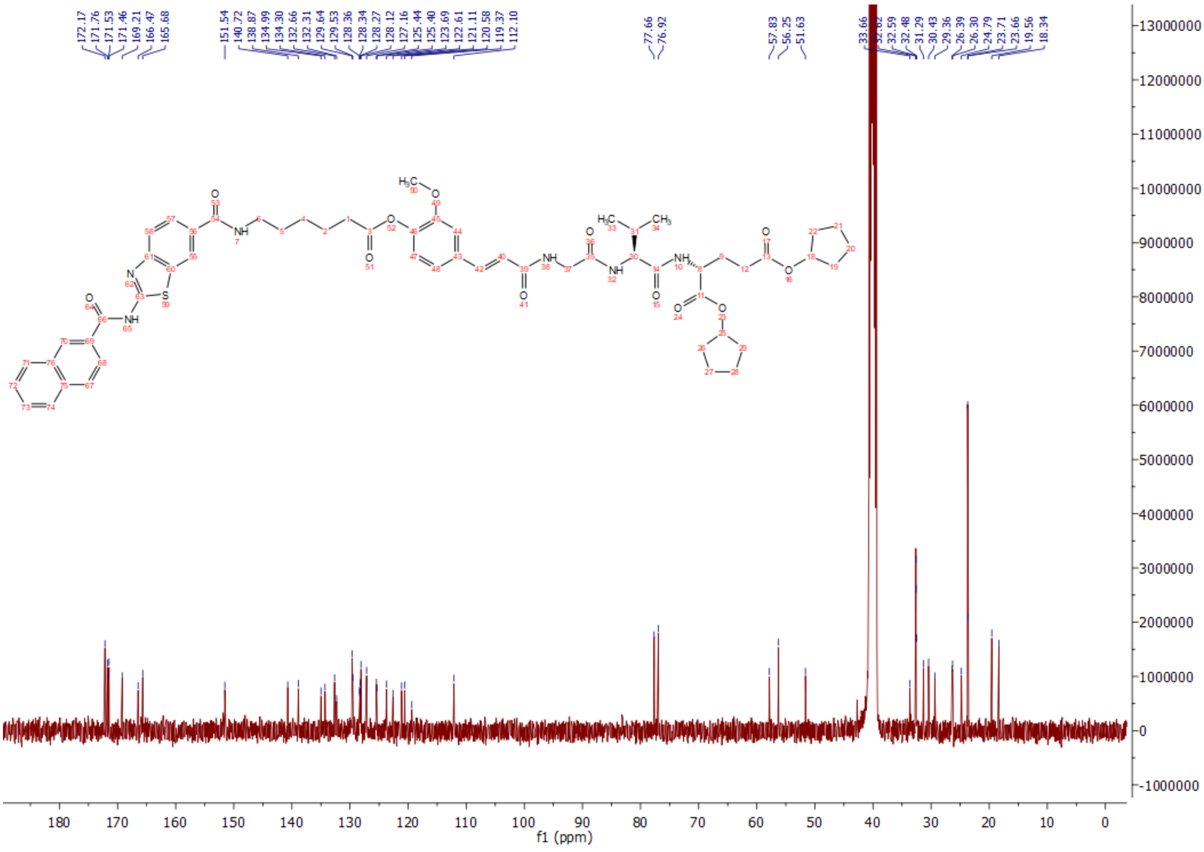

7.) K. Bedard, D.R. Goldberg, P. Probst, The retinoic acid inducible gene "rig-1" pathway and methods of use thereof. WO2020033782.
